# P2RY2 is a purinergic immune checkpoint linking extracellular ATP to immune evasion and adaptive resistance to immunotherapy

**DOI:** 10.1101/2025.10.09.681049

**Authors:** Zhaoqing Hu, Hitoshi Matsuo, Shangce Du, Cecilia Berzain Battioni, Lena Jassowicz, Rafael Carretero, Melanie Sator-Schmitt, Xiyue Zhao, Beiping Miao, Cansu Eris, Helena Engel, Mohamed A.A. Mahmoud, Elke Laport, Yanling Xiao, Ilse Hofmann, Christel Herold-Mende, Chong Sun

## Abstract

Extracellular ATP (eATP) accumulates substantially in the tumor microenvironment (TME) and rises further during immunotherapy. While canonically an immune-activating “danger” signal, eATP also promotes immunosuppression in tumors, thus far largely attributed to its metabolite, adenosine. Here, we identify direct eATP signaling through P2RY2 as a dominant, adenosine-independent mechanism of immune resistance. Specifically, eATP–P2RY2 signaling serves as the primary upstream driver of COX-1/2 upregulation and consequent accumulation of immunosuppressive PGE₂ in the TME, uncovering the long-sought TME-specific trigger of pathological COX–PGE₂ hyperactivation in solid tumors. Genetic deletion or pharmacologic inhibition of P2RY2 eliminates both baseline and therapy-induced intratumoral PGE₂, restores antitumor T cell responses, and reverses resistance to CAR-T, TCR-T, checkpoint blockade, and TIL therapies. Given that persistently elevated eATP is a hallmark of solid tumors, our work reveals a fundamental mechanism by which tumors hijack innate “danger” signaling to establish immune suppression and develop adaptive resistance to immunotherapy. These findings establish P2RY2 as a purinergic immune checkpoint with translational potential for combinatorial cancer immunotherapies.

**HIGHLIGHTS:**

- eATP in the TME drives baseline and therapy-induced immune evasion via P2RY2
- eATP-P2RY2 signaling is the primary cause of PGE₂ accumulation in the TME
- P2RY2 propels a feedback loop that enforces adaptive resistance to immunotherapies
- P2RY2 blockade reprograms the TME and restores immunotherapy responsiveness

## INTRODUCTION

Adenosine triphosphate (ATP), the central energy currency of the cell, is typically confined intracellularly at millimolar concentrations, while extracellular ATP (eATP) is maintained at nanomolar levels in tissues under physiological conditions^1–3^. However, cellular stress and injury, as well as cell death trigger ATP release into the extracellular space as a damage-associated molecular pattern (DAMP) that activates innate immuniy^3,4^. As a result, eATP accumulates markedly in the TME, reaching concentrations over 1,000-fold higher than in healthy tissues^2,4–6^. Notably, cancer therapies, such as chemotherapy and radiation, further elevate eATP levels, through both active secretion and passive release following treatment-induced cellular stress and lysis^7–11^.

eATP exhibits paradoxical immunomodulatory roles^3,12^. Acutely, it acts as a pro-inflammatory “danger” signal that activates innate immune responses and facilitates adaptive immune priming. It promotes dendritic cell maturation and inflammasome-mediated IL-1β secretion via P2Y11 and P2X7 receptors^8,13–15^, drives neutrophil chemotaxis via P2RY2^16^, and directly boosts T cell activation and migration via P2X7^17–20^ and P2X4^21^, respectively.

In contrast, persistently high concentrations of eATP (micromolar–millimolar range) exerts immunosuppressive effects. Prolonged engagement of the P2X7 receptor by eATP induces T cell dysfunction or death^22^ and augments the suppressive activity of myeloid-derived suppressor cells (MDSCs)^23^. Moreover, high eATP levels can also directly trigger death of dendritic cells and macrophages via P2X7^24–26^, as well as thymocytes via P2X1^27^. Furthermore, eATP is rapidly hydrolyzed by ectonucleotidases CD39 and CD73 into adenosine, a potent immunosuppressive metabolite that inhibits effector T and NK cell functions and promotes regulatory T cell activity^28,29^.

In contrast to the extensively studied adenosine pathway, the direct immunological impact of persistently elevated eATP levels in the TME remains poorly defined^3,5,30^. eATP signals through a family of purinergic receptors with diverse and context-dependent functions, yet its net effect on antitumor immunity, particularly during therapy-induced eATP surges, has not been elucidated. Clarifying how sustained eATP signaling influences immunotherapy efficacy represents a critical and underexplored question in tumor immunology.

Prostaglandin E₂ (PGE₂), synthesized by cyclooxygenases COX-1 and COX-2, is prominent immunosuppressive mediator in tumors^31^. Solid tumors frequently exhibit elevated COX-1/2 activity and PGE₂ accumulation, which promote tumor progression and resistance to immunotherapy^32–38^. Excessive PGE₂ shapes an immunosuppressive TME through affecting various types of immune cells: it impairs dendritic cell function^33,38^; drives macrophages polarization toward an M2-like phenotype^39–41^; expands MDSCs^39,42–44^; recruits and activates regulatory T cells^45^; inhibits NK cell activity^46,47^; and suppresses effector T cell expansion and functions^33,35,36,48–50^.

Despite the prevalence and pleiotropic immunosuppressive functions of intratumoral PGE₂ in cancer, the upstream driver of chronically elevated COX-1/2 expression and PGE₂ accumulation in tumors has remained undefined. While many candidates have been implicated based on in vitro data, including oncogenic pathways (e.g. EGFR, KRAS), inflammatory mediators (e.g. IL-1β, TLR ligands), and purinergic receptors (e.g. P2RY2, P2RY6), the primary cause of COX–PGE₂ hyperactivation in the TME remains elusive, with no definitive in vivo evidence identifying a dominant trigger^51–63^.This question gains added importance given our findings that immunotherapy further amplifies intratumoral PGE₂ levels (Figure 3E-H), reinforcing the need to identify upstream drivers of this immunosuppressive axis.

In the absence of a known tumor-specific trigger, current clinical strategies to counter PGE₂-mediated immunosuppression rely on broad COX inhibitors or PGE₂ receptor antagonists. These approaches have yielded mixed clinical outcome across trials^64–68^. Moreover, systematic COX inhibition, due to the physiological importance of prostaglandin signaling in tissue homeostasis, has been associated with significant toxicity^69–77^. These findings underscore the need for a tumor-selective, upstream intervention to disrupt the COX–PGE₂ immunosuppressive axis at its origin.

Our study identifies the eATP–P2RY2 signaling axis as the dominant upstream driver of aberrant COX-1/2 expression and chronic PGE₂ accumulation in solid tumors, revealing a previously unrecognized adenosine-independent mechanism of immune evasion. In addition to driving baseline immunosuppression, therapy-induced surges of eATP in the TME further amplify PGE₂ production via P2RY2, fueling adaptive resistance to immunotherapy.

Notably, eATP–P2RY2 signaling emerges as a prevalent mechanism of immunotherapy resistance across diverse tumor types. In multiple mouse models of pancreatic and colorectal cancer that acquire resistance to immunotherapies, genetic or pharmacologic disruption of P2RY2 eliminated intratumoral PGE₂, reprogrammed the TME to support T cell infiltration, functionality and persistence, and restore responses to CAR-T, TCR-T, and checkpoint blockade therapies.

Furthermore, in ex vivo assays with human tumors, P2RY2 blockade (via a small-molecule inhibitor or a newly developed monoclonal antibody) enhanced autologous TIL-mediated tumor cell killing. Consistently, melanoma patients with high tumoral P2RY2 expression exhibited significantly shorter survival following anti–PD-1 therapy, linking P2RY2 activity to immune resistance in patients.

Together, our findings delineate a tumor-selective, targetable immunosuppressive circuit whereby a canonical innate “danger” signal (eATP) is co-opted via P2RY2 to drive both baseline and therapy-induced adaptive immune evasion. These results establish P2RY2 as a promising therapeutic target for enhancing cancer immunotherapies.

## RESULTS

### eATP, independent of its metabolites, suppresses T cell function via tumor-intrinsic purinergic receptor P2RY2

Elevated levels of eATP in the TME are a hallmark of solid tumors^5,12^. In four humanized and syngeneic in vivo tumor models, we observed that eATP concentrations in tumor interstitial fluid ranged from 32 to 140 µM, exceeding levels in normal tissues (0.01–0.06 µM) and peripheral blood (0.02–0.1 µM) by over a thousand-fold (Figure 1A-C and S1A). Notably, intratumoral eATP levels further increased upon treatment of tumor-bearing mice with TCR-T, CAR-T, or immune checkpoint blockade (ICB) therapies (Figure 1A-C and S1A). Despite these immunotherapy interventions, tumors exhibited only modest responses and eventually developed resistance (Figure 4G-J, R and S).

**Figure. 1.**
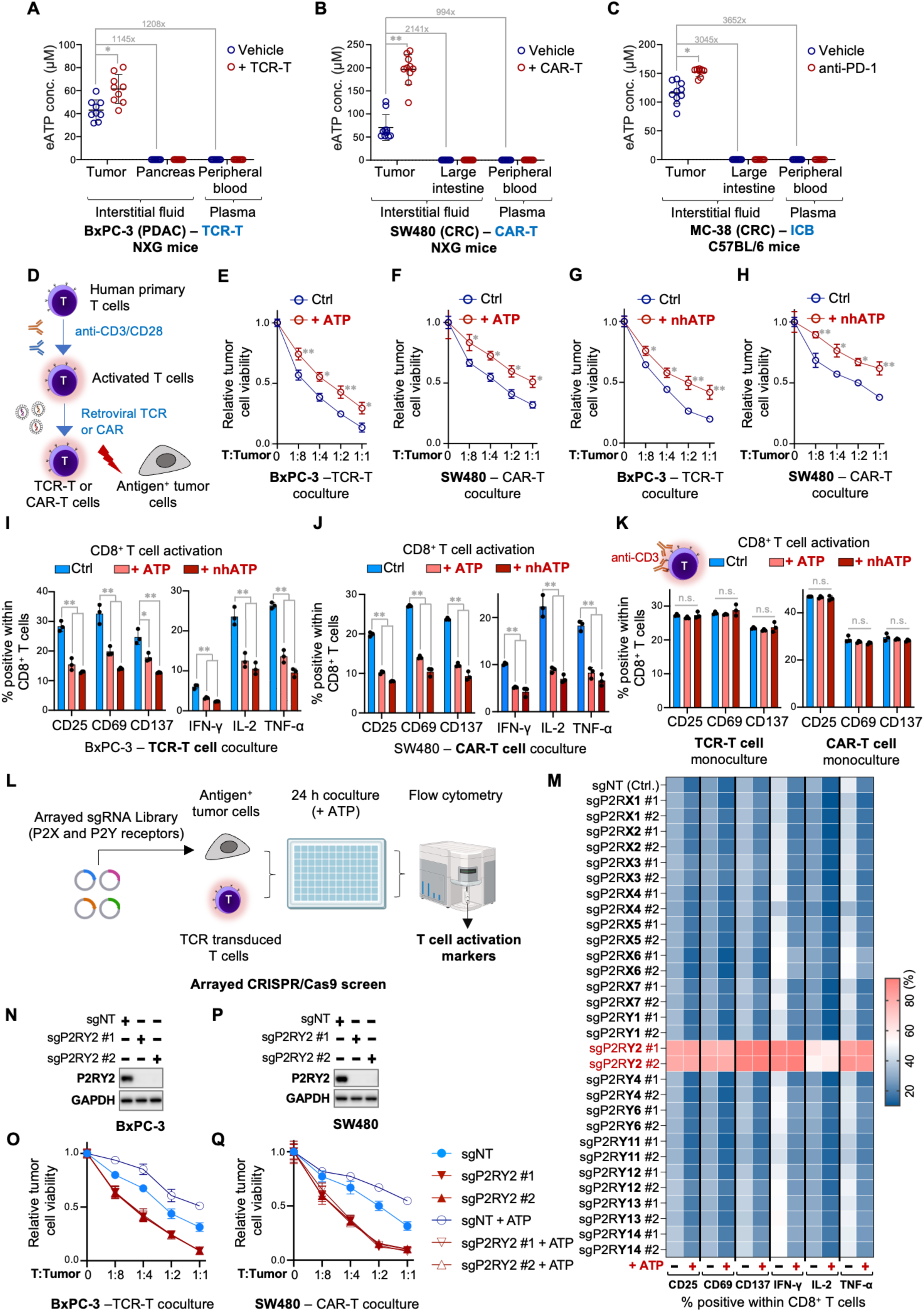
eATP, independent of its metabolites, suppresses T cell function via tumor-intrinsic purinergic receptor P2RY2. (**A**–**C**) Extracellular ATP (eATP) concentrations are markedly elevated in the TME compared to corresponding normal tissues and further increase upon immunotherapy. Interstitial fluid from tumors and matched normal tissues (large intestine for SW480 and MC-38 tumors; pancreas for BxPC-3 tumors), as well as peripheral blood plasma, were collected from both immunotherapy-treated and vehicle control groups. eATP levels in the interstitial fluid and plasma were measured using the CellTiter-Glo assay. Fold changes are indicated. (A) Human TCR-T cell therapy against human PDAC BxPC-3 tumors in vivo. Human T cells from a healthy donor were activated and transduced with a MART-1-specific TCR (clone 1D3) targeting the HLA-A2-restricted MART-1 epitope. To enable TCR-T cell recognition, BxPC-3 cells were transduced with a fusion construct encoding the MART-1 epitope (ELAGIGILTV), HLA-A2, and β2-microglobulin (B2M) to ensure stable antigen presentation. The resulting tumor cells were implanted into NXG immunodeficient mice, followed by treatment with either vehicle control (Vehicle) or MART-1 TCR-T cells (+TCR-T). **(B)** Human CAR-T cell therapy against human CRC SW480 tumors in vivo. Human T cells from a healthy donor were activated and transduced with a CEA-specific CAR (clone MFE-23). SW480 cells, which constitutively express CEA, are recognized by the CEA CAR-T cells. After tumor implantation in NXG mice, treatment with either vehicle control (Vehicle) or CEA CAR-T cells (+CAR-T) was administered. (C) ICB therapy against syngeneic mouse CRC MC-38 tumors in vivo. Murine CRC MC-38 cells were subcutaneously implanted into immunocompetent C57BL/6 mice. After tumor establishment, mice were treated with either vehicle control (Vehicle) or PD-1 blocking antibody (anti–PD-1). **(D)** Schematic illustration of the antigen-specific T cell–tumor cell coculture system. Primary human T cells were activated with anti-CD3/CD28 and retrovirally transduced with a TCR or CAR targeting tumor cells that were either loaded with or naturally expressed a matched antigen. For all coculture assays and humanized in vivo models presented in this manuscript, CAR-T and TCR-T cells were generated following this scheme. The specific CAR, TCR, and antigens used are detailed in the individual experiments. (**E**, **F**) Supplementation with exogenous ATP enhances tumor cell survival in the presence of tumor-reactive TCR-T and CAR-T cells. As illustrated in Figure 1D, MART-1 TCR–transduced T cells were cocultured with MART-1 epitope–loaded BxPC-3 cells (E), and CEA CAR–transduced T cells were cocultured with SW480 tumor cells that constitutively express CEA (F). Cocultures were conducted in the absence or presence of 200 μM ATP at specified effector-to-target (T:Tumor) ratios. After 72 hours, tumor cell viability was assessed using the CellTiter-Blue Cell Viability assay. (**G, H**) Supplementation with non-hydrolyzable ATP analog (nhATP) enhances tumor cell survival following coculture with tumor-reactive TCR-T and CAR-T cells. The same experimental setup as described in panels (E, F) was used, except that the supplemented ATP was replaced with 200 μM of the non-hydrolyzable analog ATPγS. (**I**, **J**) eATP suppresses T cell activation in coculture with tumor cells. MART-1 TCR–transduced T cells (I) or CEA CAR–transduced T cells (J) were cocultured with antigen-positive BxPC-3 or SW480 cells, respectively, for 24 hours in the absence or presence of 200 μM ATP or the non-hydrolyzable analog ATPγS (nhATP). CD8⁺ T cell activation was assessed by flow cytometry, measuring the expression of activation markers (CD25, CD69, CD137) and cytokine production (IFN-γ, IL-2, TNF-α). **(K)** ATP does not affect T cell activation in the absence of tumor cells. TCR-T and CAR-T cells, described in panels (E–J), were stimulated with 0.1 μg/mL anti-CD3 in the absence or presence of 200 μM ATP or the non-hydrolyzable analog ATPγS (nhATP), followed by flow cytometry analysis of CD25, CD69, and CD137 expression. **(L)** Schematic illustration of a purinergic receptor–centered, arrayed CRISPR screen to identify mediators of eATP-induced T cell suppression. Cas9-expressing, MART-1 epitope–loaded BxPC-3 cells were individually transduced with sgRNAs targeting each P2X and P2Y purinergic receptor and cocultured with MART-1 TCR– transduced T cells in the absence or presence of 200 μM supplemented ATP. T cell activation markers and effector cytokines were assessed individually after 24 hours by flow cytometry. **(M)** Heatmap summarizing T cell activation across cocultures with tumor cells harboring knockout of individual P2X and P2Y receptors. Among all receptors tested, only P2RY2 knockout (highlighted in red) substantially reversed ATP-mediated suppression of T cell activation, restoring expression of CD25, CD69, CD137, IFN-γ, IL-2, and TNF-α in CD8⁺ T cells. (**N, P**) Efficient CRISPR/Cas9-mediated knockout of P2RY2 in tumor cells confirmed by immunoblotting. Western blot analysis of P2RY2 expression in Cas9-expressing BxPC-3 (N) and SW480 (P) cells transduced with sgRNAs targeting P2RY2 (sgP2RY2 #1 and sgP2RY2 #2) or a non-targeting control sgRNA (sgNT). GAPDH served as a loading control. (**O**, **Q**) P2RY2 knockout in tumor cells abolishes eATP-mediated protective effect against tumor-reactive T cells. P2RY2-deficient (sgP2RY2 #1, #2) and P2RY2-proficient (sgNT) BxPC-3 (O) and SW480 (Q) tumor cells were cocultured with tumor-reactive TCR-T or CAR-T cells at indicated effector-to-target (T:Tumor) ratios for 72 hours, with or without ATP supplementation. Tumor cell viability was assessed using the CellTiter-Blue Cell Viability Assay. Data represent the mean ± standard deviation of biological replicates (n ≥ 3). For panels (A), the sample sizes are: vehicle (n=9), + TCR-T (n=9). For panels (B), the sample sizes are: vehicle (n=10), + CAR-T (n=10). For panels (C), the sample sizes are: vehicle (n=10), anti-PD-1 (n=8). P values were determined using unpaired two-tailed Student’s t-test (A-C), two-way ANOVA with Šídák’s multiple comparisons test (E-H) or one-way ANOVA with Dunnett’s multiple comparisons test (I-K). A p-value ≥ 0.05 indicates non-significance (n.s.), while a p-value < 0.05 is denoted as ∗, and a p-value < 0.0001 is represented as ∗∗. See also Figure S1 and S2.

To assess the impact of eATP on antitumor T cell function, we established antigen-specific coculture systems using primary human TCR-T and CAR-T cells and tumor cells constitutively expressing or loaded with the matched antigens (Figure 1D). Both CAR- and TCR-T cells exhibited robust, dose-dependent cytotoxicity against antigen-positive tumor cells (Figure S1B and C). Supplementation with exogenous eATP increased tumor cell survival in cocultures of PDAC (BxPC-3) and CRC (SW480) cells with TCR-T or CAR-T cells (Figure 1E and F). This protective effect was recapitulated in cocultures involving tumor cells from additional cancer types, including breast, melanoma, lung, prostate, and liver cancers (Figure S1D).

Since eATP can be hydrolyzed into AMP, ADP, and adenosine by cell-surface ectonucleotidases ^78^, we next tested whether the observed protective effects were directly attributable to eATP itself. Supplementation with the non-hydrolyzable ATP (nhATP) analog ATPγS similarly enhanced tumor cell survival in cocultures with antigen-matched TCR-T and CAR-T cells, confirming direct eATP involvement (Figure 1G and H). Importantly, neither ATP nor ATPγS affected tumor cell proliferation in monoculture (Figure S1E and F). Parallel experiments using antigen-specific mouse T cells and CRC cells (Figure S1G) yielded comparable results: ATP and ATPγS do not influence tumor cell proliferation (Figure S1H and I) but do enhance tumor cell survival in the presence of tumor-reactive T cells (Figure S1J and K). These findings suggest that eATP itself protects tumor cells from tumor-reactive T cells.

Consistent with these protective effects, the addition of exogenous eATP or ATPγS to the coculture potently suppressed T cell activation, as evidenced by reduced expression of activation markers (CD25, CD137, CD69) and decreased production of effector cytokines (IL-2, IFN-γ, TNF-α) compared to untreated controls. These suppressive effects were observed in both CD8^+^ (Figure 1I and J; Figure S2A and B) and CD4^+^ (CD3^+^CD8^−^) T cell populations (Figure S2C and D).

Notably, eATP supplementation had no effect on T cell activation when T cells were stimulated with agonist anti-CD3 antibodies in the absence of tumor cells (Figure 1K; Figure S2E-G), suggesting that eATP-mediated T cell suppression operates through tumor cells rather than directly on T cells.

eATP primarily signals through a family of plasma membrane receptors known as P2 purinergic receptors, which include ATP-gated cation channels (P2X receptors) and G protein–coupled receptors (P2Y receptors). To identify the receptor(s) in tumor cells responsible for ATP-mediated T cell suppression, we generated an arrayed CRISPR knockout sgRNA library targeting P2 receptors and individually introduced them into BxPC3 tumor cells. These modified tumor cells were cocultured with tumor-reactive T cells in the presence or absence of eATP, and T cell activation was measured (Figure 1L). eATP supplementation in coculture suppressed T cell activity across most tumor cell variants (Figure 1M). Strikingly, among the receptors tested, P2RY2 emerged as the only one whose deletion markedly enhanced T cell activation, both in the absence and presence of supplemented eATP, and fully eliminated the impact of eATP supplementation (Figure 1M).

Consistent with this, eATP supplementation increased tumor cell survival in the presence of TCR-T or CAR-T cells, whereas disruption of P2RY in tumor cells completely abolished this protective effect (Figure 1N-Q). Notably, even without supplemented eATP, P2RY2-deficiency in tumor cells lifted T cell suppression (Figure 1M) and increased tumor cell viability when cocultured with tumor-reactive T cells (Figure 1N-Q), in line with the accumulation of eATP up to micromolar levels following antigen-specific T cell – tumor cell interactions (Figure S2H and I).

Together, the findings that (i) nhATP mimics the effect of native ATP and (ii) deletion of the cognate eATP receptor P2RY2 abolishes the impact of eATP supplementation collectively demonstrate that eATP suppresses antitumor T cell responses through tumor-intrinsic P2RY2, independently of its hydrolyzed metabolites.

### Tumor-intrinsic P2RY2 suppresses antitumor T cell responses across diverse cancer types

Having identified a potent role for tumor-expressed P2RY2 in suppressing T cell activity, we next sought to evaluate the generality of this function across a broader panel of tumor models. We tested the effect of P2RY2 knockout in 12 additional human tumor cell lines (together with BxPC3 and SW480, comprising a total of 14 lines) spanning colorectal (CRC), pancreatic (PDAC), melanoma, breast, prostate, hepatocellular (HCC), and non–small cell lung cancer (NSCLC) origins (Figure S3A). To establish antigen-specific T cell–tumor cell interactions, we employed two systems: (1) MART-1–specific TCR-T cells cocultured with tumor cells engineered to present the MART-1 peptide on matched MHC class I, and (2) carcinoembryonic antigen (CEA)–specific CAR-T cells cocultured with tumor cells constitutively expressing CEA. In all tested lines, P2RY2 knockout had no impact on intrinsic tumor cell proliferation in the absence of T cells (Figure S3B). Remarkably, upon exposure to tumor-reactive T cells, P2RY2-deficient tumor cells exhibited significantly reduced viability compared to the P2RY2-proficient counterparts across all cancer types (Figure 2A). This reduced tumor cell survival in the cocultures can be explained by markedly enhanced activation and effector function in both CD8⁺ T cells (Figure 2B and C) and CD8⁻ T cell subsets (Figure S3C).

**Figure. 2.**
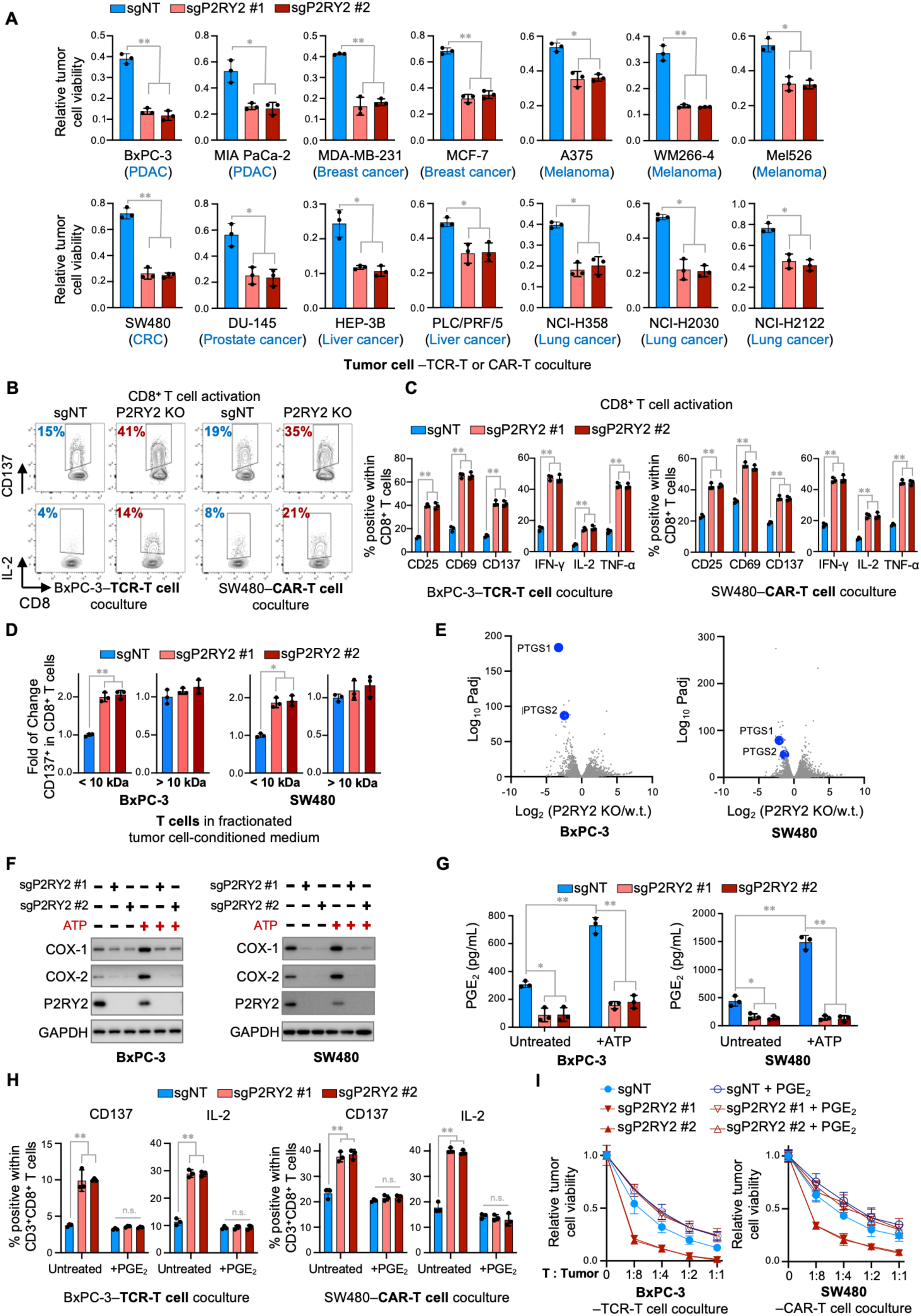
P2RY2 is a predominant driver of the COX-1/2–PGE₂ axis across diverse tumor types, inhibiting antitumor T cell responses. **(A)** P2RY2 knockout in tumor cells leads to enhanced tumor killing by tumor-reactive T cells across multiple cancer types. P2RY2-deficient (sgP2RY2 #1, sgP2RY2 #2) and P2RY2-proficient (sgNT) tumor cells from various cancer types were cocultured with T cells transduced with either a CEA-specific CAR or a MART-1 TCR (clone 1D3, CD8-independent) that targets the HLA-A2-restricted MART-1 epitope, as illustrated in Figure 1D. P2RY2-proficient (sgNT) cells served as controls. Tumor cell viability was assessed using the CellTiter-Blue Cell Viability Assay. To enable antigen-specific T cell–tumor cell interactions, different strategies were applied depending on the endogenous expression of CEA, MART-1, and HLA-A2 in each tumor cell line: CEA-expressing tumor cells (SW480) were directly cocultured with CEA CAR-T cells. MART-1 and HLA-A2 double-positive tumor cells (WM266-4, Mel526) were directly cocultured with MART-1 TCR-T cells. HLA-A2⁺ but MART-1⁻ tumor cells (MDA-MB-231, MCF-7, A375) were preloaded with the MART-1 epitope by peptide pulsing or lentiviral transduction before coculture with MART-1 TCR-T cells. Tumor cells lacking both MART-1 and HLA-A2 (BxPC-3, MIA PaCa-2, DU-145, HEP-3B, PLC/PRF/5, NCI-H358, NCI-H2030, and NCI-H2122) were transduced with a fusion construct encoding MART-1 epitope (ELAGIGILTV), HLA-A2, and β2-microglobulin (B2M) before coculture. Tumor cell viability was assessed using the CellTiter-Blue Cell Viability Assay after 72 hours of coculture with antigen-specific CAR-T or TCR-T cells. **(B, C)** P2RY2 knockout in tumor cells enhances T cell activation in coculture with tumor-reactive TCR-T and CAR-T cells. P2RY2-proficient (sgNT) or P2RY2-deficient (sgP2RY2 #1, sgP2RY2 #2) BxPC-3 and SW480 tumor cells were cocultured with TCR-T cells or CAR-T cells, as described in Figure 2A, for 24 hours, followed by flow cytometry analysis of T cell activation markers. Representative contour plots are shown, with numbers indicating the percentage of CD8⁺ T cells positive for CD137 and IL-2 (B). Bar graphs quantify CD137, CD69, and CD25 activation markers and cytokine production (IFN-γ, IL-2, and TNF-α) in CD8⁺ T cells (C). **(D)** The <10 kDa fraction of conditioned medium from tumor cells is responsible for P2RY2-mediated T cell suppression. Conditioned medium was collected from P2RY2-proficient (sgNT) and P2RY2-deficient (sgP2RY2 #1, sgP2RY2 #2) BxPC-3 and SW480 tumor cells and fractionated into <10 kDa and >10 kDa molecular weight components using Amicon® Ultra Centrifugal Filters. T cells were incubated in the fractionated conditioned medium in the presence of anti-CD3 (0.1 μg/mL) for 24 hours, and CD137 expression in CD8⁺ T cells was assessed by flow cytometry as a readout of T cell activation. The fold change in the percentage of CD137⁺ T cells between P2RY2-proficient and P2RY2-deficient conditions is shown. **(E)** P2RY2 knockout significantly downregulates PGE₂ biosynthesis genes. Volcano plots display differentially expressed genes in P2RY2-proficient (w.t.) and P2RY2-deficient (KO) BxPC-3 (left) and SW480 (right) tumor cells. The x-axis shows the change in gene expression between P2RY2-deficient and wild-type cells on a logarithmic (base 2) scale. The y-axis represents the statistical significance (–Log₁₀ adjusted p-value). *PTGS1* (COX-1) and *PTGS2* (COX-2) are highlighted. **(F)** P2RY2 is a predominant driver of COX-1 and COX-2 expression. Western blot analysis of COX-1, COX-2, and P2RY2 expression in P2RY2-proficient (sgNT) and P2RY2-deficient (sgP2RY2 #1, sgP2RY2 #2) BxPC-3 (left) and SW480 (right) tumor cells, with or without treatment by 200 μM exogenously supplemented ATP for 24 hours. GAPDH served as a control. **(G)** P2RY2 knockout diminishes PGE₂ production. Culture medium conditioned by P2RY2-proficient (sgNT) and P2RY2-deficient (sgP2RY2 #1, sgP2RY2 #2) BxPC3 (left) and SW480 (right) tumor cells, either in the absence (Untreated) or presence (+ATP) of 200 μM exogenously supplemented ATP for 24 hours, was collected. PGE₂ concentrations in the conditioned medium were quantified using ELISA. **(H)** Exogenous PGE₂ abolishes the enhanced T cell activation induced by P2RY2 knockout. P2RY2-proficient (sgNT) or P2RY2-deficient (sgP2RY2 #1, sgP2RY2 #2) BxPC-3 and SW480 tumor cells were cocultured with tumor-reactive TCR-T cells or CAR-T cells, for 24 hours. Flow cytometry analysis was performed to assess T cell activation, measuring CD137 and IL-2 in CD8^+^ T cells. **(I)** Exogenous PGE₂ rescues P2RY2-deficient cells from tumor-reactive T cells. P2RY2-proficient (sgNT) and P2RY2-deficient (sgP2RY2 #1, sgP2RY2 #2) BxPC-3 or SW480 tumor cells were cocultured with tumor-reactive TCR-T or CAR-T cells for 72 hours, with or without PGE₂ supplementation. Tumor cell viability was measured using the CellTiter-Blue Cell Viability Assay. Data represent the mean ± standard deviation of biological replicates (n ≥ 3). P values were determined using Two-way ANOVA with Tukey’s multiple comparisons test (G) or one-way ANOVA with Dunnett’s multiple-comparison test (A, C, D, H). A p-value ≥ 0.05 indicates non-significance (n.s.), while a p-value < 0.05 is denoted as ∗, and a p-value < 0.0001 is represented as ∗∗. See also Figure S3 and S4.

Conversely, overexpression of P2RY2 in tumor cells suppressed T cell activation (Figure S3D and E) and conferred protection against T cell–mediated killing, again without affecting tumor cell proliferation (Figure S3F). These findings establish that tumor-intrinsic P2RY2 acts as a conserved suppressor of antitumor T cell responses across diverse cancer types.

### The eATP-P2RY2 axis dominates COX-1/2 expression across cancer types to drive PGE₂-mediated suppression of antitumor T cell responses

To understand the mechanism by which tumor-intrinsic P2RY2 mediates T cell suppression, we first assessed whether direct contact between T cells and tumor cells was required. Primary human T cells were cultured in conditioned medium (CM) collected from P2RY2-proficient or P2RY2-deficient SW480 and BxPC3 tumor cells. Because these T cells were not exposed to tumor antigens, their baseline activation was induced by agonist anti-CD3 antibodies. Notably, T cells cultured in CM from P2RY2-deficient tumor cells showed significantly higher levels of activation compared to those exposed to CM from P2RY2-proficient cells (Figure S4A and B), suggesting that soluble factor(s) mediate P2RY2-driven T cell suppression.

To identify these factors, we fractionated serum-free CM from tumor cells using a 10 kDa molecular weight cutoff filter to separate small molecules (<10 kDa, primarily metabolites) from larger ones (>10 kDa, mainly proteins). Relative T cell activation induced by anti-CD3 stimulation was then assessed in the presence of each fraction. The <10 kDa fraction from P2RY2-proficient cells significantly suppressed T cell activation compared to that from P2RY2-deficient cells, while the >10 kDa fractions showed minimal differences (Figure 2D). These findings indicate that small, secreted molecule(s) are the primary mediators of P2RY2-dependent T cell suppression.

In parallel, we performed differential gene expression analysis on eATP-stimulated BxPC3 and SW480 cells with or without P2RY2 expression. Notably, PTGS1 and PTGS2, the genes encoding COX-1 and COX-2—the rate-limiting enzymes for biosynthesis of a secreted small molecule - prostaglandin E₂ (PGE₂), were among the most significantly downregulated transcripts in P2RY2-deficient cells from both tumor lines (Figure 2E).

Under baseline conditions, relatively low levels of eATP were detected in tumor cell cultures (Figure S4E), consistent with the notion that ATP can be released both actively and passively by tumor cells ^79^. In this setting, quantitative PCR analysis showed that P2RY2 knockout markedly reduced PTGS1/2 expression, whereas ectopic overexpression of P2RY2 elevated their mRNA levels (Figure S4C and D). Conversely, supplementation with exogenous eATP further induced PTGS1/2 expression in P2RY2-proficient cells, while P2RY2 deficiency completely abolished this induction (Figure S4C and D), underscoring the critical role of P2RY2 in driving COX gene (PTGS1/2) expression.

At the protein level, eATP stimulation upregulated both COX-1 and COX-2 in P2RY2-proficient tumor cells, whereas, notably, P2RY2 disruption substantially reduced COX-1 and COX-2 expression under both baseline and stimulated conditions (Figure 2F). This effect was consistent across tumor cell lines from diverse tissue origins, including colorectal, pancreatic, breast, melanoma, prostate, liver, and lung cancers (Figure 2F and Figure S4F). Correspondingly, ELISA analysis showed that eATP supplementation increased PGE₂ levels in culture supernatants from P2RY2-proficient tumor cells, while P2RY2 knockout markedly reduced PGE₂ accumulation at baseline and following eATP stimulation (Figure 2G). Conversely, P2RY2 overexpression strongly elevated PGE₂ levels (Figure S4G).

Functionally, exogenous PGE₂ addition abrogated the enhanced activation of TCR-T and CAR-T cells cocultured with antigen-matched P2RY2-deficient tumor cells (Figure 2H). Consistently, in coculture with tumor-reactive TCR-T or CAR-T cells, PGE₂ supplementation restored the survival of P2RY2-deficient tumor cells to levels comparable to those of P2RY2-proficient cells (Figure 2I).

These results establish that tumor-intrinsic P2RY2 strongly sustains COX-1/2 expression to drive PGE₂ production, thereby impairing antitumor T cell responses across cancer types.

### The eATP–P2RY2 axis is the primary driver of intratumoral PGE2 accumulation in vivo

Elevated PGE₂ levels are commonly observed across a wide range of solid tumors and are strongly associated with resistance to T cell–based immunotherapies^31,33–38,80^. Although COX-1/2-mediated PGE2 synthesis is well characterized, the upstream trigger responsible for hyperactivation of this pathway in the TME has remained undefined. Given that P2RY2 strongly drives COX-1/2 expression and PGE2 production across diverse tumor types (Figure 2F and G; Figure S4F), and that its ligand, eATP, is highly elevated in the TME (Figure 1A-C; Figure S1A) in in vivo tumor models, we hypothesized that the eATP-P2RY2 axis is a major driver of intratumoral PGE2 accumulation.

To test this hypothesis, we employed multiple humanized and syngeneic in vivo tumor models. In humanized systems, immunodeficient NXG mice were implanted with human CRC (SW480) or PDAC (BxPC-3) tumor cells, either P2RY2-proficient or -deficient, followed by treatment with antigen-matched human CAR-T or TCR-T cells (Figure 3A and B). These models provide a controlled setting to evaluate the impact of tumor-intrinsic P2RY2 inhibition on human T cell– tumor interactions in vivo.

**Figure 3.**
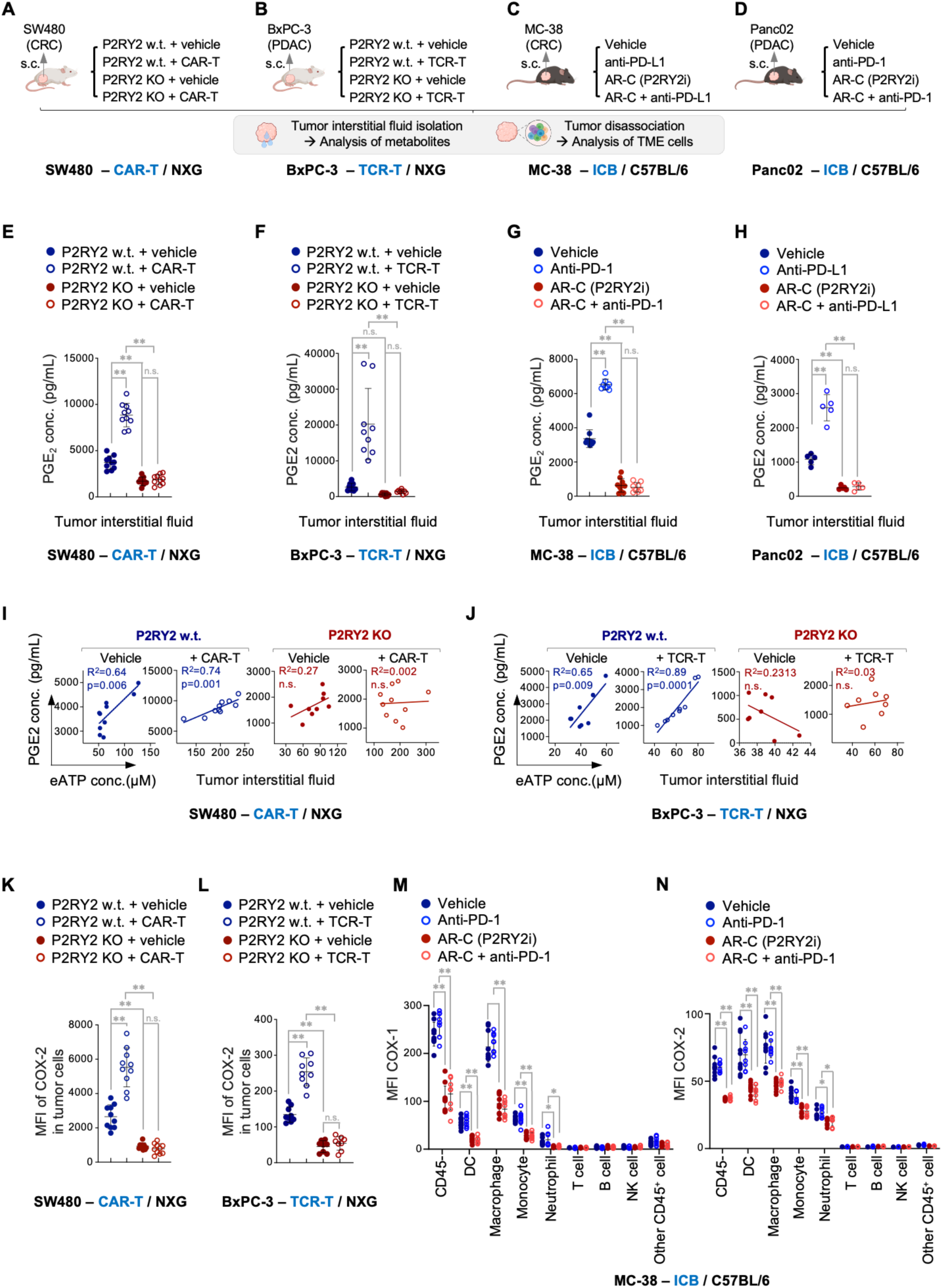
The eATP–P2RY2 axis is the primary driver of baseline and immunotherapy-induced PGE₂ accumulation in the TME. **(A)** Schematic illustration of an in vivo model of human CAR-T cell therapy against human tumors, evaluating the impact of tumor-intrinsic P2RY2 on the TME, CAR-T cell responses, and therapeutic efficacy. Human CRC SW480 cells, which constitutively express CEA, were genetically edited using CRISPR/Cas9 to generate P2RY2 knockout (KO) cells, with P2RY2 wild-type (w.t.) cells serving as controls. Tumor cells were subcutaneously (s.c.) implanted into NXG immunodeficient mice. After tumor establishment, mice bearing P2RY2 w.t. or KO tumors were treated with either vehicle control or CEA CAR-T cells. **(B)** Schematic illustration of an in vivo model of human TCR-T cell therapy against human tumors, assessing the impact of tumor-intrinsic P2RY2 on the TME, TCR-T cell responses, and therapeutic efficacy. To enable antigen-specific recognition by MART-1 TCR-T cells, human BxPC-3 cells were transduced with a fusion construct encoding the MART-1 epitope (ELAGIGILTV), HLA-A2, and β2-microglobulin (B2M) to ensure stable antigen presentation. The antigen-positive BxPC-3 cells were genetically modified using CRISPR/Cas9 to generate P2RY2 knockout (KO) cells, with P2RY2 wild-type (w.t.) cells serving as controls. The resulting tumor cells were subcutaneously implanted into NXG immunodeficient mice. After tumor establishment, mice bearing P2RY2 w.t. or KO tumors were treated with either vehicle control or MART-1 TCR-T cells. **(C)** Schematic illustration of an in vivo model of anti–PD-1 therapy against syngeneic mouse tumors, assessing the effect of systemic P2RY2 inhibition on antitumor immune responses and therapeutic efficacy. Murine CRC MC-38 cells were subcutaneously implanted into immunocompetent C57BL/6 mice. After tumor establishment, mice were treated with one of the following: vehicle control (Vehicle); PD-1 blocking antibody (anti–PD-1); P2RY2 inhibitor AR-C118925XX (AR-C); or combination therapy (AR-C + anti–PD-1). **(D)** Schematic illustration of an in vivo model of anti–PD-L1 therapy against syngeneic mouse tumors, assessing the effect of systemic P2RY2 inhibition on antitumor immune responses and therapeutic efficacy. Murine PDAC Panc02 cells were subcutaneously implanted into immunocompetent C57BL/6 mice. After tumor establishment, mice were treated with one of the following: vehicle control (Vehicle); PD-L1 blocking antibody (anti–PD-L1); P2RY2 inhibitor AR-C118925XX (AR-C); or combination therapy (AR-C + anti–PD-L1). Sample collection: Tumor tissues and the corresponding organs of origin, large intestine (A, C) and pancreas (B, D), were harvested at the endpoint. Interstitial fluid was isolated from both tumors and normal tissues. Peripheral blood was collected for plasma isolation. Tumor tissue was dissociated to generate single-cell suspensions for flow cytometry analysis. (**E**-**H**) Genetic depletion or pharmacological inhibition of P2RY2 effectively abrogates both baseline and immunotherapy-induced intratumoral PGE₂ accumulation. Tumor interstitial fluid from SW480 tumors (E), BxPC-3 tumors (F), MC-38 tumors (G), and Panc02 tumors (H), across all experimental and control groups as illustrated in Fig. 3, A to D, was collected at the endpoint. PGE₂ concentrations in the interstitial fluid were quantified by ELISA. (**I**, **J**) The positive correlation between eATP and PGE₂ concentrations in the TME is dependent on P2RY2 expression in tumors. Scatter plots depict the correlation between eATP and PGE₂ concentrations in tumor interstitial fluid from BxPC-3–TCR-T model (I) and SW480–CAR-T model (J) tumors in NXG mice. Tumors were either P2RY2 wild-type (w.t., blue) or P2RY2 knockout (KO, red) in the vehicle control (Vehicle) and CEA CAR-T cells treated (+CAR-T) groups. R² (coefficient of determination) and p-values are indicated. (**K, L**) Genetic depletion of P2RY2 in tumors diminishes both baseline and immunotherapy-induced COX-2 expression in vivo. SW480 tumors (K) and BxPC-3 tumors (L), across all experimental and control groups as illustrated in Fig. 3, A and B, were harvested at the endpoint and dissociated. COX-2 expression in tumor cells was assessed by flow cytometry. (**M**, **N**) Global P2RY2 inhibition by small-molecule P2RY2 antagonist AR-C118925XX (AR-C) reduces COX-1 and COX-2 expression in the TME. Expression levels of COX-1 (M) and COX-2 (N) in immune (DCs, macrophages, monocytes, neutrophils, T cells, B cells, NK cells, and other CD45⁺ cells) and non-immune (CD45⁻) cell populations from MC-38 tumor dissociates (as illustrated in Fig. 3C) were assessed by flow cytometry. Data are presented as mean ± standard deviation of biological replicates. For panels (E, I, K), the sample sizes are: P2RY2 wild-type (w.t.) + vehicle (n=10), P2RY2 knockout (KO) + vehicle (n=9), P2RY2 w.t. + CAR-T (n=10), and P2RY2 KO + CAR-T (n=10). For panels (F, J, L), the sample sizes are: P2RY2 wild-type (w.t.) + vehicle (n=9), P2RY2 knockout (KO) + vehicle (n=8), P2RY2 w.t. + TCR-T (n=9), and P2RY2 KO + TCR-T (n=8). For panels (G, M, N), the sample sizes are: vehicle (n=10), anti-PD-1 (n=8), AR-C (n=10), and AR-C + anti-PD-1 (n=8). For panel (H) are: vehicle (n=5), anti-PD-L1 (n=5), AR-C (n=5), and AR-C + anti-PD-L1 (n=5). Statistical significance was determined using one-way ANOVA with Tukey’s multiple comparisons test (E-H, K-N), or simple linear regression analysis (I, J). A p-value greater than 0.05 indicates non-significance (n.s.), while p < 0.05 is denoted as ∗, and p < 0.0001 is denoted as ∗∗. See also Figure S5.

Complementarily, we used syngeneic mouse models of CRC (MC-38) and PDAC (Panc02) in immunocompetent mice, treated with ICB, a selective P2RY2 inhibitor (AR-C118925XX, hereafter “AR-C”), or their combination (Figure 3C and D), enabling assessment of effect of global P2RY2 inhibition on both tumor and host (immune) compartments under immunotherapy. The functional activity of AR-C was shown to be mediated specifically through tumoral P2RY2 inhibition: AR-C treatment significantly enhanced CD8⁺ T cell activation and cytokine production when tumor cells expressed P2RY2, but failed to exert any additional effects in P2RY2-knockout cells (Figure S5A and B). Accordingly, AR-C enhanced tumor cell killing by TCR-T and CAR-T cells in a P2RY2-dependent manner, showing no benefit in P2RY2-deficient targets (Figure S5C and D).

Across all four tumor models, genetic depletion or pharmacological inhibition of P2RY2 consistently and dramatically reduced intratumoral PGE₂ levels, even lowering them below baseline (vehicle control) (Figure 3E-H), despite persistently elevated or further rising eATP concentrations in the TME (Figure S5E-H).

In P2RY2-proficient tumors, eATP concentrations strongly correlated with PGE₂ levels (Figure 3I and J, P2RY2 w.t.), whereas this correlation was completely abolished in P2RY2 deficient tumors (Figure 3I and J, P2RY2 KO), establishing P2RY2 as the critical molecular link coupling eATP accumulation to PGE₂ production within the TME.

Consistent with reduced PGE₂ in the TME, expression of COX-2 (a key enzyme responsible for PGE₂ biosynthesis) was markedly downregulated in P2RY2-deficient tumor cells, both at untreated and TCR-T or CAR-T treated conditions (Figure 3K and L). Similarly, pharmacological inhibition of P2RY2, either alone or combined with ICB, substantially decreased COX-1 and COX-2 expression, not only in tumor and stromal cell populations (CD45^−^ cells), but also in tumor infiltrating immune populations, predominantly myeloid cells (Figure 3M and N).

Taken together, these findings establish the eATP–P2RY2 signaling axis as the dominant upstream driver of intratumoral PGE₂ accumulation across tumor types and therapeutic contexts.

### Immunotherapy amplifies eATP–P2RY2–PGE₂ signaling to drive adaptive resistance

Notably, treatment with immunotherapeutic agents alone, including CAR-T, TCR-T, and ICB, further increased eATP levels in the TME, beyond the already high baseline (Figure S5E-H). This therapy-driven surge in eATP was accompanied by a marked increase in intratumoral PGE₂ (Figure 3E-H), suggesting that tumors exploit the additional eATP released during immunotherapy to amplify PGE₂-mediated immune resistance.

Strikingly, co-targeting P2RY2 completely prevented this therapy-induced PGE₂ spike and instead reduced PGE₂ levels below baseline (Figure 3E-H). These findings indicate that P2RY2 blockade not only disrupts the basal immunosuppressive environment but also circumvents an adaptive immunotherapy resistance mechanism that is broadly operative.

### P2RY2 disruption restores T cell function and overcomes resistance to CAR-T and TCR-T therapies in vivo

Having established the eATP–P2RY2 axis as a major driver of PGE₂ accumulation in the TME, at baseline and even more so during immunotherapy, we next asked whether targeting P2RY2 could relieve T cell suppression and overcome therapy resistance.

We first evaluated this in humanized tumor models, in which human CRC (SW480) or PDAC (BxPC3) tumors were treated with antigen-matched human CAR-T or TCR-T cells in immunodeficient mice (Figure 3A and B). Disruption of tumor-intrinsic P2RY2 markedly increased the abundance of adoptively transferred human CD3⁺ CAR-T and TCR-T cells within tumors and raised the proportion of CD8⁺ T cells among the infiltrating T cell populations (Figure 4A and D).

**Figure 4.**
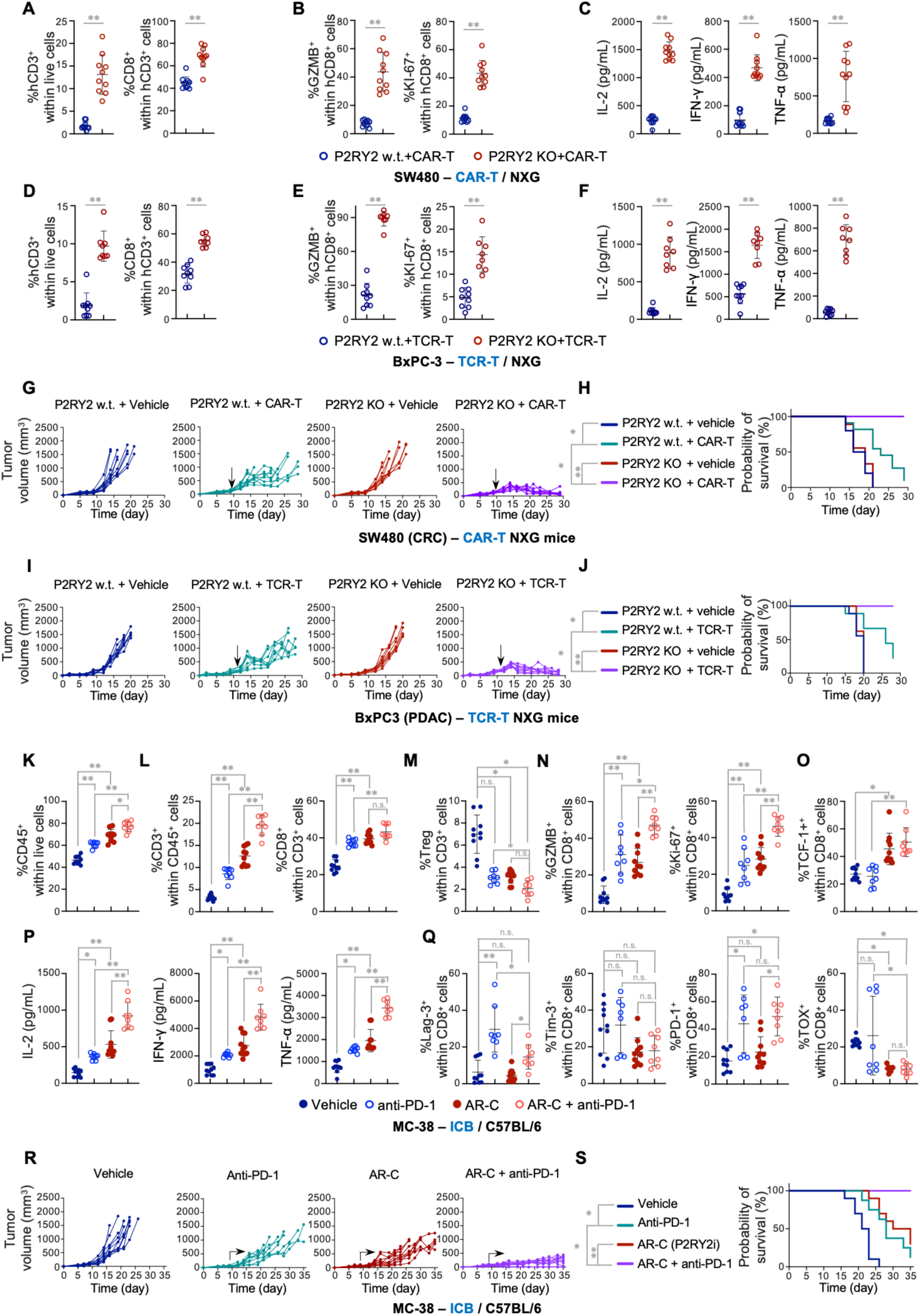
Targeting P2RY2 enhances T cell function and the efficacy of CAR-T, TCR-T, and ICB therapies in humanized and syngeneic tumor models. (**A-C**) P2RY2-proficient (P2RY2 w.t.) and -deficient (P2RY2 KO) human CRC (SW480) tumors were treated with human CEA CAR-T cells in immunodeficient NXG mice, and harvested at the end point (as described in Fig. 3A). (A) P2RY2-deficient tumors show increased abundance of CAR-T cells in the TME. The percentages of human CD3⁺ T cells within all live cells in the single-cell suspensions of SW480 tumors, as well as CD8⁺ T cells within the CD3⁺ population were quantified by flow cytometry. (B) P2RY2-deficient tumors exhibit enhanced cytotoxicity and proliferation of CAR-T cells in the TME. Granzyme B (GZMB) and Ki-67 expression in human CD8⁺ CAR-T cells within SW480 tumors were assessed by flow cytometry. (C) P2RY2-deficient tumors demonstrate increased effector cytokine accumulation in the TME. IL-2, IFN-γ, and TNF-α concentrations in tumor interstitial fluid were quantified using LEGENDplex™ assays. (**D-F**) P2RY2-proficient (P2RY2 w.t.) and -deficient (P2RY2 KO) human PDAC (BxPC-3) tumors were treated with human TCR-T cells in immunodeficient NXG mice, and harvested at the end point (as described in Fig. 3B). (D) P2RY2-deficient tumors show increased abundance of TCR-T cells in the TME. The percentages of human CD3⁺ T cells within all live cells in the single-cell suspensions of BxPC-3 tumors, as well as CD8⁺ T cells within the CD3⁺ population were quantified by flow cytometry. **(E)** P2RY2-deficient tumors show enhanced cytotoxicity and proliferation of TCR-T cells in the TME. Granzyme B (GZMB) and Ki-67 expression in human CD8⁺ TCR-T cells within BxPC-3 tumors were assessed by flow cytometry. (F) P2RY2-deficient tumors exhibit increased effector cytokine accumulation in the TME. IL-2, IFN-γ, and TNF-α concentrations in tumor interstitial fluid were quantified using LEGENDplex™ assays. (**G**, **H**) Genetic disruption of P2RY2 improves the efficacy of CAR-T therapy. Immunodeficient NXG mice bearing P2RY2-proficient (P2RY2 w.t.) and -deficient (P2RY2 KO) SW480 tumors were treated with either vehicle control or CEA CAR-T cells (as described in Fig. 3A), and tumor volume was monitored over time (G). Survival probability was assessed based on the percentage of tumor-bearing mice that remained alive throughout the study (H). CAR-T cell infusion was performed on day 9 post-tumor implantation (indicated by the arrow). (**I**, **J**) Genetic disruption of P2RY2 improves the efficacy of TCR-T therapy. Immunodeficient NXG mice bearing P2RY2-proficient (P2RY2 w.t.) and -deficient (P2RY2 KO) BxPC-3 tumors were treated with either vehicle control or MART-1 TCR-T cells (as described in Fig. 3B), and tumor volume was monitored over time (I). Survival probability was assessed based on the percentage of tumor-bearing mice that remained alive throughout the study (J). TCR-T cell infusion was performed on day 12 post-tumor implantation (indicated by the arrow). (**K-S**) Immunocompetent C57BL/6 mice bearing syngeneic mouse CRC (MC-38) tumors were treated with one of the following conditions: Vehicle control (Vehicle); PD-1 blocking antibody (anti-PD-1); small-molecule P2RY2 antagonist (AR-C); Combination therapy (AR-C + anti-PD-1), and tumors were harvested at the endpoint (as illustrated in Fig. 3C). (K, L) Pharmacological P2RY2 inhibition increases infiltration of CD45⁺ immune cells (K), as well as CD3⁺ and CD8⁺ T cells (L), in the TME. The percentage of CD45⁺ immune cells within all live cells (K), as well as CD3⁺ T cells within CD45⁺ immune cells and CD8⁺ T cells within the CD3⁺ population (L), were assessed by flow cytometry in tumor dissociates. (M) Pharmacological P2RY2 inhibition reduces the proportion of Tregs in the TME. The percentage of Tregs (CD4^+^Foxp3⁺) within CD3⁺ T cells was assessed by flow cytometry. (N) Pharmacological P2RY2 inhibition increases cytotoxicity and proliferation of CD8⁺ T cells in the TME. The expression of granzyme B (GZMB) and Ki-67 within tumor-infiltrating CD8⁺ T cells was quantified by flow cytometry. (O) Pharmacological P2RY2 inhibition increases the frequency of TCF-1⁺ progenitor-like CD8⁺ T cells in the TME. The percentage of TCF-1⁺ CD8⁺ T cells was quantified by flow cytometry. (P) Pharmacological P2RY2 inhibition enhances effector cytokine accumulation in the TME. Concentrations of IL-2, IFN-γ, and TNF-α in tumor interstitial fluid were measured using LEGENDplex™ assays. (Q) Combined P2RY2 inhibition and ICB does not increase (and even reduces) T cell exhaustion markers compared to anti-PD-1 treatment alone in the TME. The percentages of LAG-3⁺, Tim-3⁺, PD-1⁺, and TOX⁺ cells within tumor-infiltrating CD8⁺ T cells were quantified by flow cytometry. (R, S) Pharmacological P2RY2 inhibition enhances the efficacy of ICB therapy. Tumor volume was monitored over time (R). Survival probability was assessed based on the percentage of tumor-bearing mice that remained alive throughout the study (S). Therapeutic intervention was initiated on day 8 post-tumor implantation (indicated by the arrow). Data are presented as mean ± standard deviation of biological replicates. For panels (A-C, G, H), the sample sizes are: P2RY2 wild-type (w.t.) + vehicle (n=10), P2RY2 knockout (KO) + vehicle (n=9), P2RY2 w.t. + CAR-T (n=10), and P2RY2 KO + CAR-T (n=10). For panels (D-F, I, J), the sample sizes are: P2RY2 wild-type (w.t.) + vehicle (n=9), P2RY2 knockout (KO) + vehicle (n=8), P2RY2 w.t. + TCR-T (n=9), and P2RY2 KO + TCR-T (n=8). For panels (K-S), the sample sizes are: vehicle (n=10), anti-PD-1 (n=8), AR-C (n=10), and AR-C + anti-PD-1 (n=8). Statistical significance was determined using unpaired two-tailed Student’s t-test (A-F), one-way ANOVA with Tukey’s multiple comparisons test (K-Q), or Kaplan–Meier survival analysis (H, J, S). A p-value greater than 0.05 indicates non-significance (n.s.), while p < 0.05 is denoted as ∗, and p < 0.0001 is denoted as ∗∗. See also Figure S6.

Human CD8⁺ T cells in P2RY2-deficient tumors expressed higher levels of Granzyme B and Ki-67, indicating enhanced cytolytic function and proliferation (Figure 4B and E), and CD3⁺CD8⁻ T cells similarly showed elevated Ki-67 expression (Figure S6A and C).

Correspondingly, in P2RY2-deficient tumors, effector cytokines (human IL-2, IFN-γ, TNF-α) were elevated in the tumor interstitial fluid (Figure 4C and F), while T cell exhaustion markers (PD-1 and LAG-3) were unchanged or reduced (Figure S6B and D), suggesting more durable T cell responses.

The restoration of T cell functionality translated into improved therapeutic outcomes. CAR-T or TCR-T monotherapy modestly delayed tumor growth in P2RY2-proficient tumors, while P2RY2 disruption alone had no impact (Figure 4G and I), consistent with in vitro data showing that P2RY2 does not affect intrinsic tumor cell proliferation (Figure S3B). Remarkably, combing P2RY2 disruption with CAR-T or TCR-T therapy dramatically enhanced tumor suppression (Figure 4G and I) and prolonged survival (Figure 4H and J).

### Pharmacological P2RY2 inhibition remodels the TME and synergizes with ICB in syngeneic tumor models

To assess the systemic impact of global P2RY2 targeting, we next evaluated its effect in syngeneic CRC (MC38) and PDAC (Panc02) tumor models in immunocompetent mice (Figure 3C and D), enabling evaluation of both tumor- and host-intrinsic contributions.

Treatment with either AR-C (a selective P2RY2 inhibitor) or PD-(L)1 checkpoint blockade individually increased CD45⁺ immune cell infiltration into tumors, while the combination therapy induced the highest levels of infiltration (Figure 4K; Figure S6E). P2RY2 inhibition increased the abundance of CD3⁺ T cells among the infiltrating immune cells, and elevated the proportion of CD8⁺ T cells within the T cell compartment (Figure 4L; Figure S6F), with both effects further augmented by combination treatment. In contrast, the frequencies of immunosuppressive regulatory T cells (Tregs) slightly decreased or remained unchanged (Figure 4M; Figure S6G). Additionally, P2RY2 inhibition expanded intratumoral NK cells (Figure S6H and N) and reduced CD11b^+^ myeloid populations (Figure S6I and O) – the dominant COX-1/2 – expressing cells (next to tumor cells) in the TME (Figure 3M and N).

Further phenotypic analysis revealed that the combined inhibition of P2RY2 and PD-(L)1 boosted both cytolytic activity (granzyme B) and proliferation (Ki-67) of CD8^+^ T cells, outperforming either treatment alone (Figure 4N and Figure S6J). These tumor-infiltrating T cells also demonstrated strongly upregulated TCF1 expression (Figure 4O), a marker of stem-like, long-lived T cells^81–83^. Enhanced CD4⁺ T cell proliferation was also observed (Figure S6P).

In line with these findings, IL-2, IFN-γ, and TNF-α production was elevated under each monotherapy and further augmented by combination therapy (Figure 4P; Figure S6K). Importantly, in the presence of P2RY2 inhibition, exhaustion markers (Lag-3, Tim-3, PD-1, TOX) did not increase, and some even decreased, compared to controls or ICB monotherapy (Figure 4Q). Of note, the proportion of Ki-67⁺PD-1⁺ CD8⁺ T cells – a population associated with favorable responses to checkpoint blockade^84,85^, was highest in the combination treatment groups (Figure S6L and Q).

Consistent with these immunophenotypic changes, combining P2RY2 inhibition with PD-1 or PD-L1 blockade achieved profound tumor regression and the greatest survival benefit (Figure 4R and S; Figure S6M). These findings highlight P2RY2 as a central node of resistance to PD-(L)1 checkpoint blockade and demonstrate that its inhibition reshapes the TME to foster durable antitumor T cell responses.

### A selective monoclonal P2RY2-blocking antibody enhances tumor elimination by CAR-T therapy

To date, the most advanced P2RY2 antagonist, AR-C118925, exhibits poor pharmacokinetics^86–88^, limiting its translational potential. We took an alternative approach by generating monoclonal antibodies targeting P2RY2. One lead antibody selectively bound to P2RY2-expressing cells (Figure 5A) and significantly enhanced antigen-specific T cell–mediated tumor killing (Figure 5B). Both binding and functional activity were completely abolished in P2RY2-deficient cells (Figure 5A and B), confirming the antibody’s selective activity via P2RY2.

**Figure 5.**
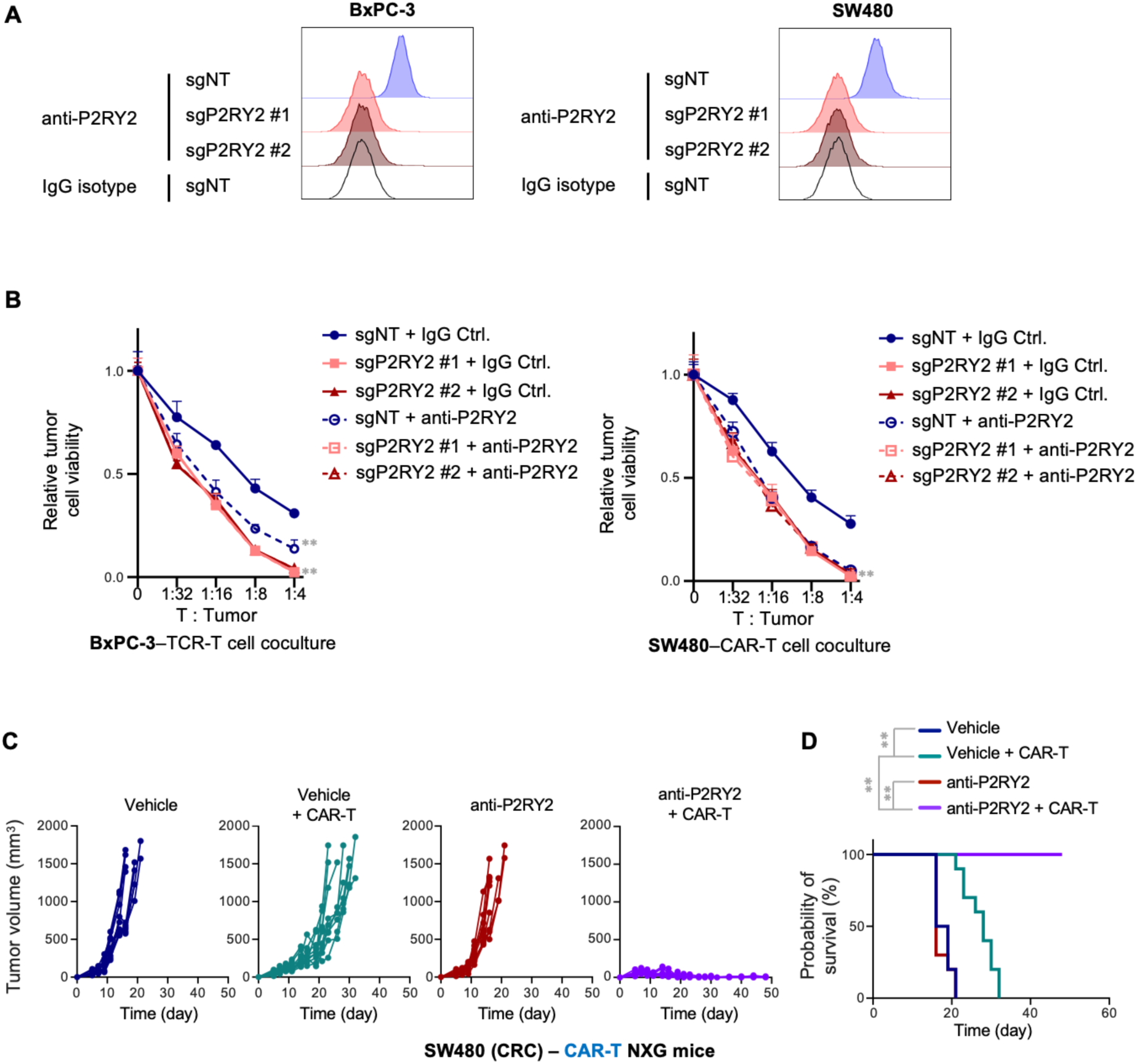
Characterization of a selective P2RY2 blocking monoclonal antibody. **(A)** The anti-P2RY2 monoclonal antibody selectively binds P2RY2-expressing tumor cells. BxPC-3 (left) and SW480 (right) cells, either P2RY2-proficient (sgNT) or P2RY2-deficient (sgP2RY2 #1 and sgP2RY2 #2), were stained with the anti-P2RY2 monoclonal antibody (clone 160#5, 10 μg/mL) or an IgG isotype control, followed by fluorophore-conjugated secondary antibody staining, and analyzed by flow cytometry. **(B)** The anti-P2RY2 monoclonal antibody enhances antigen-specific T cell–mediated tumor killing in a P2RY2-dependent manner. BxPC-3 tumor cells (MART-1 epitope⁺) were cocultured with MART-1 TCR-T cells (left), and SW480 tumor cells (CEA⁺) were cocultured with CEA CAR-T cells (right) at the indicated T cell–to–tumor cell ratios in the presence of the anti-P2RY2 antibody (clone 160#5, 1μg/mL) or an IgG isotype control. Tumor cell viability was assessed after 24 hours of coculture using the CellTiter-Blue Cell Viability Assay. P2RY2 blockade mimicked the effect of P2RY2 knockout (sgP2RY2 #1 and #2), reducing tumor cell viability following TCR-T or CAR-T cell attack. No effect of the antibody was observed in P2RY2-deficient conditions, confirming that its function is P2RY2-specific. **(C, D)** A monoclonal P2RY2 blocking antibody enhances the efficacy of CAR-T therapy. Immunodeficient NXG mice bearing SW480 tumors were treated with one of the following conditions: Vehicle control (Vehicle); CEA CAR-T cell (CAR-T) (injected on day 7); monoclonal P2RY2 blocking antibody (anti-P2RY2) (twice per week from day 6); Combination therapy (anti-P2RY2 + CAR-T). Tumor volume was monitored over time (C). Survival probability was assessed based on the percentage of tumor-bearing mice that remained alive throughout the study (D). Data are presented as mean ± standard deviation of biological replicates (n ≥ 3). For panels (C and D), the sample sizes are: Vehicle (n=10), CAR-T (n=10), anti-P2RY2 (n=10), anti-P2RY2 + CAR-T (n=10). Statistical analysis was performed using two-way ANOVA with Tukey’s multiple comparisons test (B) or Kaplan–Meier survival analysis (D). A p-value ≥ 0.05 indicates non-significance (n.s.), p < 0.05 is denoted as *, and p < 0.0001 is indicated as **.

We next evaluated the therapeutic potential of antibody-mediated P2RY2 blockade in vivo. Consistent with the effects of genetic P2RY2 deletion, antibody treatment significantly improved tumor control (Figure 5C) and extended survival of tumor-bearing mice treated with CAR-T cells (Figure 5D).

Together, these findings highlight the therapeutic potential and tractability of P2RY2 as a target for enhancing T cell-based cancer immunotherapy.

### P2RY2 blockade enhances the efficacy of autologous TILs against matched tumor cells

Finally, to further assess the clinical relevance of targeting P2RY2, we evaluated its blockade in autologous TILs – matched primary tumor cells ex vivo cultures. Given that PGE_2_ impairs TIL function and expansion in human melanoma^35,36^ and that our data establish P2RY2 as a primary driver of PGE_2_ production, we next examined whether P2RY2 blockade could enhance TIL activity against melanoma (Figure 6A).

**Figure 6.**
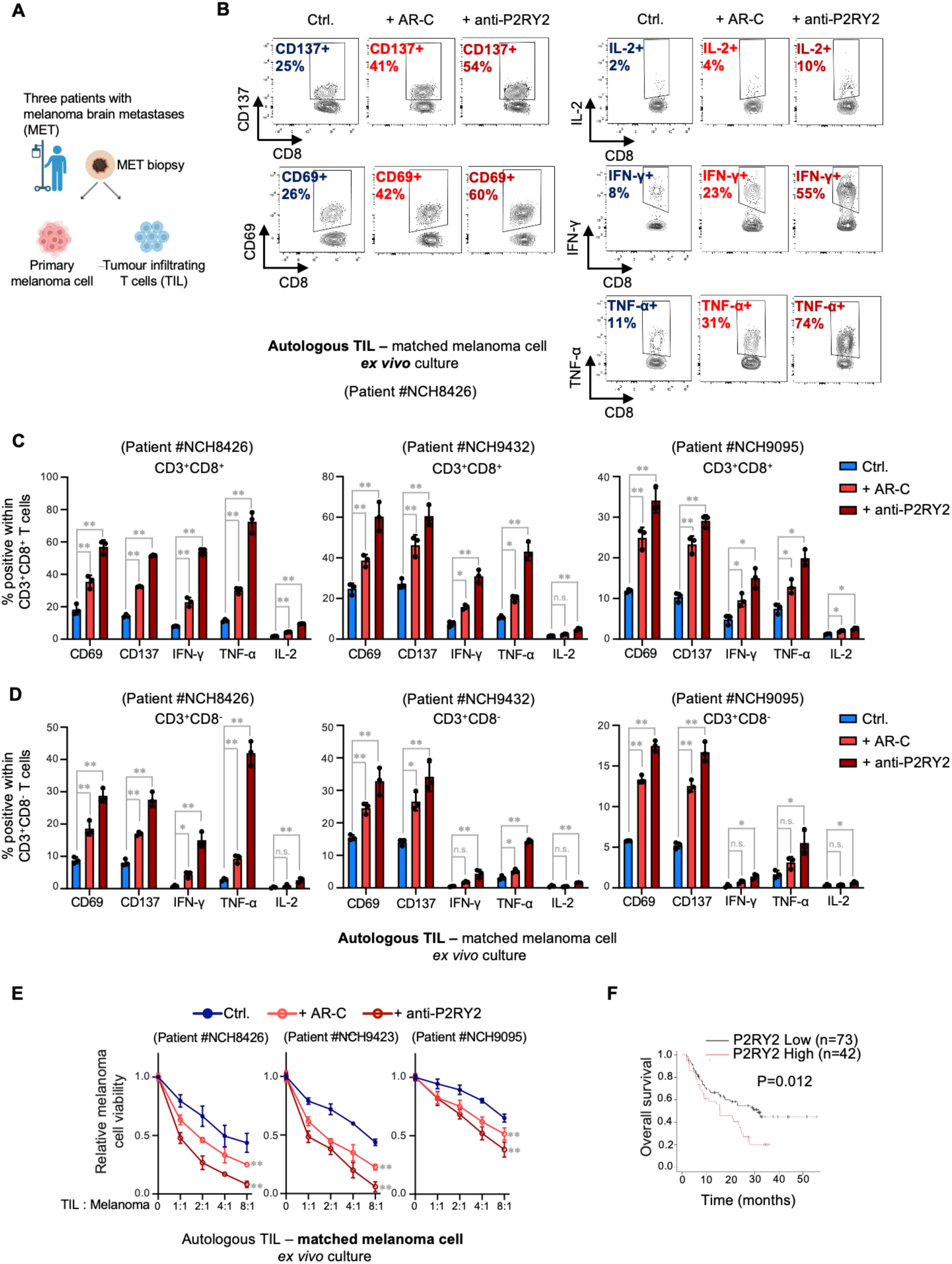
Pharmacological P2RY2 inhibition restores the activity of autologous TILs against matched primary tumor cells ex vivo. (**A**) Schematic illustration of the autologous TIL–primary melanoma cell coculture system. Biopsies of melanoma brain metastases (MET) were collected from patients. TILs and primary melanoma cells were isolated and cocultured to investigate tumor–immune interactions. (**B-D**) P2RY2 inhibition, using either a small-molecule compound or a monoclonal antibody, restores the activity of autologous TILs against matched primary melanoma cells. TILs isolated from melanoma brain metastasis were cocultured with autologous primary melanoma cells under three conditions: untreated control (Ctrl), treatment with the small-molecule P2RY2 inhibitor AR-C 118925XX (AR-C), or treatment with a newly developed monoclonal P2RY2 blocking antibody (anti-P2RY2). T cell activation markers, CD137, CD69, IL-2, IFN-γ, and TNF-α, in CD8⁺ T cells were assessed by flow cytometry. Representative contour plots for Patient #NCH8426 showing the percentage of CD8⁺ T cells expressing each activation marker are presented in (B). Quantification of activation markers in CD3⁺CD8⁺ and CD3⁺CD8⁻ TILs from all three independent patients (NCH8426, NCH9423, and NCH9095) is shown in (C, D). **(E)** Pharmacological P2RY2 inhibition enhances autologous TIL-mediated killing of matched melanoma cells. TILs isolated from three melanoma brain metastases (Patients #NCH8426, #NCH9423, and #NCH9095) were cocultured with matched primary melanoma cells at varying TIL-to-melanoma ratios, as indicated. The cocultures were treated with P2RY2 inhibitor AR-C 118925XX (AR-C), the monoclonal P2RY2 blocking antibody (anti-P2RY2), or left untreated (Ctrl.). Tumor cell viability was assessed after 72 hours using the CellTiter-Blue Cell Viability Assay. **(F)** High P2RY2 expression is associated with unfavorable overall survival in anti-PD-1-treated melanoma patients. Kaplan-Meier survival analysis of melanoma patients receiving Nivolumab (anti-PD-1) therapy, stratified by P2RY2 expression levels (low vs. high). Patients with high P2RY2 expression (n=42, red line) exhibited significantly shorter overall survival compared to those with low P2RY2 expression (n=73, black line), with a p-value of 0.012. Data represent the mean ± standard deviation of biological replicates. Statistical analysis was performed using one-way ANOVA with Dunnett’s multiple comparisons test (C, D), two-way ANOVA with Tukey’s multiple comparisons test (E) or Kaplan–Meier survival analysis (F). A p-value ≥ 0.05 indicates non-significance (n.s.), while a p-value < 0.05 is denoted as * and a p-value < 0.0001 is indicated as **.

In three independent autologous TIL–matched melanoma pairs from resected brain metastases, pharmacological blockade of P2RY2, using either AR-C or the monoclonal antibody, significantly boosted TIL activity. Enhanced expression of activation markers (CD137, CD69) and elevated effector cytokine production (IL-2, IFN-γ, TNF-α) were observed in both CD3⁺CD8⁺ (Figure 6B and C) and CD3⁺CD8⁻ TIL subsets (Figure 6D). Correspondingly, P2RY2 blockade markedly improved TIL-mediated tumor cell killing (Figure 6E). Moreover, in silico survival analysis revealed that melanoma patients with high baseline P2RY2 expression had significantly shorter overall survival following anti–PD-1 therapy compared to those with low expression (Figure 6F). This clinical correlation aligns with our findings, reinforcing the notion that P2RY2-driven immunosuppression adversely impacts immunotherapy outcomes in patients.

## DISCUSSION

Our study identifies that persistent P2RY2 signaling in the TME is a previously unrecognized mechanism driving potent resistance to diverse T cell–based cancer immunotherapies. Mechanistically, the abnormally high levels of eATP in solid tumors chronically activates P2RY2, which in turn serves as the primary upstream driver of COX-1/2-mediated PGE₂ accumulation in the TME. This discovery addresses a long-standing question regarding the primary trigger of sustained COX activity and elevated PGE₂ levels in solid tumors.

These findings provide a critical addition to our understanding of how the highly enriched eATP in the TME influences immunotherapy responsiveness. While ATP catabolism into adenosine is a well-recognized immunosuppressive pathway, our results demonstrate that P2RY2 inhibition restores antitumor T cell responses and substantially enhances immunotherapy efficacy without reducing adenosine levels (Figure S6R-U). Moreover, although acute eATP exposure can directly stimulate T cells via P2X receptors during early activation^20^, our data show that chronic eATP accumulation in tumors predominantly promotes T cell suppression by acting on the TME. This reveals a distinct and previously underappreciated mechanism of immune evasion, whereby persistently elevated eATP, independent of its hydrolysis to adenosine, dominantly suppresses immune responses via P2RY2 signaling in the TME.

We further uncover a paradoxical feedback loop in which T cell–based immunotherapies inadvertently amplify the eATP–P2RY2–PGE₂ pathway. In our models, treatment with adoptive T cells (CAR-T or TCR-T) or ICB led to additional tumor cell stress and death, which in turn released bursts of ATP, further elevating intratumoral eATP and PGE₂ levels (Figure 3E-H; Figure S5E-H). This therapy-induced surge of eATP and consequent PGE₂ dampened T cell function and fostered adaptive resistance, effectively creating a self-limiting cycle to the immunotherapy. Strikingly, P2RY2 blockade broke this feedback loop: inhibiting P2RY2 prevented the accumulation of PGE₂ despite continued cell death, thereby preserving T cell activity and antitumor efficacy. These results highlight P2RY2 inhibition as a promising strategy to sustain and enhance the effectiveness of existing immunotherapies by preemptively disabling a key resistance circuit.

Our results across multiple experimental systems underscore the broad therapeutic potential of targeting P2RY2. In diverse humanized and syngeneic mouse tumor models, as well as ex vivo human tumor cell–autologous TIL cocultures, P2RY2 inhibition markedly enhanced T cell activity and improved tumor control when combined with CAR-T, TCR-T, ICB, or TIL treatment. We achieved these benefits using a selective small-molecule antagonist (AR-C118925XX) and a newly developed P2RY2-blocking monoclonal antibody, demonstrating the druggability of P2RY2 and its translational promise. Consistent with these findings, analysis of patient tumor samples revealed that high P2RY2 expression correlates with poor responses to anti–PD-1 therapy, highlighting the clinical relevance of this pathway in human immunotherapy resistance.

Furthermore, targeting P2RY2 offers a tumor-selective strategy to counteract PGE₂-mediated immunosuppression. While hyperactivity of the COX-PGE₂ axis is strongly associated with poor outcomes across cancer types, COX enzymes also play critical homeostatic roles in normal tissues. Consequently, systemic COX or prostaglandin inhibition has been linked to significant side effects, including increased thrombotic cardiovascular events such as myocardial infarction^69–71^, gastrointestinal ulceration and bleeding^72–74^, and kidney injury^75–77^. Moreover, clinical efforts to inhibit PGE₂ signaling in combinatorial cancer immunotherapies trials have largely focused on targeting COX enzymes or downstream PGE₂ receptors, but these approaches have yielded mixed results^64–68^. In contrast, P2RY2 signaling is predominantly active within the TME, driven by the abnormally high eATP concentrations, and remains largely quiescent in healthy tissues where eATP levels are minimal. Thus, P2RY2 inhibition disrupts a tumor-specific immunosuppressive circuit while sparing normal immune and physiological functions. Supporting this, systemic P2RY2 blockade with the tool compound AR-C118925XX dramatically lowered intratumoral PGE₂ levels without observable adverse effects, consistent with the minimal phenotypic abnormalities reported in P2ry2-knockout mice^89^.

Despite the therapeutic promise, no P2RY2 inhibitor has yet advanced to the clinic. The only available selective tool compound, AR-C118925XX, exhibits poor pharmacokinetics, which limits its translational potential. To overcome this challenge, we developed a monoclonal antibody against P2RY2, which exhibits high specificity and potent antagonistic activity in in vivo (Figure 5C and D), and ex vivo (Figure 6B-E) tumor models. This proof-of-concept biologic not only reinforces the druggability of P2RY2 but also provides a potential translational path forward for targeting P2RY2 in patients.

Although our study focused on solid tumors, the eATP–P2RY2–mediated immune evasion may also be relevant in certain hematologic malignancies. Notably, elevated eATP levels have been documented in the leukemic bone marrow niche^90^, raising the possibility that P2RY2-driven suppression could operate in that context as well, and P2RY2 blockade might similarly enhance immunotherapeutic efficacy against leukemias.

In summary, we uncover a previously underappreciated mechanism of immune evasion whereby tumors exploit elevated eATP in the TME to drive immunosuppression through P2RY2 signaling. Notably, we identify the eATP–P2RY2 axis as the primary driver of intratumoral COX hyperactivation and PGE₂ accumulation, filling a critical gap in identifying the primary origin of sustained COX–PGE₂ activity in solid tumors. This pathway not only mediates baseline immune escape but also promotes adaptive resistance driven by therapy-induced ATP release and subsequent PGE₂ surges. Together, our findings establish P2RY2 inhibition as a broadly applicable and tumor-selective strategy to overcome immune resistance, offering a promising avenue to improve patient responses across a wide range of T cell–based immunotherapies.

## RESOURCE AVAILABILITY

### Lead contact

Further information and requests for resources and reagents should be directed to and will be fulfilled by the lead contact, Chong Sun.

### Materials availability

Requests for materials should be directed to the corresponding author. Material transfer will be conducted under a materials transfer agreement (MTA) between the German Cancer Research Center (DKFZ) and the recipient’s host institution. Additional information required for data reanalysis is available from the corresponding author upon request.

### Data and code availability

The transcriptomic data generated in this study have been deposited in the Gene Expression Omnibus (GEO) under accession number GSE289055. Expression data for patients receiving immune checkpoint blockade (ICB) therapy are publicly available at KM plotter. Source data from this study were deposited on Mendeley as indicated in the Key Resources Table. Additional information required for data reanalysis is available from the lead contact upon request.

## ACKNOLEDGEMENT

We thank the core facilities at the German Cancer Research Centre (DKFZ) in Heidelberg, including Flow Cytometry, the Centre for Preclinical Research, Next-Generation Sequencing, Proteomics, and Light Microscopy facilities, for their support. This work is supported by the European Research Council: ERC starting grant, DRILL, 101078722 (to CS) (Funded by the European Union. The views and opinions expressed are those of the authors only and do not necessarily reflect those of the European Union or the European Research Council Executive Agency. Neither the European Union nor the granting authority can be held responsible for them); DKFZ/Bayer Innovation Alliance grant P2RY2 (to CS); DKFZ-MOST (Israel Ministry of Science and Technology) cooperation program grant, Ca 208 (to CS).

## AUTHOR CONTRIBUTIONS

Conceptualization: CS, ZH; Methodology: ZH, HM, SD, CBB, LJ, RC, MS, XZ, BM, CE, HE, MAAM, EL, IH, CH, CS; Investigation: ZH, HM, LJ, YX, IH, CH, CS; Writing – original draft: CS, ZH; Writing – review & editing: ZH, HM, SD, CBB, LJ, RC, MS, XZ, BM, CE, HE, MAAM, EL, YX, IH, CH, CS; Funding Acquisition: CS; Supervision: CS, IH, CH; Project administration: CS.

## STAR★METHODS

### Key resources table

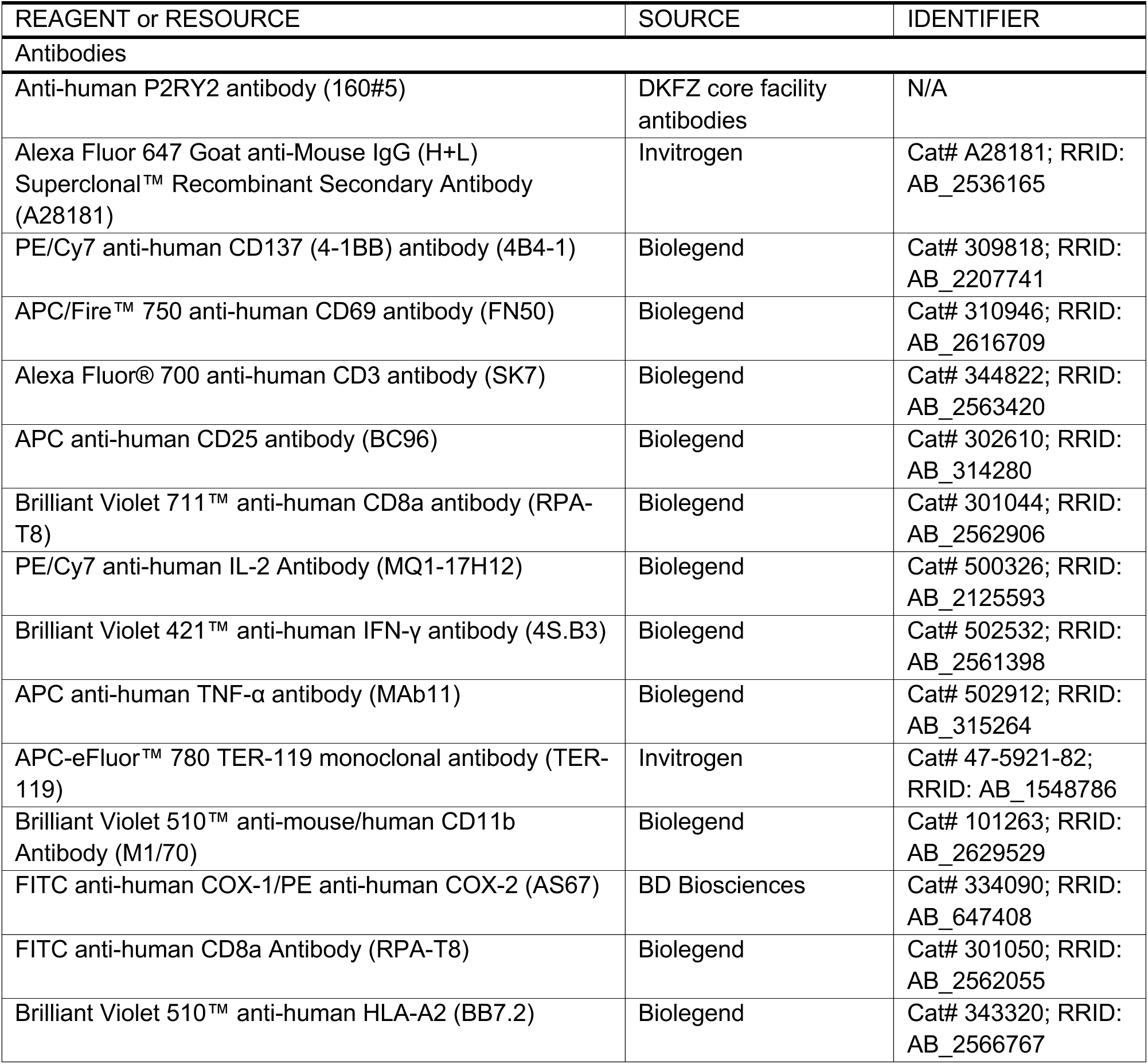
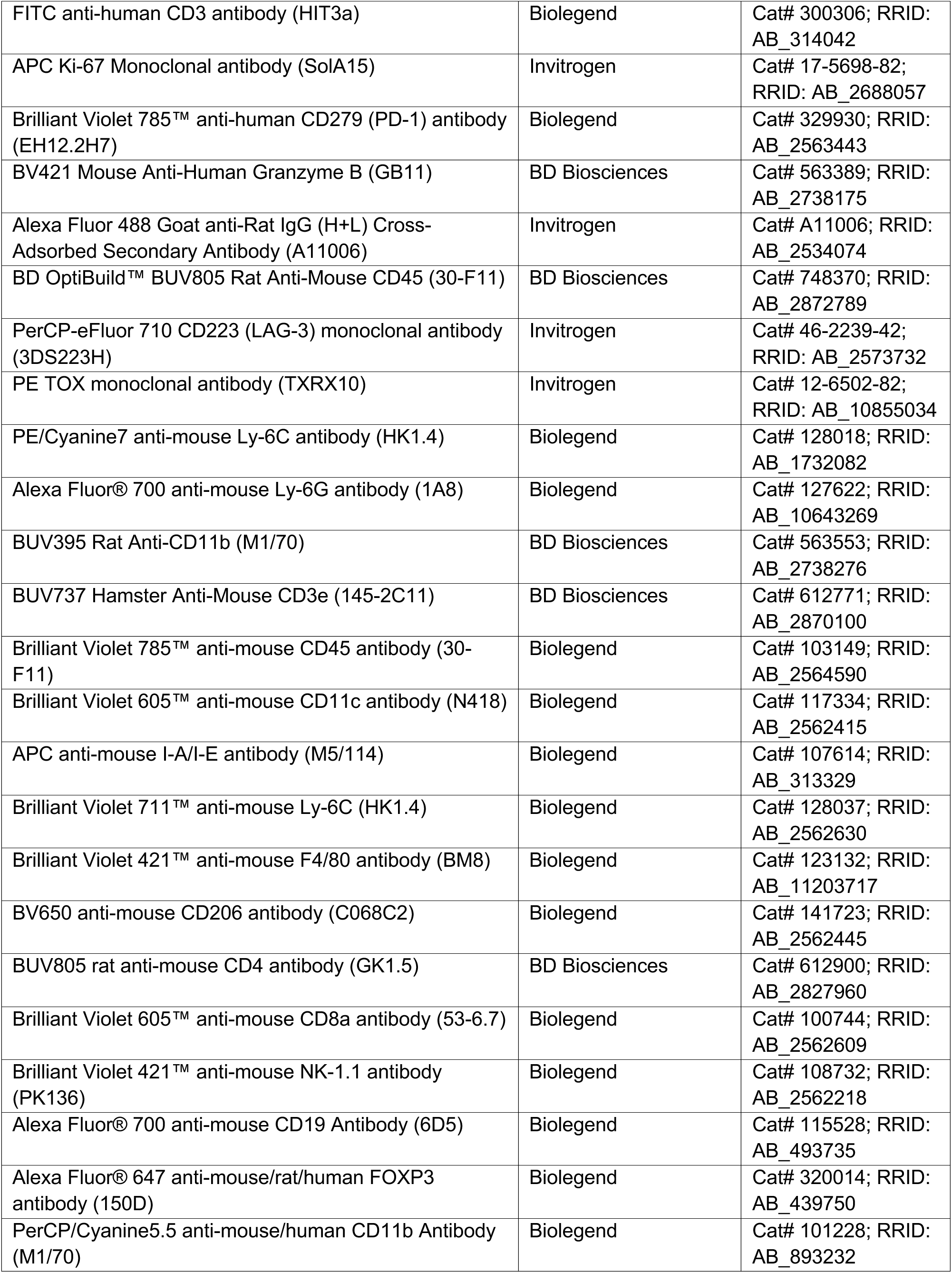
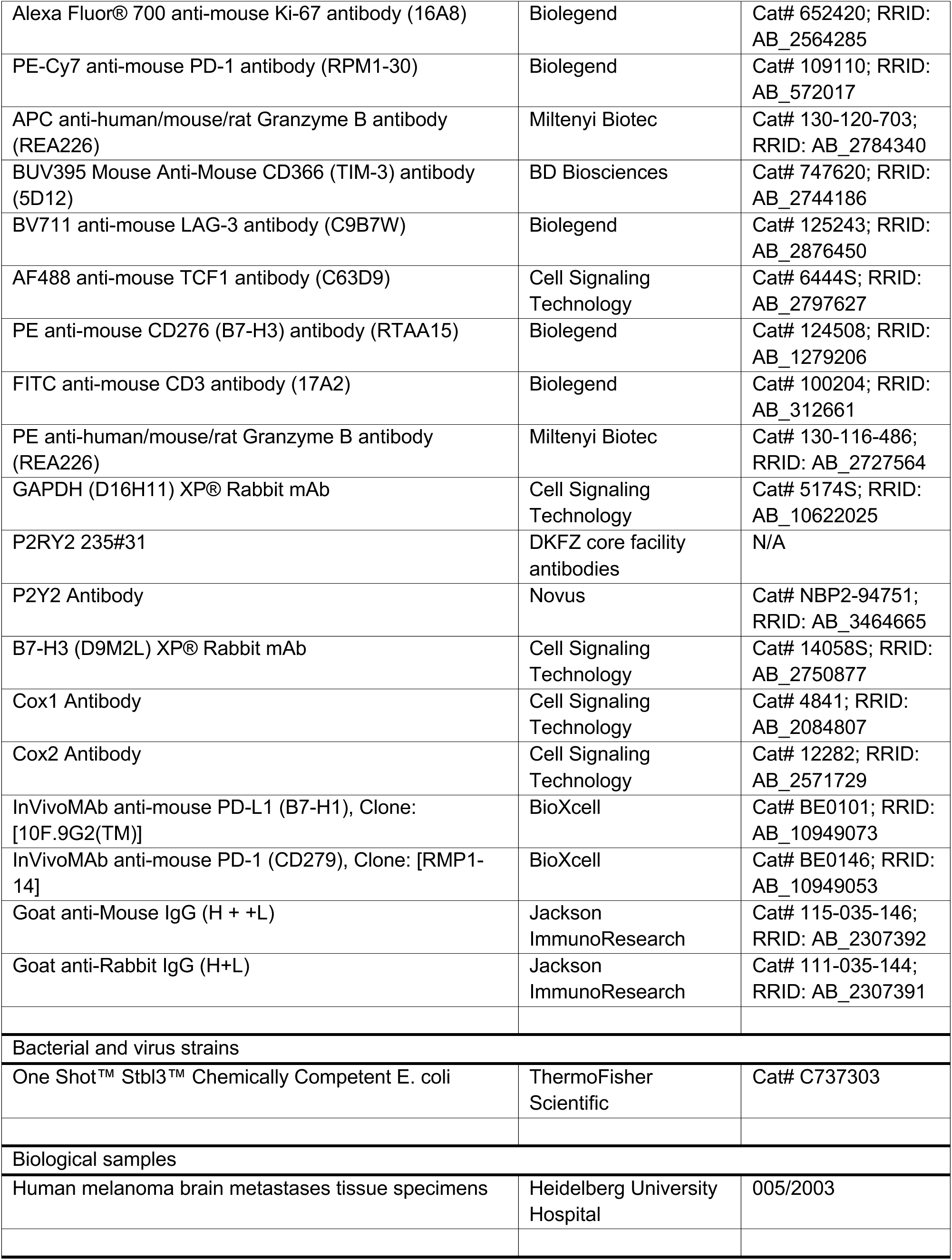
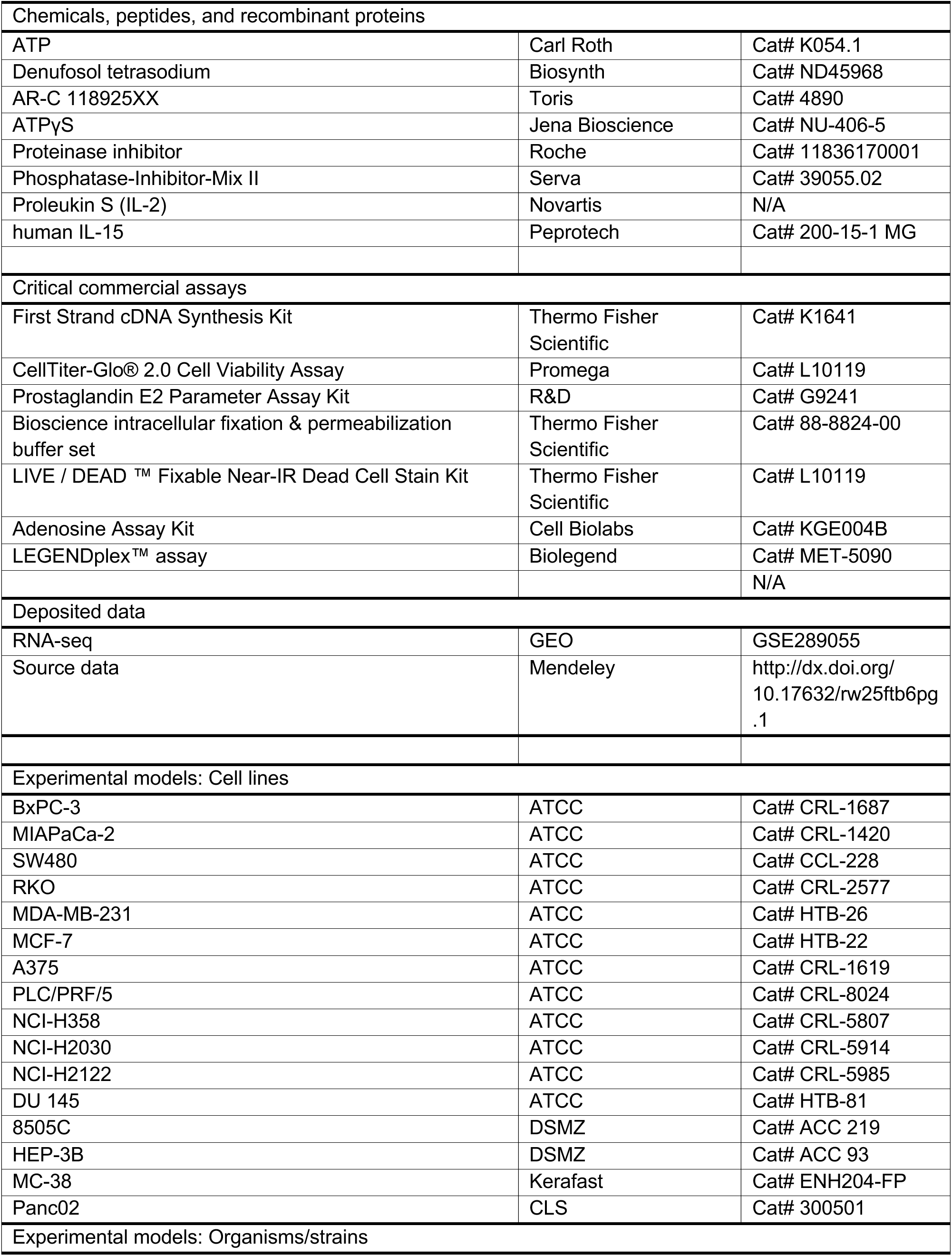
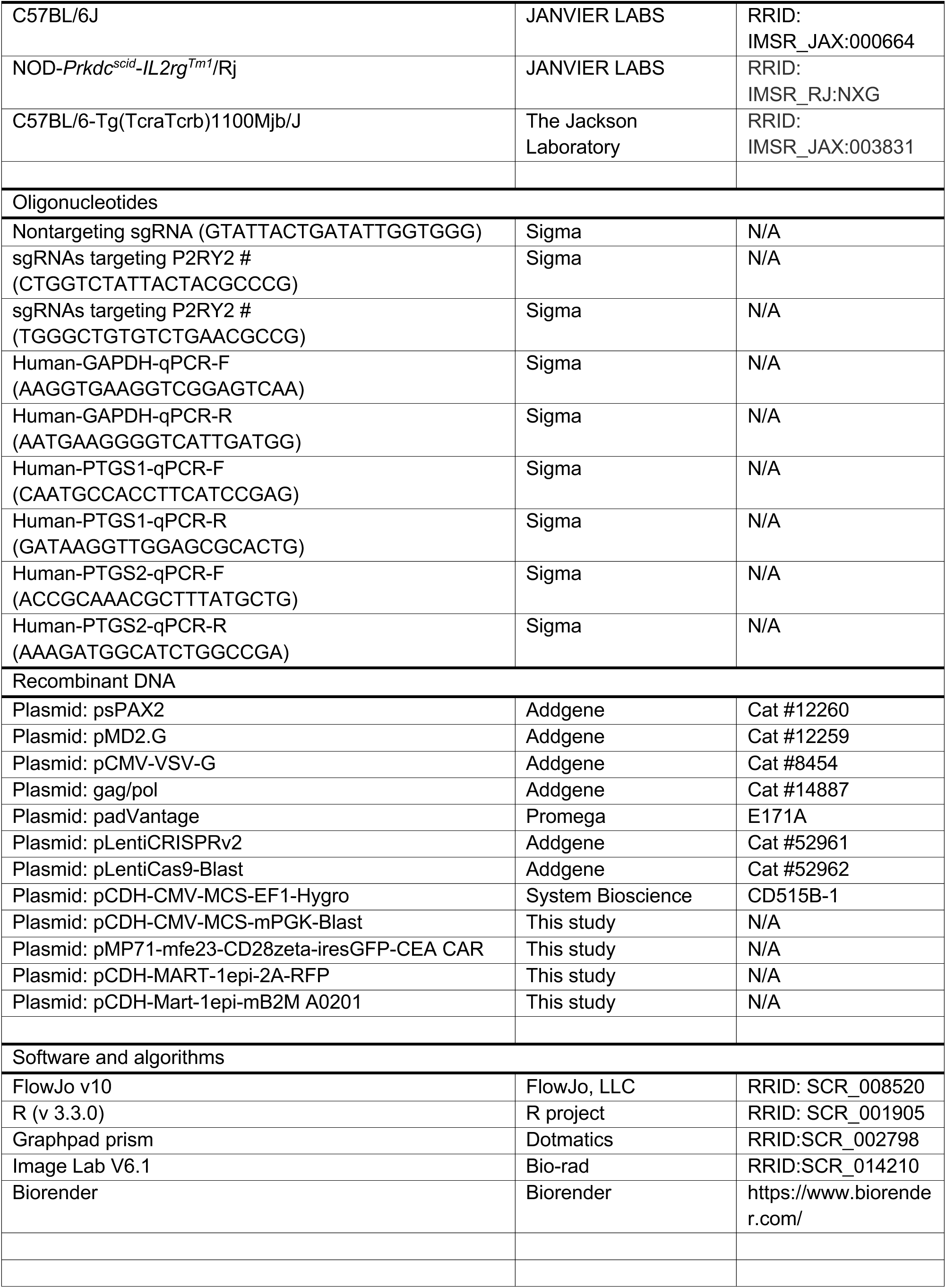

### Experimental model and study participant details

#### Arrayed CRISPR/Cas9 screen

To generate single-gene knockouts targeting specific purinergic receptors in BxPC-3 cells, a Cas9-expressing BxPC-3 clone was transduced with individual sgRNAs listed in Table S1. Prior to sgRNA transduction, BxPC-3 cells were engineered to express a fusion construct encoding the MART-1 epitope (ELAGIGILTV), HLA-A2, and β2-microglobulin (B2M), to enable antigen-specific interactions between tumor cells and T cells. Following sgRNA transduction, cells were subjected to puromycin selection for three days to enrich for successfully transduced populations. Puromycin was then withdrawn, and cells were allowed to recover for an additional three days.

For tumor cell–T cell coculture assays, 5,000 BxPC-3 cells were seeded per well in flat-bottom 96-well plates and incubated overnight to allow cell adhesion. Subsequently, 5,000 T cells were added to each well. After 24 hours of coculture, T cells were harvested and analyzed by flow cytometry to assess activation markers and effector cytokine production.

#### Cell lines

BxPC-3, MIAPaCa-2, SW480, RKO, MDA-MB-231, MCF-7, A375, PLC/PRF/5, NCI-H358, NCI-H2030, NCI-H2122, and DU-145 cells were obtained from the American Type Culture Collection (ATCC). 8505C, and HEP-3B cells were obtained from the Deutsche Sammlung von Mikroorganismen und Zellkulturen GmbH (DSMZ). MC-38 cells were obtained from Kerafast, and Panc02 cells were obtained from CLS. BxPC-3, MIAPaCa-2, SW480, RKO, MCF-7, A375, Mel526, 8505C, HEP-3B, PLC/PRF/5, NCI-H358, NCI-H2030, NCI-H2122, and DU-145 cells were cultured in RPMI 1640 medium (Thermo Fisher Scientific, Cat# 21875091). MDA-MB-231 cells were cultured in DMEM (Thermo Fisher Scientific, Cat# 11965092). All culture media were supplemented with 10% fetal bovine serum (FBS, Thermo Fisher Scientific, Cat# 10270-106), 100 U/mL penicillin-streptomycin (Thermo Fisher Scientific, Cat# 15140122), and GlutaMAX (Thermo Fisher Scientific, Cat# 15140122). Mouse Panc02 and MC-38 cells were cultured in RPMI 1640 medium (Thermo Fisher Scientific, Cat# 21875091) supplemented with 10% FBS (Thermo Fisher Scientific, Cat# 10270-106), 100 U/mL penicillin-streptomycin (Thermo Fisher Scientific, Cat# 15140122), GlutaMAX (Thermo Fisher Scientific, Cat# 15140122), and 55 μM β-mercaptoethanol.

#### Flow cytometry

Cell suspensions were labelled with conjugated antibodies as described in the *Antibodies and compounds* section. Dead cells were excluded using either 4′,6-diamidino-2-phenylindole (DAPI) incorporation or the LIVE/DEAD™ Fixable Near-IR Dead Cell Stain Kit (Thermo Fisher Scientific, Cat# L10119). All washes and reagent dilutions were performed in PBS supplemented with 2% FCS. Data acquisition was carried out using a BD LSR Fortessa HTS Wolfgang cytometer (BD Biosciences) operated with the FACS-Diva software system. Flow cytometry data were further analyzed using FlowJo software.

#### Generation of knockout and ectopic expression cell lines

Knockout cell lines were generated using the CRISPR/Cas9 system. Parental cells were transduced with the pLentiCRISPR v2 vector (Addgene, #52961) containing either a nontargeting sgRNA (GTATTACTGATATTGGTGGG) or sgRNAs targeting P2RY2. For human P2RY2 knockout, the following sgRNA sequences were used: sgP2RY2 #1: CTGGTCTATTACTACGCCCG; sgP2RY2 #2: TGGGCTGTGTCTGAACGCCG. Transduced cells were subjected to puromycin selection, and were further sorted by FACS. Knockout efficiency was validated using Western blot or flow cytometry.

To generate cell lines with ectopic expression of P2RY2, the coding sequences were obtained from Ensembl, codon-optimized, synthesized by Integrated DNA Technologies or Twist Bioscience, and clonedinto the pCDH-EF1α-MCS-EF1-Blast vector, a modified version of pCDH-EF1α-MCS-EF1-Hygro (System Biosciences, #CD510B-1).

Lentivirus production was performed by co-transfecting HEK293T cells with the transfer plasmids containing the target sequences and the following packaging plasmids: pPAX2 (Addgene, #12260) and pVSV-G (Addgene, #12259). Two days post-transfection, the lentiviral supernatant was collected and used for cell transduction.

#### RNA sequencing data analysis

Raw reads from the paired-end fastq files were aligned to GRCh38 using STAR (2.7.10a). Raw read counts per gene were measured by FeatureCounts (2.0.1) and were normalised to count per million. Ensemble IDs were converted to gene symbols using biomaRt (2.48.3) in R. Differentially expressed genes were identified using DeSeq2 (1.40.2).

#### Reverse transcription-quantitative polymerase chain reaction (RT-QPCR)

RNA was extracted using the RNeasy Plus Mini Kit (Qiagen, Cat# 74134) according to the manufacturer’s protocol. cDNA synthesis was then performed with the Maxima First Strand cDNA Synthesis Kit (Thermo Fisher Scientific, Cat# K1641). mRNA transcripts were amplified using gene-specific primers and SYBR Green Master Mix (Thermo Fisher Scientific, Cat# A25742), with quantification carried out on a ViiA 7 Real-Time PCR System (Applied Biosystems). Relative mRNA expression levels were normalized to GAPDH as the reference gene. The primers used for PCR are listed in the key resources table.

#### Quantification of eATP, PGE_2_, adenosine, and cytokines

Extracellular ATP level was determined by CellTiter-Glo® 2.0 Cell Viability Assay (G9241, Promega) according to the manufacturer’s instructions. PGE_2_ concentration was quantified by Prostaglandin E2 Parameter Assay Kit (KGE004B, R&D). Adenosine concentration was determined by Adenosine Assay Kit (MET-5090, Cell Biolabs) according to the manufacturer’s instructions. Mouse and human cytokines were determined by LEGENDplex™ assay (Biolegend) according to the manufacturer’s instructions.

#### Western Blot Analysis

Cells for Western blot analysis were seeded in 6-well plates and treated as described in the figure legends. Protease inhibitor (cOmplete Mini EDTA-free, Roche, Cat# 11836170001) was freshly added to RIPA lysis buffer. Cells were washed with PBS before lysis with RIPA buffer. The lysate was collected using a cell lifter (Corning, Cat# CLS3008-100EA), incubated on ice for 30 minutes, and then centrifuged at 14,000 rpm for 15 minutes at 4°C. The supernatant was collected, and protein concentration was determined using the BCA Protein Assay Kit (Thermo Scientific, Cat# 71285-3). Normalized protein samples were loaded onto the NuPAGE Gel Electrophoresis System (Invitrogen, Cat# WG1403BOX) and processed according to the manufacturer’s instructions.

#### MART-1-TCR-T and CEA CAR-T generation

MART-1-specific T cells (TCR clone #1D3) were generated as described by Jorritsma et al. (2007)^95^. The TCR construct was generously provided by Prof. Ton Schumacher and Prof. John Haanen (Netherlands Cancer Institute).

To generate CEA CAR-T cells, 7 million HEK293T cells were seeded in a 10 cm culture dish one day before transfection. Transfection was performed using a plasmid mixture containing: 7.5 μg of the retroviral transfer plasmid: pMP71_MFE-23-CD28z-ires-GFP (CEA CAR construct, ScFv of MFE-23 described in patent US5876691A); 4.5 μg of pGag/pol; 2.94 μg of pVSV-G; 1.92 μg of pAdvanced plasmid; 50 μL of PEI MAX reagent was used for transfection. Retroviral supernatant was collected and added to Retronectin-treated 24-well plates (5 μg/mL Retronectin; overnight incubation at 4°C). Plates were centrifuged at 3,500 rpm for 2 hours with no deceleration. After centrifugation, the supernatant was removed, and 2.5 × 10⁵ T cells were added per well in 500 μL of T cell medium (RPMI 1640 supplemented with 10% FBS, 5 ng/mL IL-15, and 50 U/mL IL-2). On the following day, an additional 500 μL of fresh T cell medium was added. Transduction efficiency was assessed by flow cytometry before use in coculture experiments.

#### Cocultures of tumor cells and T cells

For each well of a 6-well plate, 50,000–200,000 tumor cells were seeded, while for 96-well plates, 5,000– 10,000 tumor cells were seeded. T cells and antigen-expressing tumor cells were mixed at ratios ranging from 1:1 to 1:32 as indicated. After incubation at 37°C and 5% CO₂ for the specified durations, T cells were collected and stained for T cell activation markers, which were analysed using flow cytometry. To assess cytokine expression in T cells, Brefeldin A was added 6 hours before harvesting. Tumor cell viability was assessed using either the CellTiter Blue Assay (Promega, Cat# G8020).

#### T cell activation in conditional medium with 10 kDa filtering

Tumor cells were seeded at 200,000/mL and incubated overnight. Once adherent, the supernatant was removed and replaced with serum-free medium, followed by incubation for 24 hours. The conditional medium was then collected and centrifuged at 1,800 rpm for 5 minutes at 4°C to remove tumor cells. The supernatant was transferred to an Amicon® Ultra Centrifugal Filter (10 kDa MWCO) and centrifuged at 4,000g for 10 minutes at 4°C to isolate components larger or smaller than 10 kDa. The isolated medium was then supplemented with 10% FBS and 0.1 µg/mL anti-CD3 (OKT-3, Bio X Cell) antibody before incubation with CD3⁺ T cells for 8–24 hours. T cell activation was subsequently assessed by FACS analysis.

#### Mice

All animal experiments were conducted in accordance with protocols approved by the Ethical Committee of the Karlsruhe Regional Council. The experiments adhered to local and national guidelines and regulations. For the study, we purchased 8-week-old female NXG immunodeficient mouse NOD-Prkdcscid-IL2rgTm1/Rj mice (JANVIER LABS) or immune competent C57BL/6J mice (JANVIER LABS or Charles River). Mice were housed under Specific Pathogen-Free (SPF) conditions in the Central Animal Laboratory at DKFZ Heidelberg and were randomly assigned to experimental groups.

#### Humanized tumor models

For experiments with humanized tumor models, NXG immunodeficient mice (NOD-Prkdcscid-IL2rgTm1/Rj) from JANVIER LABS were used. 5 × 10⁶ sgNT/P2RY2-KO BxPC-3 or SW480 cells, suspended in a maximum volume of 200 µL of 50% Matrigel, were injected subcutaneously (s.c.) into NXG mice. When tumors reached a size of 50–300 mm³ (day 12 for SW480 xenografts, day 9 for BxPC-3 xenografts), 5 × 10⁶ MART-1 TCR-T or CEA CAR-T cells were injected intratumorally (i.tu.) into tumor-bearing mice in 200 µL of PBS. Mice received an intraperitoneal (i.p.) injection of IL-2 (6.5 × 10^4^ IU) and IL-15 (1 µg), dissolved in 200 µL of PBS, once daily starting one day prior to T cell transfer and continuing for three consecutive days. Mice were sacrificed if tumor volume exceeded 1,500 mm³ or if any tumor dimension exceeded 1.5 cm, as measured using a calliper gauge. Tumor volume was calculated using the formula: Volume = 1/2 × Length × Width^2^. Tumors, mouse tissues, and organs were harvested at the end point. Tumors were dissociated by Tumor Dissociation Kit (130-095-929, Miltenyi Biotec) according to the manufacturer’s instructions for further flow cytometry analysis.

#### Syngeneic tumor models

1 × 10⁶ Panc02 or MC-38 cells, suspended in 100 µL of 50% Matrigel, were injected subcutaneously (s.c.) into the right flank of mice. When tumors reached 50–300 mm³, tumor-bearing mice were randomized into four groups: 1. aPD-1/aPD-L1 group: Mice received 200 µg/mouse intraperitoneal (i.p.) treatment of either anti-PD-1 or anti-PD-L1 blocking antibodies (InVivoMAb anti-mouse PD-1 (CD279), Clone: [RMP1-14]; InVivoMAb anti-mouse PD-L1 (B7-H1), Clone: [10F.9G2(TM)], BioXcell) twice per week. 2. AR-C group: Mice received 20 mg/kg i.p. treatment of AR-C 118925XX once daily. 3. AR-C + aPD-(L)1 group: Mice received both AR-C 118925XX and aPD-1/aPD-L1 treatments. 4. Control (Vehicle) group: Mice received vehicle injections at the same frequency as the treatment groups. Tumor volume was monitored every 2–3 days. Mice were sacrificed if tumor volume exceeded 1,500 mm³ or if any tumor dimension exceeded 1.5 cm, as measured using a calliper gauge. Tumor volume was calculated using the formula: Volume = 1/2 × Length × Width^2^. Mouse tissues and organs were harvested at the end point. Tumors were dissociated by Tumor Dissociation Kit (130-095-929, Miltenyi Biotec) according to the manufacturer’s instructions for further flow cytometry analysis.

#### Collection of interstitial fluid

The method for extracting tumor interstitial fluid was adapted from Matas-Nadal et al., 2019^96^. Tumors or organs were isolated and washed once with PBS to remove any blood or surface fluids. Excess PBS was gently blotted with tissue paper, and the samples were immediately transferred to 1.5 mL centrifuge tubes. These steps were performed rapidly to minimize evaporation. Samples were then centrifuged at 10,000g for 20 minutes at 4°C, after which 5–50 µL of interstitial fluid, accumulated at the bottom of the tube, was carefully recovered.

#### P2RY2 monoclonal antibody generation

Monoclonal anti-P2RY2 antibody were generated with the support of the Core Facility Antibodies of the German Cancer Research Center (DKFZ), according to the principles of Köhler and Milstein’s hybridoma technology^97^. In brief, mouse immunization was performed by genetic immunization^98^. Anti P2RY2 antibody-producing B-lymphocytes isolated from positively reacted mice were fused with Sp2/0 murine myeloma cells (RRID: CVCL_2199) to produce hybridoma cell clones. The supernatant from validated monoclonal clones was used.

#### Generation of primary tumor cells from human melanoma brain metastases

Human melanoma brain metastases tissue specimens were resected at the Department of Neurosurgery, Heidelberg University Hospital, and processed immediately. All patients gave their written informed consent in accordance with the Declaration of Helsinki and its later amendments and the study was approved by the ethics committee of the Medical Faculty, University of Heidelberg (reference number: 005/2003). Single-cell suspensions were prepared by combined enzymatic and mechanical dissociation using a gentleMACS Octo Dissociator in Hanks′ Balanced Salt solution (HBSS, Thermo Fisher Scientific, #14025050) and used to generate adherent primary tumor cells by cultivation in DMEM, high glucose, GlutaMAX™ supplement (Thermo Fisher Scientific, #61965059), 1 % Penicillin-Streptomycin (10.000 U/ml, Thermo Fisher Scientific, #15140122), 10 % FCS (Merck, #S0615). Melan-A expression was confirmed by flow cytometry (Melan-A Antibody (A103), Santa Cruz, #sc-20032, 1:100 dilution in permeabilization buffer, ThermoFisher Scientific, #88-8824-00).

#### Isolation of TILs from human melanoma brain metastases

TILs were isolated from cryo-preserved human melanoma brain metastasis-derived single-cell suspensions. T cells were isolated using the EasySep™ Human CD3 Positive Selection Kit II (Stemcell, # 17851) and an EasySep™ magnet (Stemcell, #18000) according to manufactureŕs instructions. T cells were stimulated once by 25 µl / ml ImmunoCult™ Human CD3/CD28/CD2 T Cell Activator (Stemcell, #10970) and further expanded for one week in T cell medium (ImmunoCult™-XF T Cell Expansion Medium (Stemcell, #10981), 0.2 % Penicillin-Streptomycin (10.000 U/ml, Thermo Fisher Scientific, #15140122), 6000 U/ml IL2 (Stemcell #78036.3)

#### Survival analysis of melanoma patients who received anti-PD-1 treatment

The Kaplan–Meier Plotter database (http://kmplot.com/analysis/) was used to assess the relationship between P2RY2 mRNA levels and overall survival (OS) in a melanoma cohort receiving Nivolumab treatment. To generate Kaplan–Meier survival curves, the following steps were performed: 1. The Kaplan– Meier Plotter database was accessed. 2. The “Start KM Plotter for Immunotherapy” option was selected. 3. The P2RY2 gene name was entered into the query field. 4. Automatic cutoff values were applied for stratification. 5. Sample acquisition: “Pretreatment” was selected. Hazard ratios (HRs), 95% confidence intervals (CIs), and log-rank P values were calculated and displayed on the website. P values < 0.05 were considered statistically significant.

#### Statistics

All in vitro experiments were biologically repeated independently three to four times, yielding consistent results. Statistical analyses were conducted using GraphPad Prism (v10; GraphPad Software). Data points represent biological replicates and are expressed as mean ± standard deviation (s.d.). The statistical tests used to calculate P values are detailed in the figure legends. Biological replicates were included to ensure reproducibility, and the number of independent replicates for each experiment is specified in the figure legends.

## SUPPLEMENTAL INFORMATION

**- Document S1. Figures S1–S10**

Figure S1. eATP protects tumor cells from tumor-reactive TCR-T and CAR-T cells, related to Figure 1.

Figure S2. eATP suppresses T cell activation in coculture with tumor cells, but not in T cell monoculture, related to Figure 1.

Figure S3. P2RY2 expression in tumor cells suppresses antitumor T cell responses, related to Figure 2.

Figure S4. P2RY2 is a predominant driver of the COX-1/2-PGE₂ axis across tumor types, related to Figure 2.

Figure S5. P2RY2 targeting does not reduce intratumoral eATP levels, related to Figure 3.

Figure S6. Targeting P2RY2 remodels the TME and enhances anti-tumor T cell responses, related to Figure 4.

**- Table S1. The sequences of sgRNAs targeting P2X and P2Y purinergic receptors, related to Figures 1L and M.**

## REFERENCES

1. Greiner, J.V., and Glonek, T. (2021). Intracellular ATP Concentration and Implication for Cellular Evolution. Biology (Basel) 10. 10.3390/biology10111166.

2. Pellegatti, P., Raffaghello, L., Bianchi, G., Piccardi, F., Pistoia, V., and Di Virgilio, F. (2008). Increased level of extracellular ATP at tumor sites: in vivo imaging with plasma membrane luciferase. PLoS One 3, e2599. 10.1371/journal.pone.0002599.

3. Di Virgilio, F., Sarti, A.C., Falzoni, S., De Marchi, E., and Adinolfi, E. (2018). Extracellular ATP and P2 purinergic signalling in the tumour microenvironment. Nat Rev Cancer 18, 601–618. 10.1038/s41568-018-0037-0.

4. Kroemer, G., Galluzzi, L., Kepp, O., and Zitvogel, L. (2013). Immunogenic cell death in cancer therapy. Annu Rev Immunol 31, 51–72. 10.1146/annurev-immunol-032712-100008.

5. Di Virgilio, F., and Adinolfi, E. (2017). Extracellular purines, purinergic receptors and tumor growth. Oncogene 36, 293–303. 10.1038/onc.2016.206.

6. Grygorczyk, R., Boudreault, F., Ponomarchuk, O., Tan, J.J., Furuya, K., Goldgewicht, J., Kenfack, F.D., and Yu, F. (2021). Lytic Release of Cellular ATP: Physiological Relevance and Therapeutic Applications. Life (Basel) 11. 10.3390/life11070700.

7. Michaud, M., Martins, I., Sukkurwala, A.Q., Adjemian, S., Ma, Y., Pellegatti, P., Shen, S., Kepp, O., Scoazec, M., Mignot, G., et al. (2011). Autophagy-dependent anticancer immune responses induced by chemotherapeutic agents in mice. Science 334, 1573–1577. 10.1126/science.1208347.

8. Ghiringhelli, F., Apetoh, L., Tesniere, A., Aymeric, L., Ma, Y., Ortiz, C., Vermaelen, K., Panaretakis, T., Mignot, G., Ullrich, E., et al. (2009). Activation of the NLRP3 inflammasome in dendritic cells induces IL-1beta-dependent adaptive immunity against tumors. Nat Med 15, 1170–1178. 10.1038/nm.2028.

9. Martins, I., Tesniere, A., Kepp, O., Michaud, M., Schlemmer, F., Senovilla, L., Seror, C., Metivier, D., Perfettini, J.L., Zitvogel, L., and Kroemer, G. (2009). Chemotherapy induces ATP release from tumor cells. Cell Cycle 8, 3723–3728. 10.4161/cc.8.22.10026.

10. Zanoni, M., Sarti, A.C., Zamagni, A., Cortesi, M., Pignatta, S., Arienti, C., Tebaldi, M., Sarnelli, A., Romeo, A., Bartolini, D., et al. (2022). Irradiation causes senescence, ATP release, and P2X7 receptor isoform switch in glioblastoma. Cell Death Dis 13, 80. 10.1038/s41419-022-04526-0.

11. Ohshima, Y., Tsukimoto, M., Takenouchi, T., Harada, H., Suzuki, A., Sato, M., Kitani, H., and Kojima, S. (2010). gamma-Irradiation induces P2X(7) receptor-dependent ATP release from B16 melanoma cells. Biochim Biophys Acta 1800, 40–46. 10.1016/j.bbagen.2009.10.008.

12. Kepp, O., Bezu, L., Yamazaki, T., Di Virgilio, F., Smyth, M.J., Kroemer, G., and Galluzzi, L. (2021). ATP and cancer immunosurveillance. EMBO J 40, e108130. 10.15252/embj.2021108130.

13. Wilkin, F., Duhant, X., Bruyns, C., Suarez-Huerta, N., Boeynaems, J.M., and Robaye, B. (2001). The P2Y11 receptor mediates the ATP-induced maturation of human monocyte-derived dendritic cells. J Immunol 166, 7172–7177. 10.4049/jimmunol.166.12.7172.

14. Ferrari, D., La Sala, A., Chiozzi, P., Morelli, A., Falzoni, S., Girolomoni, G., Idzko, M., Dichmann, S., Norgauer, J., and Di Virgilio, F. (2000). The P2 purinergic receptors of human dendritic cells: identification and coupling to cytokine release. FASEB J 14, 2466–2476. 10.1096/fj.00-0031com.

15. Aymeric, L., Apetoh, L., Ghiringhelli, F., Tesniere, A., Martins, I., Kroemer, G., Smyth, M.J., and Zitvogel, L. (2010). Tumor cell death and ATP release prime dendritic cells and efficient anticancer immunity. Cancer Res 70, 855–858. 10.1158/0008-5472.CAN-09-3566.

16. Chen, Y., Corriden, R., Inoue, Y., Yip, L., Hashiguchi, N., Zinkernagel, A., Nizet, V., Insel, P.A., and Junger, W.G. (2006). ATP release guides neutrophil chemotaxis via P2Y2 and A3 receptors. Science 314, 1792–1795. 10.1126/science.1132559.

17. Brock, V.J., Wolf, I.M.A., Er-Lukowiak, M., Lory, N., Stahler, T., Woelk, L.M., Mittrucker, H.W., Muller, C.E., Koch-Nolte, F., Rissiek, B., et al. (2022). P2X4 and P2X7 are essential players in basal T cell activity and Ca(2+) signaling milliseconds after T cell activation. Sci Adv 8, eabl9770. 10.1126/sciadv.abl9770.

18. Loomis, W.H., Namiki, S., Ostrom, R.S., Insel, P.A., and Junger, W.G. (2003). Hypertonic stress increases T cell interleukin-2 expression through a mechanism that involves ATP release, P2 receptor, and p38 MAPK activation. J Biol Chem 278, 4590–4596. 10.1074/jbc.M207868200.

19. Schenk, U., Westendorf, A.M., Radaelli, E., Casati, A., Ferro, M., Fumagalli, M., Verderio, C., Buer, J., Scanziani, E., and Grassi, F. (2008). Purinergic control of T cell activation by ATP released through pannexin-1 hemichannels. Sci Signal 1, ra6. 10.1126/scisignal.1160583.

20. Yip, L., Woehrle, T., Corriden, R., Hirsh, M., Chen, Y., Inoue, Y., Ferrari, V., Insel, P.A., and Junger, W.G. (2009). Autocrine regulation of T-cell activation by ATP release and P2X7 receptors. FASEB J 23, 1685–1693. 10.1096/fj.08-126458.

21. Ledderose, C., Liu, K., Kondo, Y., Slubowski, C.J., Dertnig, T., Denicolo, S., Arbab, M., Hubner, J., Konrad, K., Fakhari, M., et al. (2018). Purinergic P2X4 receptors and mitochondrial ATP production regulate T cell migration. J Clin Invest 128, 3583–3594. 10.1172/JCI120972.

22. Romagnani, A., Rottoli, E., Mazza, E.M.C., Rezzonico-Jost, T., De Ponte Conti, B., Proietti, M., Perotti, M., Civanelli, E., Perruzza, L., Catapano, A.L., et al. (2020). P2X7 Receptor Activity Limits Accumulation of T Cells within Tumors. Cancer Res 80, 3906–3919. 10.1158/0008-5472.CAN-19-3807.

23. Bianchi, G., Vuerich, M., Pellegatti, P., Marimpietri, D., Emionite, L., Marigo, I., Bronte, V., Di Virgilio, F., Pistoia, V., and Raffaghello, L. (2014). ATP/P2X7 axis modulates myeloid-derived suppressor cell functions in neuroblastoma microenvironment. Cell Death Dis 5, e1135. 10.1038/cddis.2014.109.

24. Coutinho-Silva, R., Persechini, P.M., Bisaggio, R.D., Perfettini, J.L., Neto, A.C., Kanellopoulos, J.M., Motta-Ly, I., Dautry-Varsat, A., and Ojcius, D.M. (1999). P2Z/P2X7 receptor-dependent apoptosis of dendritic cells. Am J Physiol 276, C1139–1147. 10.1152/ajpcell.1999.276.5.C1139.

25. Surprenant, A., Rassendren, F., Kawashima, E., North, R.A., and Buell, G. (1996). The cytolytic P2Z receptor for extracellular ATP identified as a P2X receptor (P2X7). Science 272, 735–738. 10.1126/science.272.5262.735.

26. Bidula, S., Dhuna, K., Helliwell, R., and Stokes, L. (2019). Positive allosteric modulation of P2X7 promotes apoptotic cell death over lytic cell death responses in macrophages. Cell Death Dis 10, 882. 10.1038/s41419-019-2110-3.

27. Chvatchko, Y., Valera, S., Aubry, J.P., Renno, T., Buell, G., and Bonnefoy, J.Y. (1996). The involvement of an ATP-gated ion channel, P(2X1), in thymocyte apoptosis. Immunity 5, 275-283. 10.1016/s1074-7613(00)80322-2.

28. Vijayan, D., Young, A., Teng, M.W.L., and Smyth, M.J. (2017). Targeting immunosuppressive adenosine in cancer. Nat Rev Cancer 17, 709–724. 10.1038/nrc.2017.86.

29. Allard, B., Allard, D., Buisseret, L., and Stagg, J. (2020). The adenosine pathway in immuno-oncology. Nat Rev Clin Oncol 17, 611–629. 10.1038/s41571-020-0382-2.

30. Di Virgilio, F. (2012). Purines, purinergic receptors, and cancer. Cancer Res 72, 5441–5447. 10.1158/0008-5472.CAN-12-1600.

31. Wang, D., and DuBois, R.N. (2016). The Role of Prostaglandin E(2) in Tumor-Associated Immunosuppression. Trends Mol Med 22, 1–3. 10.1016/j.molmed.2015.11.003.

32. DuBois, R.N., Radhika, A., Reddy, B.S., and Entingh, A.J. (1996). Increased cyclooxygenase-2 levels in carcinogen-induced rat colonic tumors. Gastroenterology 110, 1259–1262. 10.1053/gast.1996.v110.pm8613017.

33. Zelenay, S., van der Veen, A.G., Bottcher, J.P., Snelgrove, K.J., Rogers, N., Acton, S.E., Chakravarty, P., Girotti, M.R., Marais, R., Quezada, S.A., et al. (2015). Cyclooxygenase-Dependent Tumor Growth through Evasion of Immunity. Cell 162, 1257–1270. 10.1016/j.cell.2015.08.015.

34. Pelly, V.S., Moeini, A., Roelofsen, L.M., Bonavita, E., Bell, C.R., Hutton, C., Blanco-Gomez, A., Banyard, A., Bromley, C.P., Flanagan, E., et al. (2021). Anti-Inflammatory Drugs Remodel the Tumor Immune Environment to Enhance Immune Checkpoint Blockade Efficacy. Cancer Discov 11, 2602–2619. 10.1158/2159-8290.CD-20-1815.

35. Lacher, S.B., Dorr, J., de Almeida, G.P., Honninger, J., Bayerl, F., Hirschberger, A., Pedde, A.M., Meiser, P., Ramsauer, L., Rudolph, T.J., et al. (2024). PGE(2) limits effector expansion of tumour-infiltrating stem-like CD8(+) T cells. Nature 629, 417–425. 10.1038/s41586-024-07254-x.

36. Morotti, M., Grimm, A.J., Hope, H.C., Arnaud, M., Desbuisson, M., Rayroux, N., Barras, D., Masid, M., Murgues, B., Chap, B.S., et al. (2024). PGE(2) inhibits TIL expansion by disrupting IL-2 signalling and mitochondrial function. Nature 629, 426–434. 10.1038/s41586-024-07352-w.

37. Elewaut, A., Estivill, G., Bayerl, F., Castillon, L., Novatchkova, M., Pottendorfer, E., Hoffmann-Haas, L., Schonlein, M., Nguyen, T.V., Lauss, M., et al. (2025). Cancer cells impair monocyte-mediated T cell stimulation to evade immunity. Nature 637, 716–725. 10.1038/s41586-024-08257-4.

38. Bayerl, F., Meiser, P., Donakonda, S., Hirschberger, A., Lacher, S.B., Pedde, A.M., Hermann, C.D., Elewaut, A., Knolle, M., Ramsauer, L., et al. (2023). Tumor-derived prostaglandin E2 programs cDC1 dysfunction to impair intratumoral orchestration of anti-cancer T cell responses. Immunity 56, 1341–1358 e1311. 10.1016/j.immuni.2023.05.011.

39. Lu, W., Yu, W., He, J., Liu, W., Yang, J., Lin, X., Zhang, Y., Wang, X., Jiang, W., Luo, J., et al. (2021). Reprogramming immunosuppressive myeloid cells facilitates immunotherapy for colorectal cancer. EMBO Mol Med 13, e12798. 10.15252/emmm.202012798.

40. Bi, C., Fu, Y., Zhang, Z., and Li, B. (2020). Prostaglandin E2 confers protection against diabetic coronary atherosclerosis by stimulating M2 macrophage polarization via the activation of the CREB/BDNF/TrkB signaling pathway. FASEB J 34, 7360–7371. 10.1096/fj.201902055R.

41. Luan, B., Yoon, Y.S., Le Lay, J., Kaestner, K.H., Hedrick, S., and Montminy, M. (2015). CREB pathway links PGE2 signaling with macrophage polarization. Proc Natl Acad Sci U S A 112, 15642–15647. 10.1073/pnas.1519644112.

42. Obermajer, N., Muthuswamy, R., Odunsi, K., Edwards, R.P., and Kalinski, P. (2011). PGE(2)-induced CXCL12 production and CXCR4 expression controls the accumulation of human MDSCs in ovarian cancer environment. Cancer Res 71, 7463–7470. 10.1158/0008-5472.CAN-11-2449.

43. Obermajer, N., Muthuswamy, R., Lesnock, J., Edwards, R.P., and Kalinski, P. (2011). Positive feedback between PGE2 and COX2 redirects the differentiation of human dendritic cells toward stable myeloid-derived suppressor cells. Blood 118, 5498–5505. 10.1182/blood-2011-07-365825.

44. Sinha, P., Clements, V.K., Fulton, A.M., and Ostrand-Rosenberg, S. (2007). Prostaglandin E2 promotes tumor progression by inducing myeloid-derived suppressor cells. Cancer Res 67, 4507–4513. 10.1158/0008-5472.CAN-06-4174.

45. Thumkeo, D., Punyawatthananukool, S., Prasongtanakij, S., Matsuura, R., Arima, K., Nie, H., Yamamoto, R., Aoyama, N., Hamaguchi, H., Sugahara, S., et al. (2022). PGE(2)-EP2/EP4 signaling elicits immunosuppression by driving the mregDC-Treg axis in inflammatory tumor microenvironment. Cell Rep 39, 110914. 10.1016/j.celrep.2022.110914.

46. Bonavita, E., Bromley, C.P., Jonsson, G., Pelly, V.S., Sahoo, S., Walwyn-Brown, K., Mensurado, S., Moeini, A., Flanagan, E., Bell, C.R., et al. (2020). Antagonistic Inflammatory Phenotypes Dictate Tumor Fate and Response to Immune Checkpoint Blockade. Immunity 53, 1215–1229 e1218. 10.1016/j.immuni.2020.10.020.

47. Holt, D., Ma, X., Kundu, N., and Fulton, A. (2011). Prostaglandin E(2) (PGE (2)) suppresses natural killer cell function primarily through the PGE(2) receptor EP4. Cancer Immunol Immunother 60, 1577–1586. 10.1007/s00262-011-1064-9.

48. Wei, J., Zhang, J., Wang, D., Cen, B., Lang, J.D., and DuBois, R.N. (2022). The COX-2-PGE2 Pathway Promotes Tumor Evasion in Colorectal Adenomas. Cancer Prev Res (Phila) 15, 285–296. 10.1158/1940-6207.CAPR-21-0572.

49. Chouaib, S., Robb, R.J., Welte, K., and Dupont, B. (1987). Analysis of prostaglandin E2 effect on T lymphocyte activation. Abrogation of prostaglandin E2 inhibitory effect by the tumor promotor 12.0 tetradecanoyl phorbol-13 acetate. J Clin Invest 80, 333–340. 10.1172/JCI113077.

50. Wang, D., Cabalag, C.S., Clemons, N.J., and DuBois, R.N. (2021). Cyclooxygenases and Prostaglandins in Tumor Immunology and Microenvironment of Gastrointestinal Cancer. Gastroenterology 161, 1813–1829. 10.1053/j.gastro.2021.09.059.

51. Eliopoulos, A.G., Dumitru, C.D., Wang, C.C., Cho, J., and Tsichlis, P.N. (2002). Induction of COX-2 by LPS in macrophages is regulated by Tpl2-dependent CREB activation signals. EMBO J 21, 4831–4840. 10.1093/emboj/cdf478.

52. Nakao, S., Ogtata, Y., Shimizu, E., Yamazaki, M., Furuyama, S., and Sugiya, H. (2002). Tumor necrosis factor alpha (TNF-alpha)-induced prostaglandin E2 release is mediated by the activation of cyclooxygenase-2 (COX-2) transcription via NFkappaB in human gingival fibroblasts. Mol Cell Biochem 238, 11–18. 10.1023/a:1019927616000.

53. Choi, Y.A., Lee, D.J., Lim, H.K., Jeong, J.H., Sonn, J.K., Kang, S.S., and Baek, S.H. (2004). Interleukin-1beta stimulates matrix metalloproteinase-2 expression via a prostaglandin E2-dependent mechanism in human chondrocytes. Exp Mol Med 36, 226–232. 10.1038/emm.2004.31.

54. Marcet, B., Libert, F., Boeynaems, J.M., and Communi, D. (2007). Extracellular nucleotides induce COX-2 up-regulation and prostaglandin E2 production in human A549 alveolar type II epithelial cells. Eur J Pharmacol 566, 167–171. 10.1016/j.ejphar.2007.04.003.

55. Molina-Holgado, E., Ortiz, S., Molina-Holgado, F., and Guaza, C. (2000). Induction of COX-2 and PGE(2) biosynthesis by IL-1beta is mediated by PKC and mitogen-activated protein kinases in murine astrocytes. Br J Pharmacol 131, 152–159. 10.1038/sj.bjp.0703557.

56. Sun, R., Carlson, N.G., Hemmert, A.C., and Kishore, B.K. (2005). P2Y2 receptor-mediated release of prostaglandin E2 by IMCD is altered in hydrated and dehydrated rats: relevance to AVP-independent regulation of IMCD function. Am J Physiol Renal Physiol 289, F585–592. 10.1152/ajprenal.00050.2005.

57. Welch, B.D., Carlson, N.G., Shi, H., Myatt, L., and Kishore, B.K. (2003). P2Y2 receptor-stimulated release of prostaglandin E2 by rat inner medullary collecting duct preparations. Am J Physiol Renal Physiol 285, F711–721. 10.1152/ajprenal.00096.2003.

58. Lee, T.H., Liu, P.S., Tsai, M.M., Chen, J.L., Wang, S.J., and Hsieh, H.L. (2020). The COX-2-derived PGE(2) autocrine contributes to bradykinin-induced matrix metalloproteinase-9 expression and astrocytic migration via STAT3 signaling. Cell Commun Signal 18, 185. 10.1186/s12964-020-00680-0.

59. Boumelha, J., de Castro, A., Bah, N., Cha, H., de Carne Trecesson, S., Rana, S., Tomaschko, M., Anastasiou, P., Mugarza, E., Moore, C., et al. (2024). CRISPR-Cas9 Screening Identifies KRAS-Induced COX2 as a Driver of Immunotherapy Resistance in Lung Cancer. Cancer Res 84, 2231–2246. 10.1158/0008-5472.CAN-23-2627.

60. Fang, L., Chang, H.M., Cheng, J.C., Leung, P.C., and Sun, Y.P. (2014). TGF-beta1 induces COX-2 expression and PGE2 production in human granulosa cells through Smad signaling pathways. J Clin Endocrinol Metab 99, E1217–1226. 10.1210/jc.2013-4100.

61. Fang, L., Cheng, J.C., Chang, H.M., Sun, Y.P., and Leung, P.C. (2013). EGF-like growth factors induce COX-2-derived PGE2 production through ERK1/2 in human granulosa cells. J Clin Endocrinol Metab 98, 4932–4941. 10.1210/jc.2013-2662.

62. Cho, W., and Choe, J. (2020). Prostaglandin E2 stimulates COX-2 expression via mitogen-activated protein kinase p38 but not ERK in human follicular dendritic cell-like cells. BMC Immunol 21, 20. 10.1186/s12865-020-00347-y.

63. Xu, X., Lu, Y., Cao, L., Miao, Y., Li, Y., Cui, Y., and Han, T. (2024). Tumor-intrinsic P2RY6 drives immunosuppression by enhancing PGE(2) production. Cell Rep 43, 114469. 10.1016/j.celrep.2024.114469.

64. Kanai, O., Ito, T., Saito, Z., Yamamoto, Y., Fujita, K., Okamura, M., Hashimoto, M., Nakatani, K., Sawai, S., and Mio, T. (2021). Effect of cyclooxygenase inhibitor use on immunotherapy efficacy in non-small cell lung cancer. Thorac Cancer 12, 949–957. 10.1111/1759-7714.13845.

65. Wang, D.Y., McQuade, J.L., Rai, R.R., Park, J.J., Zhao, S., Ye, F., Beckermann, K.E., Rubinstein, S.M., Johnpulle, R., Long, G.V., et al. (2020). The Impact of Nonsteroidal Anti-Inflammatory Drugs, Beta Blockers, and Metformin on the Efficacy of Anti-PD-1 Therapy in Advanced Melanoma. Oncologist 25, e602–e605. 10.1634/theoncologist.2019-0518.

66. Wang, S.J., Khullar, K., Kim, S., Yegya-Raman, N., Malhotra, J., Groisberg, R., Crayton, S.H., Silk, A.W., Nosher, J.L., Gentile, M.A., et al. (2020). Effect of cyclo-oxygenase inhibitor use during checkpoint blockade immunotherapy in patients with metastatic melanoma and non-small cell lung cancer. J Immunother Cancer 8. 10.1136/jitc-2020-000889.

67. Quandt, Z., Jacob, S., Fadlullah, M.Z.H., Wu, C., Wu, C., Huppert, L., Levine, L.S., Sison, P., Tsai, K.K., Chow, M., et al. (2024). Phase II trial of pembrolizumab, ipilimumab, and aspirin in melanoma: clinical outcomes and translational predictors of response. BJC Reports 2, 46. 10.1038/s44276-024-00057-7.

68. Iwasa, S., Koyama, T., Nishino, M., Kondo, S., Sudo, K., Yonemori, K., Yoshida, T., Tamura, K., Shimizu, T., Fujiwara, Y., et al. (2023). First-in-human study of ONO-4578, an antagonist of prostaglandin E(2) receptor 4, alone and with nivolumab in solid tumors. Cancer Sci 114, 211–220. 10.1111/cas.15574.

69. Mukherjee, D., Nissen, S.E., and Topol, E.J. (2001). Risk of cardiovascular events associated with selective COX-2 inhibitors. JAMA 286, 954–959. 10.1001/jama.286.8.954.

70. Kearney, P.M., Baigent, C., Godwin, J., Halls, H., Emberson, J.R., and Patrono, C. (2006). Do selective cyclo-oxygenase-2 inhibitors and traditional non-steroidal anti-inflammatory drugs increase the risk of atherothrombosis? Meta-analysis of randomised trials. BMJ 332, 1302–1308. 10.1136/bmj.332.7553.1302.

71. Bresalier, R.S., Sandler, R.S., Quan, H., Bolognese, J.A., Oxenius, B., Horgan, K., Lines, C., Riddell, R., Morton, D., Lanas, A., et al. (2005). Cardiovascular events associated with rofecoxib in a colorectal adenoma chemoprevention trial. N Engl J Med 352, 1092–1102. 10.1056/NEJMoa050493.

72. Henry, D., Dobson, A., and Turner, C. (1993). Variability in the risk of major gastrointestinal complications from nonaspirin nonsteroidal anti-inflammatory drugs. Gastroenterology 105, 1078–1088. 10.1016/0016-5085(93)90952-9.

73. Carson, J.L., Strom, B.L., Soper, K.A., West, S.L., and Morse, M.L. (1987). The Association of Nonsteroidal Anti-inflammatory Drugs With Upper Gastrointestinal Tract Bleeding. Archives of Internal Medicine 147, 85–88. 10.1001/archinte.1987.00370010087021.

74. Konturek, S.J., Kwiecien, N., Obtulowicz, W., Polanski, M., Kopp, B., and Oleksy, J. (1983). Comparison of prostaglandin E2 and ranitidine in prevention of gastric bleeding by aspirin in man. Gut 24, 89–93. 10.1136/gut.24.2.89.

75. Zhang, X., Donnan, P.T., Bell, S., and Guthrie, B. (2017). Non-steroidal anti-inflammatory drug induced acute kidney injury in the community dwelling general population and people with chronic kidney disease: systematic review and meta-analysis. BMC Nephrol 18, 256. 10.1186/s12882-017-0673-8.

76. Winkelmayer, W.C., Waikar, S.S., Mogun, H., and Solomon, D.H. (2008). Nonselective and cyclooxygenase-2-selective NSAIDs and acute kidney injury. Am J Med 121, 1092–1098. 10.1016/j.amjmed.2008.06.035.

77. Zhang, J., Ding, E.L., and Song, Y. (2006). Adverse effects of cyclooxygenase 2 inhibitors on renal and arrhythmia events: meta-analysis of randomized trials. JAMA 296, 1619–1632. 10.1001/jama.296.13.jrv60015.

78. Allard, B., Longhi, M.S., Robson, S.C., and Stagg, J. (2017). The ectonucleotidases CD39 and CD73: Novel checkpoint inhibitor targets. Immunol Rev 276, 121–144. 10.1111/imr.12528.

79. Vultaggio-Poma, V., Sarti, A.C., and Di Virgilio, F. (2020). Extracellular ATP: A Feasible Target for Cancer Therapy. Cells 9. 10.3390/cells9112496.

80. Wilson, D.J., and DuBois, R.N. (2022). Role of Prostaglandin E2 in the Progression of Gastrointestinal Cancer. Cancer Prev Res (Phila) 15, 355–363. 10.1158/1940-6207.CAPR-22-0038.

81. Im, S.J., Hashimoto, M., Gerner, M.Y., Lee, J., Kissick, H.T., Burger, M.C., Shan, Q., Hale, J.S., Lee, J., Nasti, T.H., et al. (2016). Defining CD8+ T cells that provide the proliferative burst after PD-1 therapy. Nature 537, 417–421. 10.1038/nature19330.

82. Utzschneider, D.T., Charmoy, M., Chennupati, V., Pousse, L., Ferreira, D.P., Calderon-Copete, S., Danilo, M., Alfei, F., Hofmann, M., Wieland, D., et al. (2016). T Cell Factor 1-Expressing Memory-like CD8(+) T Cells Sustain the Immune Response to Chronic Viral Infections. Immunity 45, 415–427. 10.1016/j.immuni.2016.07.021.

83. Wu, T., Ji, Y., Moseman, E.A., Xu, H.C., Manglani, M., Kirby, M., Anderson, S.M., Handon, R., Kenyon, E., Elkahloun, A., et al. (2016). The TCF1-Bcl6 axis counteracts type I interferon to repress exhaustion and maintain T cell stemness. Sci Immunol 1. 10.1126/sciimmunol.aai8593.

84. Huang, A.C., Postow, M.A., Orlowski, R.J., Mick, R., Bengsch, B., Manne, S., Xu, W., Harmon, S., Giles, J.R., Wenz, B., et al. (2017). T-cell invigoration to tumour burden ratio associated with anti-PD-1 response. Nature 545, 60–65. 10.1038/nature22079.

85. Kamphorst, A.O., Pillai, R.N., Yang, S., Nasti, T.H., Akondy, R.S., Wieland, A., Sica, G.L., Yu, K., Koenig, L., Patel, N.T., et al. (2017). Proliferation of PD-1+ CD8 T cells in peripheral blood after PD-1-targeted therapy in lung cancer patients. Proc Natl Acad Sci U S A 114, 4993–4998. 10.1073/pnas.1705327114.

86. Knight, R., Kilpatrick, L.E., Hill, S.J., and Stocks, M.J. (2024). Design, Synthesis, and Evaluation of a New Chemotype Fluorescent Ligand for the P2Y2 Receptor. ACS Medicinal Chemistry Letters 15, 1127–1135. 10.1021/acsmedchemlett.4c00211.

87. Jacobson, K.A., Ivanov, A.A., de Castro, S., Harden, T.K., and Ko, H. (2009). Development of selective agonists and antagonists of P2Y receptors. Purinergic Signal 5, 75–89. 10.1007/s11302-008-9106-2.

88. Jasmer, K.J., Munoz Forti, K., Woods, L.T., Cha, S., and Weisman, G.A. (2023). Therapeutic potential for P2Y(2) receptor antagonism. Purinergic Signal 19, 401–420. 10.1007/s11302-022-09900-3.

89. Homolya, L., Watt, W.C., Lazarowski, E.R., Koller, B.H., and Boucher, R.C. (1999). Nucleotide-regulated calcium signaling in lung fibroblasts and epithelial cells from normal and P2Y(2) receptor (-/-) mice. J Biol Chem 274, 26454–26460. 10.1074/jbc.274.37.26454.

90. He, X., Wan, J., Yang, X., Zhang, X., Huang, D., Li, X., Zou, Y., Chen, C., Yu, Z., Xie, L., et al. (2021). Bone marrow niche ATP levels determine leukemia-initiating cell activity via P2X7 in leukemic models. J Clin Invest 131. 10.1172/JCI140242.

91. Jorritsma, A., Gomez-Eerland, R., Dokter, M., van de Kasteele, W., Zoet, Y.M., Doxiadis, II, Rufer, N., Romero, P., Morgan, R.A., Schumacher, T.N., and Haanen, J.B. (2007). Selecting highly affine and well-expressed TCRs for gene therapy of melanoma. Blood 110, 3564–3572. 10.1182/blood-2007-02-075010.

92. Matas-Nadal, C., Bech-Serra, J.J., Guasch-Valles, M., Fernandez-Armenteros, J.M., Barcelo, C., Casanova, J.M., de la Torre Gomez, C., Aguayo Ortiz, R., and Gari, E. (2020). Evaluation of Tumor Interstitial Fluid-Extraction Methods for Proteome Analysis: Comparison of Biopsy Elution versus Centrifugation. J Proteome Res 19, 2598–2605. 10.1021/acs.jproteome.9b00770.

93. Kohler, G., and Milstein, C. (1975). Continuous cultures of fused cells secreting antibody of predefined specificity. Nature 256, 495–497. 10.1038/256495a0.

94. Fujimoto, A., Kosaka, N., Hasegawa, H., Suzuki, H., Sugano, S., and Chiba, J. (2012). Enhancement of antibody responses to native G protein-coupled receptors using E. coli GroEL as a molecular adjuvant in DNA immunization. J Immunol Methods 375, 243–251. 10.1016/j.jim.2011.11.007.

95. A. Jorritsma et al., Selecting highly affine and well-expressed TCRs for gene therapy of melanoma. Blood 110, 3564–3572 (2007).

96. C. Matas-Nadal et al., Evaluation of Tumor Interstitial Fluid-Extraction Methods for Proteome Analysis: Comparison of Biopsy Elution versus Centrifugation. J Proteome Res 19, 2598–2605 (2020).

97. G. Kohler, C. Milstein, Continuous cultures of fused cells secreting antibody of predefined specificity. Nature 256, 495–497 (1975).

98. A. Fujimoto et al., Enhancement of antibody responses to native G protein-coupled receptors using E. coli GroEL as a molecular adjuvant in DNA immunization. J Immunol Methods 375, 243–251 (2012).

