## Supplemental data for "P2RY2 is a purinergic immune checkpoint linking extracellular ATP to immune evasion and adaptive resistance to immunotherapy"

1 Division Immune Regulation in Cancer, German Cancer Research Center (DKFZ) Heidelberg; Heidelberg, 69120, Germany.
2 Faculty of Biosciences, Heidelberg University; Heidelberg, 69120, Germany.
3 Faculty of Medicine, Heidelberg University; Heidelberg, 69120, Germany.
4 Division of Experimental Neurosurgery, Department of Neurosurgery, Heidelberg University Hospital; Heidelberg, 69120, Germany.
5 DKFZ-Bayer Joint Immunotherapeutics Laboratory, German Cancer Research Center; Heidelberg, 69120, Germany.
6 Department of Immunology, Leiden University Medical Center (LUMC); Leiden, 2333, the Netherlands.
7 Core Facility Antibodies, German Cancer Research Center; Heidelberg, 69120, Germany.
8 German Cancer Consortium (DKTK), National Center for Tumor Diseases (NCT); Heidelberg, 69120, Germany.
9 Lead contact

**Figure S1**

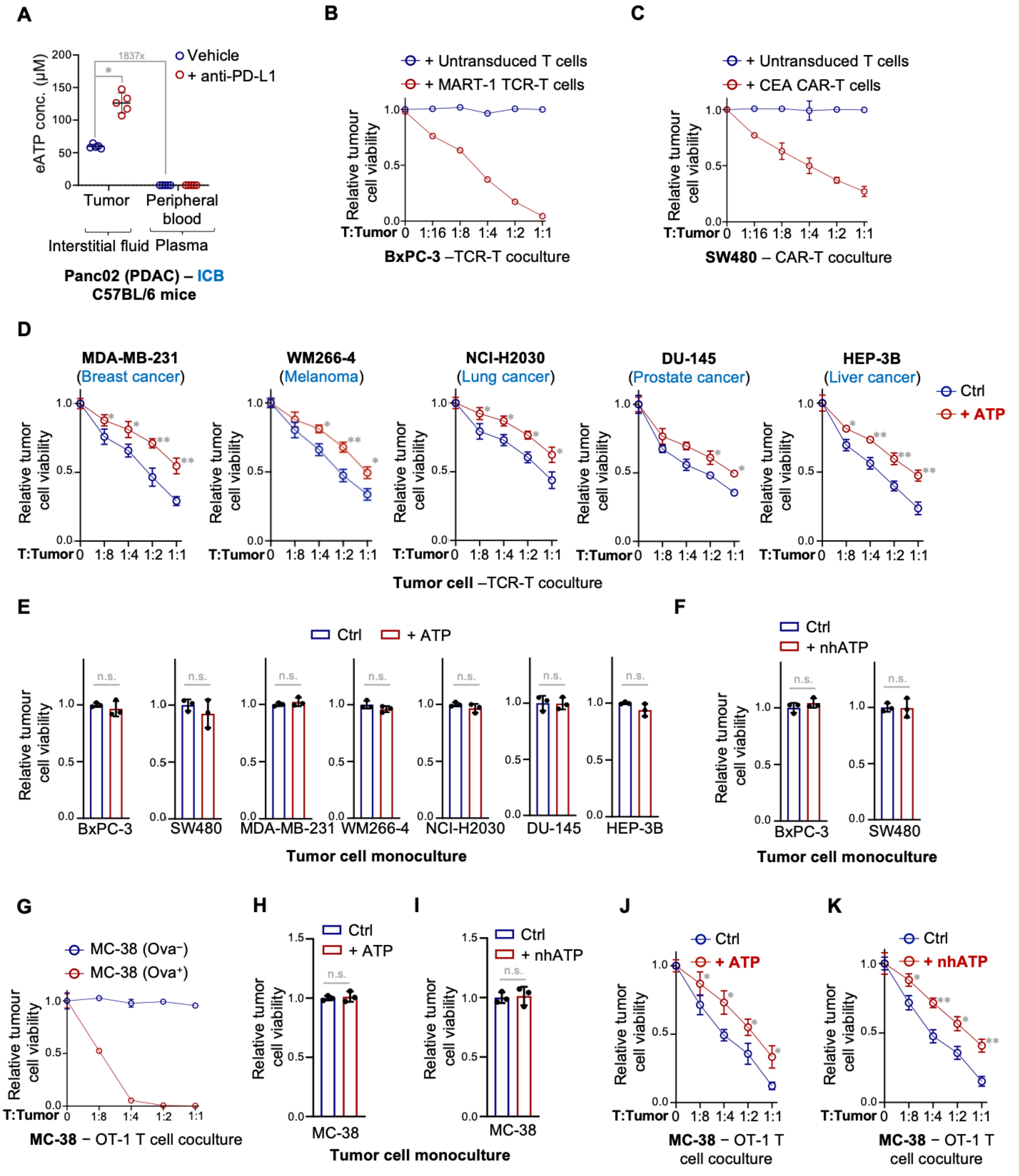


**Figure S1. eATP protects tumor cells from tumor-reactive TCR-T and CAR-T cells, related to Figure 1.**

(**A**) eATP concentrations are markedly elevated in the TME and rise further following ICB treatment. Syngeneic mouse PDAC Panc02 cells were subcutaneously implanted into immunocompetent C57BL/6 mice. After tumor establishment, mice were treated with either vehicle control (Vehicle) or a PD-L1 blocking antibody (anti–PD-L1). Interstitial fluid from tumors and peripheral blood plasma were collected from both immunotherapy-treated and vehicle control groups. eATP levels were measured using the CellTiter-Glo Cell Viability assay. Fold changes are indicated.

(**B**) MART-1 TCR-T cells efficiently eliminates antigen-loaded BxPC-3 tumor cells in a dose-dependent manner. BxPC-3 cells were transduced with a fusion construct encoding the MART-1 epitope (ELAGIGILTV), HLA-A2, and β2-microglobulin (B2M) to enable recognition by MART-1 TCR–transduced T cells (clone 1D3). Tumor cells were cocultured with either MART-1 TCR-T cells or untransduced T cells at the indicated T cell:tumor cell ratios. Tumor cell viability was assessed using the CellTiter-Blue Cell Viability Assay.

(**C**) CEA CAR-T cells specifically target SW480 tumor cells in a dose-dependent manner. SW480 tumor cells, which constitutively express CEA, were cocultured with CEA CAR-T cells or untransduced T cells at the indicated T:Tumor ratios. Tumor cell viability was assessed using the CellTiter-Blue Cell Viability Assay.

(**D**) eATP protects diverse tumor types from tumor-reactive T cells. TCR–transduced T cells were cocultured with antigen-positive tumor cells from various cancer types: MDA-MB-231 (breast cancer), WM266-4 (melanoma), NCI-H2030 (lung cancer), DU-145 (prostate cancer), and HEP-3B (liver cancer), with or without 200 μM ATP at the indicated T cell:tumor cell ratios. Tumor cell viability was assessed after 72 hours using the CellTiter-Blue Cell Viability Assay. To enable antigen-specific T cell–tumor cell interactions, different strategies were applied based on the endogenous expression of MART-1 and HLA-A2 in each tumor cell line: MART-1 and HLA-A2 double-positive tumor cells (WM266-4) were directly cocultured with MART-1 TCR–transduced T cells. HLA-A2$^+$ but MART-1$^-$ tumor cells (MDA-MB-231) were preloaded with the MART-1 epitope (ELAGIGILTV) via lentiviral transduction. Tumor cells lacking both MART-1 and HLA-A2 (DU-145, HEP-3B, and NCI-H2030) were transduced with a fusion construct encoding the MART-1 epitope, HLA-A2, and β2-microglobulin (B2M) prior to coculture.

(**E**, **F**) eATP does not affect tumor cell proliferation in monoculture. Tumor cells, including BxPC-3, SW480, MDA-MB-231, WM266-4, NCI-H2030, DU-145, and HEP-3B (E), or BxPC-3 and SW480 (F), were cultured without T cells in the absence or presence of 200 μM ATP (E) or ATPγS (nhATP) (F) for 72 hours. Tumor cell viability was assessed using the CellTiter-Blue assay.

(**G**) OT-1 T cells selectively eliminate Ova-expressing MC-38 tumor cells. Wild-type (Ova$^-$) and Ova-overexpressing (Ova$^+$) MC-38 cells were cocultured with OT-1 T cells at indicated effector-to-target (T:Tumor) ratios. Tumor cell viability was assessed using the CellTiter-Blue Cell Viability assay.

(**H**, **I**) Extracellular ATP and nhATP do not affect MC-38 tumor cell proliferation in monoculture. MC-38 cells were cultured alone in the absence or presence of 200 μM ATP (H) or ATPγS (nhATP) (I) for 72 hours. Tumor cell viability was assessed using the CellTiter-Blue Cell Viability assay.

(**J**, **K**) Extracellular ATP and nhATP protect murine tumor cells from OT-1 T cell killing. MC-38 (Ova$^+$) cells were cocultured with OT-1 T cells at the indicated T cell:tumor cell ratios in the absence or presence of 200 μM ATP (J) or ATPγS (nhATP) (K) for 72 hours. Tumor cell viability was assessed using the CellTiter-Blue Cell Viability assay.

Data represent the mean ± standard deviation of biological replicates ($n \geq 3$). Sample sizes for panels (A) are: vehicle (n=5), anti-PD-L1 (n=5). p-values were determined using unpaired two-tailed Student's t-test (A, E, F, H, I), or two-way ANOVA with Šídák's multiple comparisons test (D, J, K). A p-value ≥ 0.05 indicates non-significance (n.s.), while a p-value < 0.05 is denoted as ∗, and a p-value < 0.0001 is represented as ∗∗.

### Figure S2

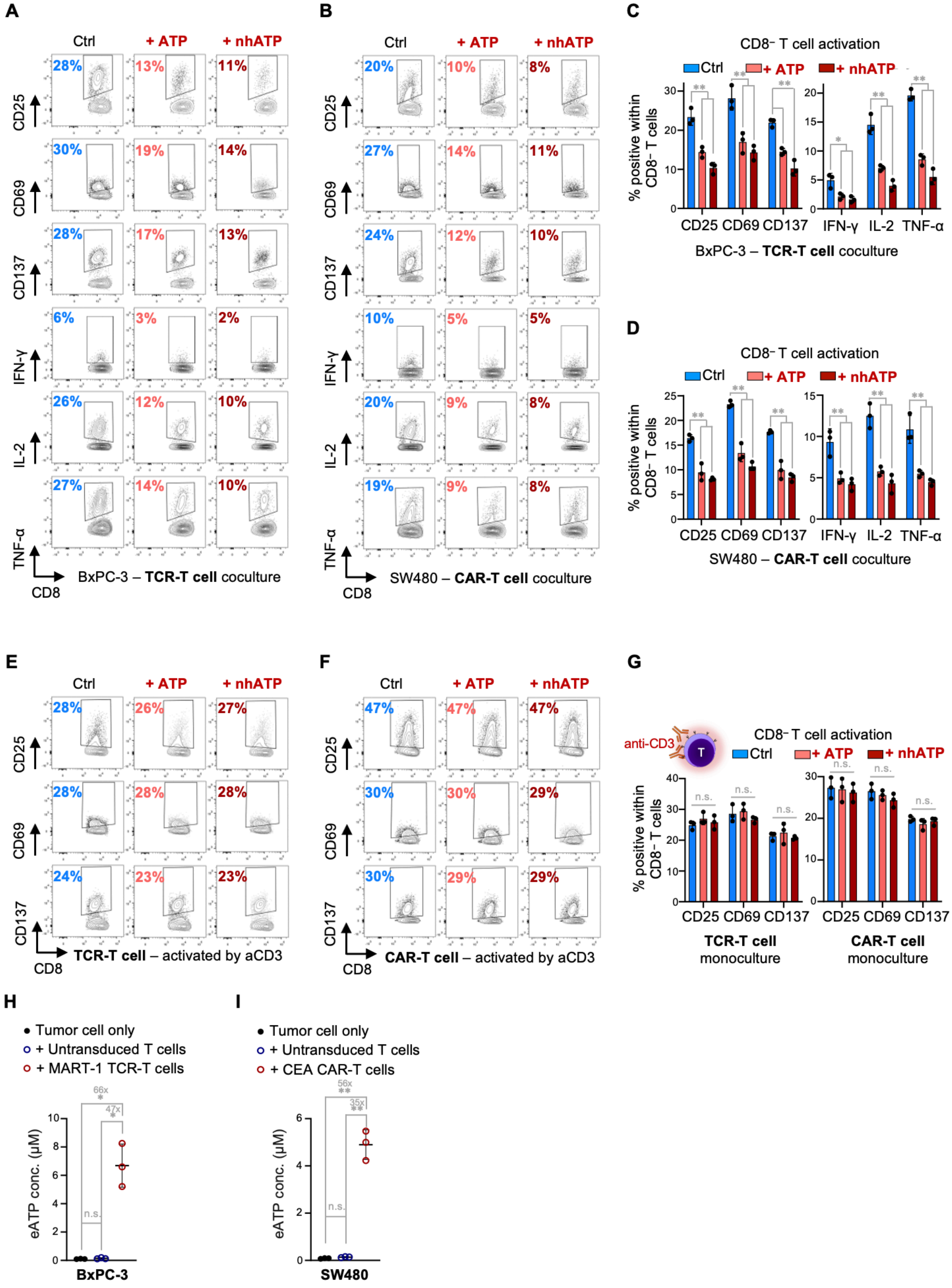
A
Ctrl
+ ATP
+ nhATP
CD25
CD69
CD137
IFN-γ
IL-2
TNF-α
CD8
BxPC-3 – TCR-T cell coculture
B
SW480 – CAR-T cell coculture
C
CD8⁻ T cell activation
% positive within CD8⁻ T cells
CD25 CD69 CD137 IFN-γ IL-2 TNF-α
BxPC-3 – TCR-T cell coculture
D
CD8⁻ T cell activation
SW480 – CAR-T cell coculture
E
TCR-T cell – activated by aCD3
F
CAR-T cell – activated by aCD3
G
anti-CD3
CD8⁻ T cell activation
n.s.
TCR-T cell monoculture
CAR-T cell monoculture
H
Tumor cell only
+ Untransduced T cells
+ MART-1 TCR-T cells
66x
47x
eATP conc. (µM)
BxPC-3
I
Tumor cell only
+ Untransduced T cells
+ CEA CAR-T cells
56x
35x
SW480

**Figure S2. eATP suppresses T cell activation in coculture with tumor cells, but not in T cell monoculture, related to Figure 1.**

(**A**, **B**) eATP suppresses $CD8^+$ T cell activation in coculture with tumor cells. MART-1 TCR–transduced T cells (A) or CEA CAR–transduced T cells (B) were cocultured with antigen-positive BxPC-3 or SW480 cells, respectively, for 24 hours in the absence or presence of 200 μM ATP or the non-hydrolyzable analog ATPγS (nhATP). $CD8^+$ T cell activation was assessed by flow cytometry. Representative contour plots are shown, with numbers indicating the percentage of $CD8^+$ T cells positive for activation markers (CD25, CD69, CD137) and cytokine production (IFN-γ, IL-2, TNF-α). Quantification of these results is presented in Fig. 1, I and J.

(**C, D**) eATP suppresses T cell activation in $CD8^-$ T cells cocultured with tumor cells. MART-1 TCR–transduced T cells (C) or CEA CAR–transduced T cells (D) were cocultured with antigen-positive BxPC-3 or SW480 cells, respectively, for 24 hours in the absence or presence of 200 μM ATP or the non-hydrolyzable analog ATPγS (nhATP). Activation of $CD8^-$ T cells was assessed by flow cytometry, measuring expression of CD25, CD69, CD137, IFN-γ, IL-2, and TNF-α. Note: The MART-1 TCR (1D3 clone) is CD8-independent. These data complement Fig. 1, I and J, which show the $CD8^+$ T cell responses.

(**E, F**) eATP does not affect $CD8^+$ T cell activation in T cell monoculture. MART-1 TCR-transduced T cells (E) or CEA CAR-transduced T cells (F) were stimulated with 0.1 μg/mL anti-CD3 antibody for 24 hours in the absence or presence of 200 μM ATP or the non-hydrolyzable analog ATPγS (nhATP). Representative contour plots are shown, with numbers indicating the percentage of $CD8^+$ T cells positive for activation markers (CD25, CD69, CD137). Quantification of these data is shown in Fig. 1K.

(**G**) eATP does not affect $CD8^-$ T cell activation in T cell monoculture. MART-1 TCR-transduced or CEA CAR-transduced T cells were stimulated with 0.1 μg/mL anti-CD3 antibody for 24 hours in the absence or presence of 200 μM ATP or the non-hydrolyzable analog ATPγS (nhATP). $CD8^-$ T cell activation was assessed by flow cytometry, measuring the percentage of cells positive for CD25, CD69, and CD137. Note: The MART-1 TCR (1D3 clone) is CD8-independent. These data complement Fig. 1K, which shows the $CD8^+$ T cell responses.

(**H, I**) eATP levels notably increase during antigen-specific T cell–tumor cell coculture. Antigen-positive BxPC-3 (H) or SW480 (I) tumor cells were cultured alone or cocultured with untransduced T cells, TCR-T cells, or CAR-T cells as indicated for 24 hours. eATP concentrations in the supernatant were measured using the CellTiter-Glo assay. Fold changes are indicated.

Data represent the mean ± standard deviation of biological replicates ($n \geq 3$). p-values were determined using one-way ANOVA with Dunnett's multiple comparisons test (C, D, G) or one-way ANOVA with Tukey's multiple comparisons test (H and I). A p-value ≥ 0.05 indicates non-significance (n.s.), while a p-value < 0.05 is denoted as ∗, and a p-value < 0.0001 is represented as ∗∗.

Figure S3

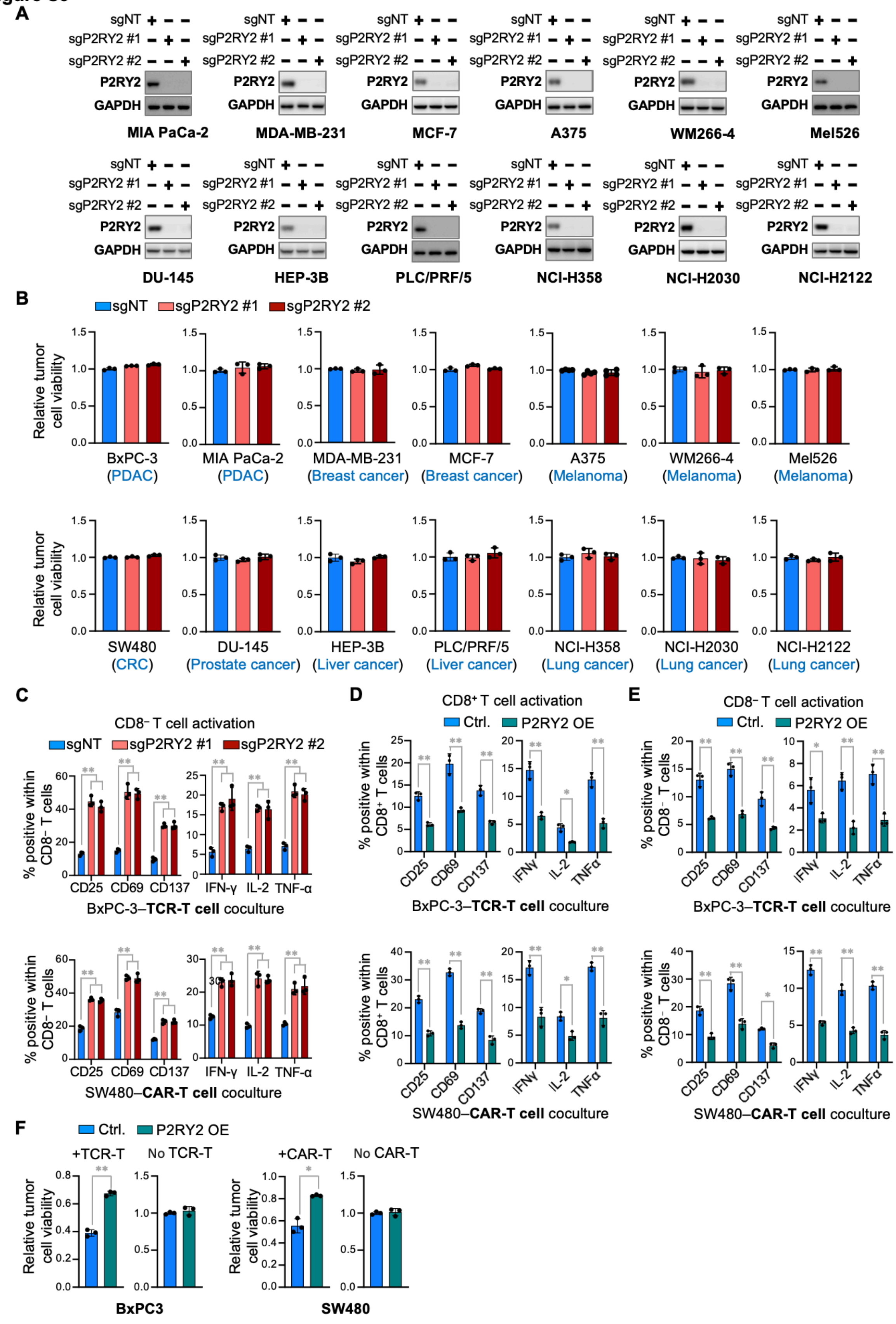

A
sgNT + − −
sgP2RY2 #1 − + −
sgP2RY2 #2 − − +
P2RY2
GAPDH
MIA PaCa-2
MDA-MB-231
MCF-7
A375
WM266-4
Mel526
DU-145
HEP-3B
PLC/PRF/5
NCI-H358
NCI-H2030
NCI-H2122
B
sgNT
sgP2RY2 #1
sgP2RY2 #2
Relative tumor cell viability
BxPC-3 (PDAC)
MIA PaCa-2 (PDAC)
MDA-MB-231 (Breast cancer)
MCF-7 (Breast cancer)
A375 (Melanoma)
WM266-4 (Melanoma)
Mel526 (Melanoma)
SW480 (CRC)
DU-145 (Prostate cancer)
HEP-3B (Liver cancer)
PLC/PRF/5 (Liver cancer)
NCI-H358 (Lung cancer)
NCI-H2030 (Lung cancer)
NCI-H2122 (Lung cancer)
C
CD8⁻ T cell activation
% positive within CD8⁻ T cells
CD25 CD69 CD137 IFN-γ IL-2 TNF-α
BxPC-3–TCR-T cell coculture
SW480–CAR-T cell coculture
D
CD8⁺ T cell activation
Ctrl.
P2RY2 OE
% positive within CD8⁺ T cells
CD25 CD69 CD137 IFNγ IL-2 TNFα
BxPC-3–TCR-T cell coculture
SW480–CAR-T cell coculture
E
CD8⁻ T cell activation
% positive within CD8⁻ T cells
BxPC-3–TCR-T cell coculture
SW480–CAR-T cell coculture
F
Ctrl.
P2RY2 OE
+TCR-T
No TCR-T
+CAR-T
No CAR-T
Relative tumor cell viability
BxPC3
SW480

**Figure S3. P2RY2 expression in tumor cells suppresses antitumor T cell responses, related to Figure 2.**

(**A**) Efficient CRISPR/Cas9-mediated knockout of P2RY2 in the panel of human tumor cell lines shown in Figure 2A. Western blot analysis of P2RY2 expression in Cas9-expressing tumors cells transduced with sgRNAs targeting P2RY2 (sgP2RY2 #1 and sgP2RY2 #2) or a non-targeting control sgRNA (sgNT). GAPDH served as a loading control.

(**B**) P2RY2 knockout does not affect the basal viability of tumor cells. P2RY2-proficient (sgNT) and P2RY2-deficient (sgP2RY2 #1, sgP2RY2 #2) tumor cells from various cancer types as indicated were cultured alone for 72 hours. Tumor cell viability was assessed using the CellTiter-Blue Cell Viability Assay.

(**C**) P2RY2 knockout in tumor cells enhances $CD8^{-}$ T cell activation. P2RY2-proficient (sgNT) or P2RY2-deficient (sgP2RY2 #1, sgP2RY2 #2) BxPC-3 and SW480 tumor cells were cocultured with tumor-reactive TCR-T cells or CAR-T cells for 24 hours. Flow cytometry analysis was performed to assess T cell activation, measuring CD137, CD69, and CD25 activation markers and cytokine production (IFN-γ, IL-2, TNF-α) in $CD3^{+}CD8^{-}$ T cells.

(**D**-**F**) P2RY2 overexpression in tumor cells suppresses T cell activation and confers resistance to antigen-specific T cell-mediated killing. P2RY2-overexpressing (P2RY2 OE) or control (Ctrl.) BxPC-3 and SW480 tumor cells were cocultured with tumor-reactive TCR-T and CAR-T cells. After 24 hours of coculture, flow cytometry analysis was performed to assess T cell activation by measuring CD25, CD69, and CD137 expression, as well as cytokine production (IFN-γ, IL-2, TNF-α) in $CD3^{+}CD8^{+}$ (D) and $CD3^{+}CD8^{-}$ (E) T cell populations. After 72 hours of coculture, tumor cell viability in the presence or absence of tumor reactive TCR-T or CAR-T cells was assessed using the CellTiter-Blue Cell Viability Assay (F).

Data represent the mean ± standard deviation of biological replicates ($n \geq 3$). p-values were determined using one-way ANOVA with Dunnett's multiple comparisons test (C) or unpaired two-tailed Student’s t-test (D-F). A p-value ≥ 0.05 indicates non-significance (n.s.), while a $p$-value $< 0.05$ is denoted as $*$, and a p-value $< 0.0001$ is represented as $**$.

### Figure S4

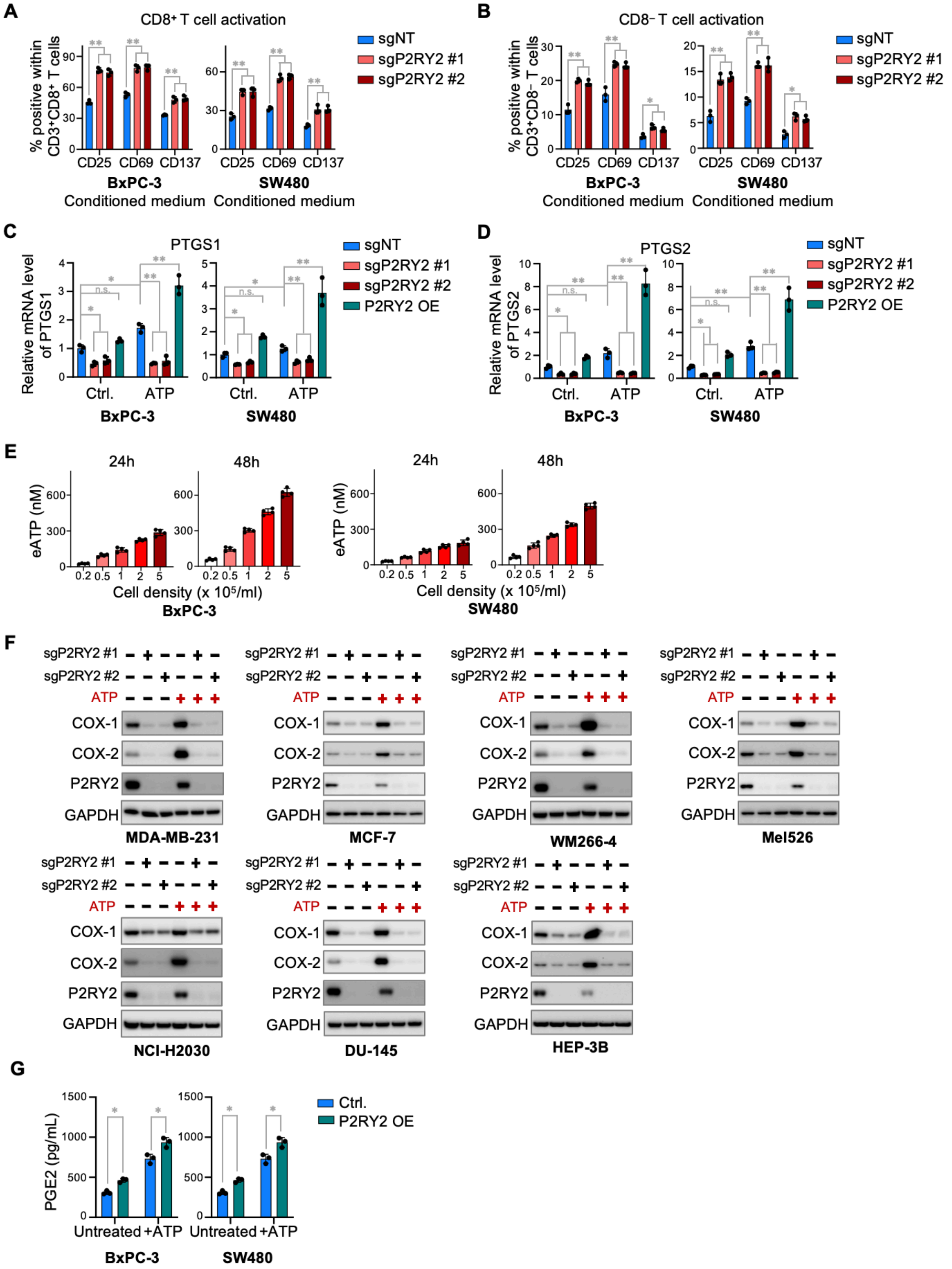
A
CD8⁺ T cell activation
% positive within CD3⁺CD8⁺ T cells
CD25 CD69 CD137
BxPC-3 Conditioned medium
SW480 Conditioned medium
sgNT
sgP2RY2 #1
sgP2RY2 #2
B
CD8⁻ T cell activation
% positive within CD3⁺CD8⁻ T cells
C
PTGS1
Relative mRNA level of PTGS1
Ctrl. ATP
BxPC-3
SW480
P2RY2 OE
D
PTGS2
Relative mRNA level of PTGS2
E
24h
48h
eATP (nM)
Cell density (x 10⁵/ml)
BxPC-3
SW480
F
sgP2RY2 #1
sgP2RY2 #2
ATP
COX-1
COX-2
P2RY2
GAPDH
MDA-MB-231
MCF-7
WM266-4
Mel526
NCI-H2030
DU-145
HEP-3B
G
PGE2 (pg/mL)
Untreated +ATP
Ctrl.
P2RY2 OE

**Figure S4. P2RY2 is a predominant driver of the COX-1/2-$PGE_2$ axis across tumor types, related to Figure 2.**

(**A, B**) Conditioned medium from P2RY2-proficient tumor cells suppresses T cell activation. Conditioned medium was collected from P2RY2-proficient (sgNT) and P2RY2-deficient (sgP2RY2 #1 and sgP2RY2 #2) BxPC-3 and SW480 tumor cells and used to culture T cells in the presence of anti-CD3 stimulation. T cell activation was assessed by flow cytometry, measuring CD25, CD69, and CD137 expression in $CD3^+CD8^+$ (A) and $CD3^+CD8^-$ (B) T cell populations.

(**C, D**) P2RY2 drives eATP-induced mRNA upregulation of *PTGS1* and *PTGS2*. Quantitative PCR (qPCR) analysis of *PTGS1* (COX-1) (C) and *PTGS2* (COX-2) (D) mRNA levels in P2RY2-proficient (sgNT), P2RY2-deficient (sgP2RY2 #1, sgP2RY2 #2), and P2RY2-overexpressing (P2RY2 OE) tumor cells under basal conditions (Ctrl.) or after treatment with 200 μM ATP (ATP) for 8 hours.

(**E**) eATP levels in cell culture supernatants. BxPC-3 and SW480 cells were seeded at the indicated densities and cultured for 24 or 48 hours. Supernatants were collected, and eATP concentrations were measured using CellTiter-Glo assay.

(**F**) P2RY2 is a predominant driver of COX-1 and COX-2 expression across tumor types. Western blot analysis of COX-1, COX-2, and P2RY2 expression in P2RY2-proficient (sgNT) and P2RY2-deficient (sgP2RY2 #1, sgP2RY2 #2) tumor cells with or without treatment by 200 μM exogenously supplemented ATP for 24 hours. GAPDH served as a loading control.

(**G**) P2RY2 overexpression enhances $PGE_2$ production by tumor cells. Culture medium conditioned by P2RY2-overexpressing (P2RY2 OE) and control (Ctrl.) tumor cells, either untreated or supplemented with 200 μM ATP for 24 hours, was collected. $PGE_2$ concentrations in the conditioned medium were quantified using ELISA.

Data represent the mean ± standard deviation of biological replicates ($n \geq 3$). P values were determined using one-way ANOVA with Dunnett's multiple comparisons test (A, B) or two-way ANOVA with Tukey's multiple comparisons test (C, D, G). A p-value ≥ 0.05 indicates non-significance (n.s.), while a p-value < 0.05 is denoted as ∗, and a p-value < 0.0001 is represented as ∗∗.

**Figure S5**

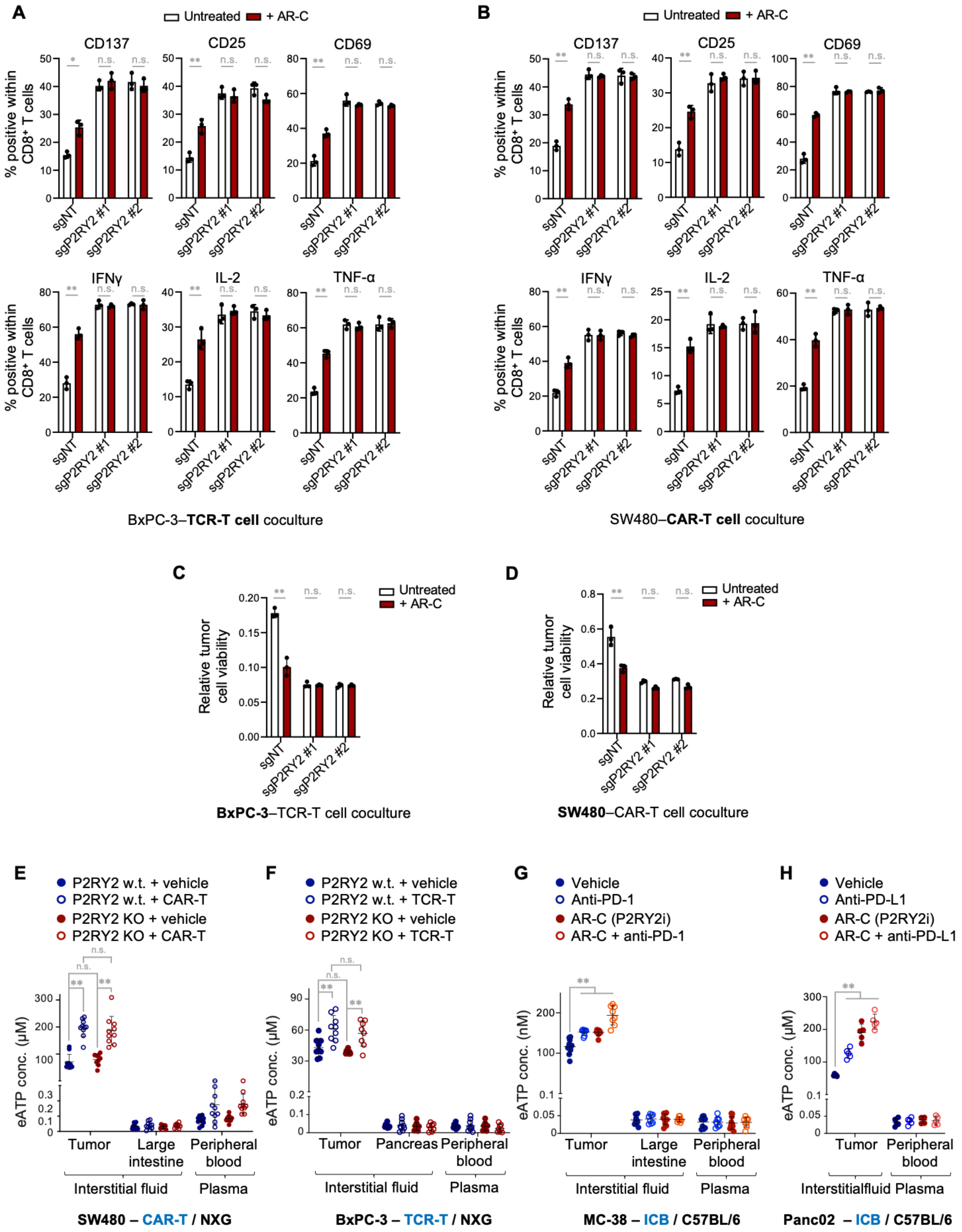
A
Untreated
+ AR-C
CD137
CD25
CD69
IFNγ
IL-2
TNF-α
% positive within CD8⁺ T cells
sgNT
sgP2RY2 #1
sgP2RY2 #2
n.s.
BxPC-3–TCR-T cell coculture
B
SW480–CAR-T cell coculture
C
Relative tumor cell viability
BxPC-3–TCR-T cell coculture
D
SW480–CAR-T cell coculture
E
P2RY2 w.t. + vehicle
P2RY2 w.t. + CAR-T
P2RY2 KO + vehicle
P2RY2 KO + CAR-T
eATP conc. (μM)
Tumor
Large intestine
Peripheral blood
Interstitial fluid
Plasma
SW480 – CAR-T / NXG
F
P2RY2 w.t. + vehicle
P2RY2 w.t. + TCR-T
P2RY2 KO + vehicle
P2RY2 KO + TCR-T
Pancreas
BxPC-3 – TCR-T / NXG
G
Vehicle
Anti-PD-1
AR-C (P2RY2i)
AR-C + anti-PD-1
eATP conc. (nM)
MC-38 – ICB / C57BL/6
H
Anti-PD-L1
AR-C + anti-PD-L1
Interstitialfluid
Panc02 – ICB / C57BL/6

**Figure S5. P2RY2 targeting does not reduce intratumoral eATP levels, related to Figure 3.**

**(A, B)** AR-C 118925XX (AR-C) enhances $CD8^+$ T cell activation only in the presence of P2RY2-proficient tumor cells. P2RY2-proficient (sgNT) or P2RY2-deficient (sgP2RY2 #1, #2) BxPC-3 and SW480 tumor cells were cocultured with tumor-reactive TCR-T or CAR-T cells for 24 hours in the absence or presence of the P2RY2 inhibitor AR-C. $CD8^+$ T cell activation was evaluated by flow cytometry, assessing expression of CD137, CD25, and CD69, and production of IFN-γ, IL-2, and TNF-α in $CD3^+CD8^+$ T cells.

**(C, D)** AR-C enhances T cell–mediated tumor cell killing in a P2RY2-dependent manner. Relative tumor cell viability was assessed after 72 hours of coculture using the CellTiter-Blue assay. AR-C enhanced T cell activation and tumor cell killing only in P2RY2-proficient tumor cells but had no additional effect in P2RY2-deficient cells, confirming that AR-C functions specifically through inhibition of tumoral P2RY2.

**(E–H)** eATP concentrations in the interstitial fluid of tumors and matched normal tissues, as well as peripheral blood plasma. Interstitial fluid was collected from all experimental and control groups in the SW480–CAR-T model (E), BxPC-3–TCR-T model (F), MC-38–anti–PD-1 model (G), and Panc02–anti–PD-L1 model (H), as described in Fig. 3A–D. eATP levels were measured using the CellTiter-Glo assay.

Data are presented as mean ± standard deviation of biological replicates ($n \geq 3$). For panel (E), the sample sizes are: P2RY2 wild-type (w.t.) + vehicle (n=10), P2RY2 knockout (KO) + vehicle (n=9), P2RY2 w.t. + CAR-T (n=10), and P2RY2 KO + CAR-T (n=10). For panel (F), the sample sizes are: P2RY2 wild-type (w.t.) + vehicle (n=9), P2RY2 knockout (KO) + vehicle (n=8), P2RY2 w.t. + TCR-T (n=9), and P2RY2 KO + TCR-T (n=8). For panel (G), the sample sizes are: vehicle (n=10), anti-PD-1 (n=8), AR-C (n=10), and AR-C + anti-PD-1 (n=8). For panel (H) are: vehicle (n=5), anti-PD-L1 (n=5), AR-C (n=5), and AR-C + anti-PD-L1 (n=5). Statistical significance was determined using two-way ANOVA with Šídák's multiple comparisons test (A-D) or two-way ANOVA with Tukey’s multiple comparisons test (E-H). Data from the vehicle and immunotherapy monotherapy groups are shown in Figure 1A–C, and Figure S1A. To assess the effect of P2RY2 targeting, these data are included here for comparison. A p-value greater than 0.05 indicates non-significance (n.s.), while $p < 0.05$ is denoted as ∗, and $p < 0.0001$ is denoted as ∗∗.

### Figure S6

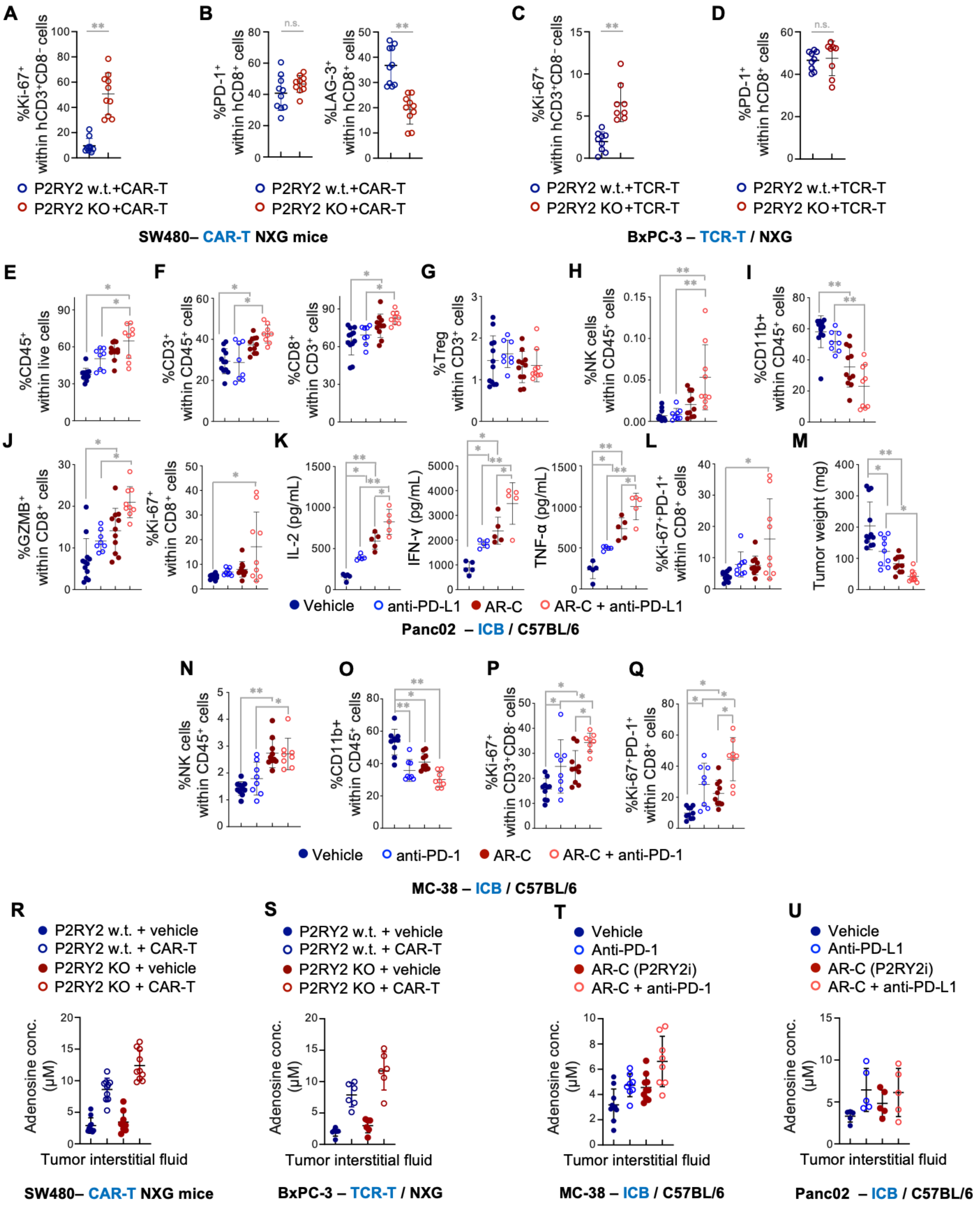

A
B
C
D
P2RY2 w.t.+CAR-T
P2RY2 KO+CAR-T
SW480– CAR-T NXG mice
P2RY2 w.t.+TCR-T
P2RY2 KO+TCR-T
BxPC-3 – TCR-T / NXG
E
F
G
H
I
J
K
L
M
Vehicle
anti-PD-L1
AR-C
AR-C + anti-PD-L1
Panc02 – ICB / C57BL/6
N
O
P
Q
Vehicle
anti-PD-1
AR-C
AR-C + anti-PD-1
MC-38 – ICB / C57BL/6
R
S
T
U
P2RY2 w.t. + vehicle
P2RY2 w.t. + CAR-T
P2RY2 KO + vehicle
P2RY2 KO + CAR-T
Vehicle
Anti-PD-1
AR-C (P2RY2i)
AR-C + anti-PD-1
Anti-PD-L1
AR-C + anti-PD-L1
Adenosine conc. (µM)
Tumor interstitial fluid
SW480– CAR-T NXG mice
BxPC-3 – TCR-T / NXG
MC-38 – ICB / C57BL/6
Panc02 – ICB / C57BL/6

**Figure S6. Targeting P2RY2 remodels the TME and enhances anti-tumor T cell responses, related to Figure 4.**

**(A, B)** As described in Fig. 3A, P2RY2-proficient (P2RY2 w.t.) and P2RY2-deficient (P2RY2 KO) human CRC (SW480) tumors were treated with human CEA CAR-T cells in immunodeficient NXG mice. Tumors were harvested at the endpoint.

(A) P2RY2-deficient tumors show increased proliferation of $CD3^+CD8^-$ CAR-T cells in the TME. Single-cell tumor suspensions were analyzed by flow cytometry to quantify Ki-$67^+$ proliferating cells within the human $CD3^+CD8^-$ T cell population in SW480 tumors.

(B)P2RY2 deficiency does not promote CAR-T cell exhaustion in the TME. The percentages of PD-$1^+$ and LAG-$3^+$ human $CD8^+$ CAR-T cells in SW480 tumors were assessed by flow cytometry.

(**C, D**) As described in Fig. 3B, P2RY2-proficient (P2RY2 w.t.) and P2RY2-deficient (P2RY2 KO) human PDAC (BxPC-3) tumors were treated with human TCR-T cells in NXG mice. Tumors were harvested at the end point.

(C) P2RY2-deficient tumors show enhanced proliferation of $CD3^+CD8^-$ TCR-T cells in the TME. Ki-$67^+$ proliferating human $CD3^+CD8^-$ T cells in BxPC-3 tumors were quantified by flow cytometry.

(D) The percentage of PD-$1^+$ human $CD8^+$ TCR-T cells in BxPC-3 tumors was assessed by flow cytometry.

(**E-M**) As described in Fig. 3D, immunocompetent C57BL/6 mice bearing syngeneic mouse PDAC (Panc02) tumors were treated with one of the following conditions: Vehicle control (Vehicle); PD-L1 blocking antibody (anti-PD-L1); small-molecule P2RY2 antagonist (AR-C); Combination therapy (AR-C + anti-PD-L1). Tumors were harvested at the endpoint.

(E, F) P2RY2 inhibition increases the infiltration of $CD45^+$ immune cells (E) and enhances the proportion of $CD3^+$ and $CD8^+$ T cells within the $CD45^+$ immune cell population (F) in the TME. The percentage of $CD45^+$ immune cells within all live cells, as well as $CD3^+$ T cells within $CD45^+$ immune cells and $CD8^+$ T cells within the $CD3^+$ population, were assessed by flow cytometry in tumor dissociates.

(G) P2RY2 inhibition does not significantly alter intratumoral Treg abundance. The percentage of $CD4^+Foxp3^+$ Tregs within $CD3^+$ T cells in Panc02 tumors was assessed by flow cytometry.

(H) Combined P2RY2 inhibition and ICB increases NK cell abundance in the TME. The percentage of NK ($CD3^-$ $CD11b^-$ NK1.$1^+$) cells within $CD45^+$ immune cells in Pac02 tumors was quantified by flow cytometry.

(I) P2RY2 inhibition reduces myeloid cell abundance in the immune infiltrates. The percentage of $CD11b^+$ myeloid cells within $CD45^+$ immune cells in Panc02 tumors was quantified by flow cytometry.

(J) P2RY2 inhibition enhances cytotoxicity and proliferation of $CD8^+$ T cells in the TME. The percentages of granzyme B $(GZMB)^+$ and Ki-$67^+$ cells within the $CD8^+$ T cell population in Panc02 tumors were quantified by flow cytometry.

(K) P2RY2 inhibition increases effector cytokine accumulation in the TME. IL-2, IFN-γ, and TNF-α concentrations in the interstitial fluid of Panc02 tumors were measured using LEGENDplex™ Assays.

(L) The percentage of Ki-$67^+$PD-$1^+$ cells within the $CD8^+$ T cell population in Panc02 tumors was quantified by flow cytometry.

(M) P2RY2 inhibition reduces tumor burden and enhances the efficacy of ICB. Tumor weight of Panc02 tumors was measured at the experimental endpoint.

(**N-Q**) As described in Fig. 3C, immunocompetent C57BL/6 mice bearing syngeneic mouse CRC (MC-38) tumors were treated with one of the following conditions: Vehicle control (Vehicle); PD-1 blocking antibody (anti-PD-1); small-molecule P2RY2 antagonist (AR-C); Combination therapy (AR-C + anti-PD-1). Tumors were harvested at the endpoint.

(N) P2RY2 inhibition increases the abundance of NK cells in the TME of MC-38 tumors. The percentage of NK ($CD3^{-}$ $CD11b^{-}$ $NK1.1^{+}$) cells within $CD45^{+}$ immune cells in tumor dissociates was assessed by flow cytometry.
(O) P2RY2 inhibition reduces myeloid cell abundance in the immune infiltrates. The percentage of $CD11b^{+}$ myeloid cells within $CD45^{+}$ immune cells in MC-38 tumors was quantified by flow cytometry.
(P) P2RY2 inhibition enhances the proliferation of $CD3^{+}CD8^{-}$ T cells in the TME. The percentage of Ki-$67^{+}$ proliferating cells within $CD3^{+}CD8^{-}$ T cells in MC-38 tumors was quantified by flow cytometry.
(Q) The percentage of Ki-$67^{+}$PD-$1^{+}$ cells within $CD8^{+}$ T cells in MC-38 tumors was quantified by flow cytometry.

(**R**-**U**) Targeting P2RY2 does not reduce intratumoral adenosine levels. Adenosine concentrations in the interstitial fluid of tumors from the SW480-CAR-T model (**R**), BxPC-3-TCR-T model (**S**), MC-38-anti-PD-1 model (**T**) and Panc02-anti-PD-L1 model (**U**), as described in Fig. 3, A to D, were measured using the Adenosine Assay Kit.

Data are presented as mean ± standard deviation of biological replicates. For panels (A, B, R), the sample sizes are: P2RY2 wild-type (w.t.) + vehicle (n=10), P2RY2 knockout (KO) + vehicle (n=9), P2RY2 w.t. + CAR-T (n=10), and P2RY2 KO + CAR-T (n=10). For panels (C, D), the sample sizes are: P2RY2 wild-type (w.t.) + vehicle (n=9), P2RY2 knockout (KO) + vehicle (n=8), P2RY2 w.t. + TCR-T (n=9), and P2RY2 KO + TCR-T (n=8). For panels (E-J, L, M) are: vehicle (n=12), anti-PD-L1 (n=8), AR-C (n=11), and AR-C + anti-PD-L1 (n=9). For panel (K, U) are: vehicle (n=5), anti-PD-L1 (n=5), AR-C (n=5), and AR-C + anti-PD-L1 (n=5). For panels (N-Q, T), the sample sizes are: vehicle (n=10), anti-PD-1 (n=8), AR-C (n=10), and AR-C + anti-PD-1 (n=8). For panel (S), the sample sizes are: P2RY2 wild-type (w.t.) + vehicle (n=6), P2RY2 knockout (KO) + vehicle (n=6), P2RY2 w.t. + TCR-T (n=6), and P2RY2 KO + TCR-T (n=6). Statistical significance was determined using unpaired two-tailed Student's t-test (A-D), or one-way ANOVA with Tukey's multiple comparisons test (E-Q). A p-value greater than 0.05 indicates non-significance (n.s.), while $p < 0.05$ is denoted as $*$, and $p < 0.0001$ is denoted as $**$.

**Table S1. The sequences of sgRNAs targeting P2X and P2Y purinergic receptors, related to Figure 1L and M.**

| Target | sgRNA sequences |
|---|---|
| sgP2RX1 #1 | AAGGGCTACCAGACCTCGAG |
| sgP2RX1 #2 | GAAGGTCAGAGTGTTACCGC |
| sgP2RX2 #1 | CACGTCCGAGCACAAAGTGT |
| sgP2RX2 #2 | CAGCCAATTTCTGGGTACGA |
| sgP2RX3 #1 | CTCACTTTAGGGATCCTCAC |
| sgP2RX3 #2 | CTTCACCTTGGTTACCACCG |
| sgP2RX4 #1 | CGGGTCTGTCAAGACGTGTG |
| sgP2RX4 #2 | GGCATCTGATTTACACACAG |
| sgP2RX5 #1 | AGACGGCTAAAAGAATAGTG |
| sgP2RX5 #2 | CTGGGCTGGAATGACGTAGT |
| sgP2RX6 #1 | GGATCGTGGTCTATGTGGTA |
| sgP2RX6 #2 | GGCAGTGCTTAAAATAGGTG |
| sgP2RX7 #1 | CAAAGGGAAGGTGTAGTCTG |
| sgP2RX7 #2 | CAGAAGGTACCTTTGCTCTG |
| sgP2RY1 #1 | ACTTCGCAGGTACTCGTCTG |
| sgP2RY1 #2 | AGCCCAGAATCAGCACCAAG |
| sgP2RY2 #1 | CACCACATATATGTTCCACC |
| sgP2RY2 #2 | CGTAACCTGCCACGACACCT |
| sgP2RY4 #1 | AAGACAACTGCATAGCTCAC |
| sgP2RY4 #2 | CCACTTCGGGCACTACGCTG |
| sgP2RY6 #1 | CATGCCATAGGGCATATAGT |
| sgP2RY6 #2 | GCATAGTTGTAGATGAGCAG |
| sgP2RY11 #1 | ACGCAGCCCAGGCATGACTG |
| sgP2RY11 #2 | AGGCCTGCATCAAGTGTCTG |
| sgP2RY12 #1 | CAGAGACTACAAAATCACCC |
| sgP2RY12 #2 | CCAACCCCAAAAATCTCTTG |
| sgP2RY13 #1 | AAATACGATCTTGAGCAACA |
| sgP2RY13 #2 | ATGAACACCACAGTGATGCA |
| sgP2RY14 #1 | AAAGTGAACTGGGACGGAAG |
| sgP2RY14 #2 | ATCCTGACACTCCATTGAGT |
